# Phylogenomics supports island contribution to metapopulation dynamics in a predominantly continental bird species

**DOI:** 10.1101/2023.09.27.559751

**Authors:** Daisuke Aoki, Masayuki Senzaki, Haruko Ando, Yoshiya Odaya, Wieland Heim, Munehiro Kitazawa, Wulf Tom, Daronja Trense, Marc Bastardot, Atsunori Fukuda, Masao Takahashi, Natsuko Kondo

## Abstract

**Aim:** Islands have recently been recognized as potential sources of biodiversity, challenging the traditional view that their small population sizes and low genetic diversity limit such roles. This raises the question of how insular genetic variation becomes incorporated into continental populations, contrary to expectations of unidirectional colonization. Here, we investigate whether and how island-derived genetic variation has influenced a continental population through population establishment and gene flow in a bird species where frequent trans-ocean dispersal is expected.

**Location:** Continental East Asia (Russian Far East), Japanese Archipelago

**Taxon:** Swinhoe’s Rail (Coturnicops exquisitus)

**Materials and Methods:** We apply integrative phylogenomics to reconstruct the spatiotemporal history of the species. Colonization sequences and gene flow are inferred by comparing four different phylogenetic reconstruction methods, using mitochondrial sequences obtained by Sanger sequencing and genome-wide data obtained by genotyping by sequencing (MIG-seq). We assess a history of colonization and gene flow based on summary statistics, demographic trajectory inference by Stairway Plot2, demographic modeling by fastsimcoal2, and species distribution modeling.

**Results:** Analyses collectively supported asymmetric gene flow from the island to the continental population, following divergence around the Middle Pleistocene. Post-divergence, the island maintained a large and stable population, while the continental population underwent a severe bottleneck, suggesting a significant evolutionary role of the island for the continental population. Additionally, evidence of recent re-establishment of the island by continental individuals indicates dynamic exchange and persistence within a continent-island metapopulation.

**Main conclusions:** The maintenance of insular genetic variation within a dynamic continent-island metapopulation may have enabled the island to act as a genetic and demographic reservoir for the continental population. Thus, continent-island metapopulation dynamics may be a key evolutionary pathway through which island populations contribute to continental genetic diversity.

## Introduction

Island biological diversity contributes disproportionately to global biodiversity (Whittaker and Fernández-Palacios 2007). Islands, comprising 3.5% of the terrestrial land, contribute nearly 20% of global species richness and more than 60% of terrestrial extinctions (Whittaker et al. 2017). Traditionally, islands were assumed to sustain smaller population sizes than continents due to their small and isolated landmasses (Leroy et al. 2021). Small populations are typically subjected to strong genetic drift, resulting in reduced genetic variation (Leroy et al. 2021; Cerca et al. 2023), which likely increases extinction risks (Frankham 2005) and leads to “evolutionary dead ends” (Vamosi et al. 2014). Consequently, islands have often been viewed as sinks where lineages evolve independently and do not contribute to global diversification (Bellemain and Ricklefs 2008).

Recently, evidence has emerged showing that islands can also serve as sources of continental colonization (Whittaker et al. 2017). Macroevolutionary (i.e., species-level) studies revealed numerous cases of continental species or lineages nested within island radiations–a pattern termed as “reverse colonization” (Bellemain and Ricklefs 2008). A review found that over one-third of the phylogenies across diverse taxa in various continent-island systems exhibited such patterns (Bellemain and Ricklefs 2008). Reverse colonization can also trigger mainland diversification. For example, the passerine radiation–now nearly 60% of global bird species–originated from colonization out of New Guinea (Jønsson and Holt 2015).

Although reverse colonization operates at the species level (i.e., a new continental species arises from an ancestral island species), its microevolutionary (i.e., population-level) mechanisms remain poorly understood. Speciation requires several population-level processes, including the establishment and persistence of populations–processes influenced by spatiotemporal dynamics of migration between the ancestral and derived populations (Harvey et al. 2019). For instance, depending on divergence levels, gene flow from ancestral populations may lead to genetic or demographic rescue (Hufbauer et al. 2015), introduce novel genetic variation (Cerca et al. 2023), or cause incompatibilities (Bank et al. 2012), thereby facilitating or impeding divergence. Given the distinct population genetic processes and environments experienced by island and continental populations (Barton 1996; Weigelt et al. 2013), these population-level processes are likely important at the initial stage of reverse colonization. To our knowledge, there has been only one study that modeled the population genetic process of continental establishment by insular organisms (Patiño et al. 2015). This study, however, used bryophytes, whose genetic variation may not be sensitive to demographic changes due to their ability for asexual reproduction (Ellegren and Galtier 2016), leaving why many sexually reproductive animals exhibit reverse colonization uncovered. Furthermore, gene flow in continent-island systems has mostly been studied in the context of divergence-with-gene-flow– how island populations diverge despite gene flow (Warren et al. 2015)–with a focus on genomic regions resisting homogenization (Feder et al. 2012). As a result, studies have been biased towards the effects of continent-derived gene flow, with limited exploration of migration dynamics and their demographic consequences in continent-island systems as a whole.

Phylogeography is a powerful framework to examine population establishment and gene flow directionality in continent-island systems, as it infers historical processes shaping present genetic patterns (Lomolino et al. 2017). When integrated with demographic reconstruction and species distribution modeling (SDM), it becomes especially effective in reconstructing spatiotemporally explicit histories. Phylogenetic trees provide patterns that emerge as a result of demographic processes, while the processes themselves remain unmodeled (Harvey et al. 2019). In contrast, demographic reconstruction using high-throughput sequencing explicitly models these processes (Nadachowska-Brzyska et al. 2022). SDMs estimate the timing and extent of historical range shifts (Harvey et al. 2019). Therefore, these methods aid inference on the direction, intensity, and timing of population establishment or gene flow.

Here, we apply integrative phylogeography to examine the direction and demographic consequences of population establishment and gene flow in a continent-island system. We selected Swinhoe’s Rail (*Coturnicops exquisitus*), the smallest Asian rail, as the ideal model. Demographic processes shape the present genetic variation of birds more strongly than in many other taxa (Ellegren and Galtier 2016), and therefore, they are expected to underpin important demographic processes in continent-island systems. Rails are known for repeated transoceanic dispersals, as inferred from phylogenetic reconstruction at the species level (Garcia-R and Matzke 2021). Swinhoe’s Rail mainly breeds in continental East Asia (Heim et al. 2019), while a limited number of island populations have recently been discovered (Senzaki et al. 2020). Therefore, we expect that Swinhoe’s Rail is prone to frequent colonization events across the sea, even at the population level, and it is ideal to examine how a currently small island population has historically contributed to the continental population establishment, persistence, and evolution.

In this study, we reconstruct the phylogeographic history of Swinhoe’s Rail to test whether island genetic variation has contributed to continental populations via population establishment and/or gene flow. We infer the direction of establishment by examining 1) whether the insular population forms an early-branching clade with nested continental populations, 2) the demographic stability of each population, and 3) habitat suitability at the time of divergence. We assume that derived populations arose through bottlenecks from stable ancestral ones (e.g., Yeung et al. 2011). We determine the direction of gene flow by 1) analyzing phylogenetic signals of gene flow, 2) estimating gene flow by demographic modeling, and 3) predicting historical range shifts that could enable migration by the SDM. Finally, we discuss how these events impacted population size and genetic variation on the continent.

## Materials and Methods

### Field sampling

Swinhoe’s Rail is a poorly studied species. Previously known breeding ranges were limited to the continent (Fig. 1a), while evidence of breeding is now absent from most of the region (Heim et al. 2019). The currently recognized breeding areas were re-discovered recently, including the Amur region (Heim et al. 2019) and Lake Baikal (Anisimova et al. 2019) in Russia, as well as parts of Japan, namely Aomori of Honshu (Miya et al. 2005) and Hokkaido (Senzaki et al. 2020) (Fig. 1a). Despite the species’ cryptic behavior and preference for inaccessible wetlands, we successfully developed sampling strategies (Heim et al. 2019; Senzaki et al. 2020) and collected feather, buccal swab, or blood samples covering the entire extent of the currently known breeding range: Amur (n = 6) and Baikal (n = 1) for continental Russia, Kushiro (n = 12) and Tomakomai (n = 8) for Hokkaido, and Aomori (n = 4) for Honshu during their breeding season (Fig. 1a, Table S1). We also collected samples of wintering birds in the Kanto region of Honshu (n = 37) and used a genetic sample from a migratory bird in Ishikari, Hokkaido (Supplementary Methods), to assess migratory connectivity and historical range shifts. Additionally, we retrieved mitochondrial data from GenBank for phylogenetic analyses (Table S1).

**Figure 1.**
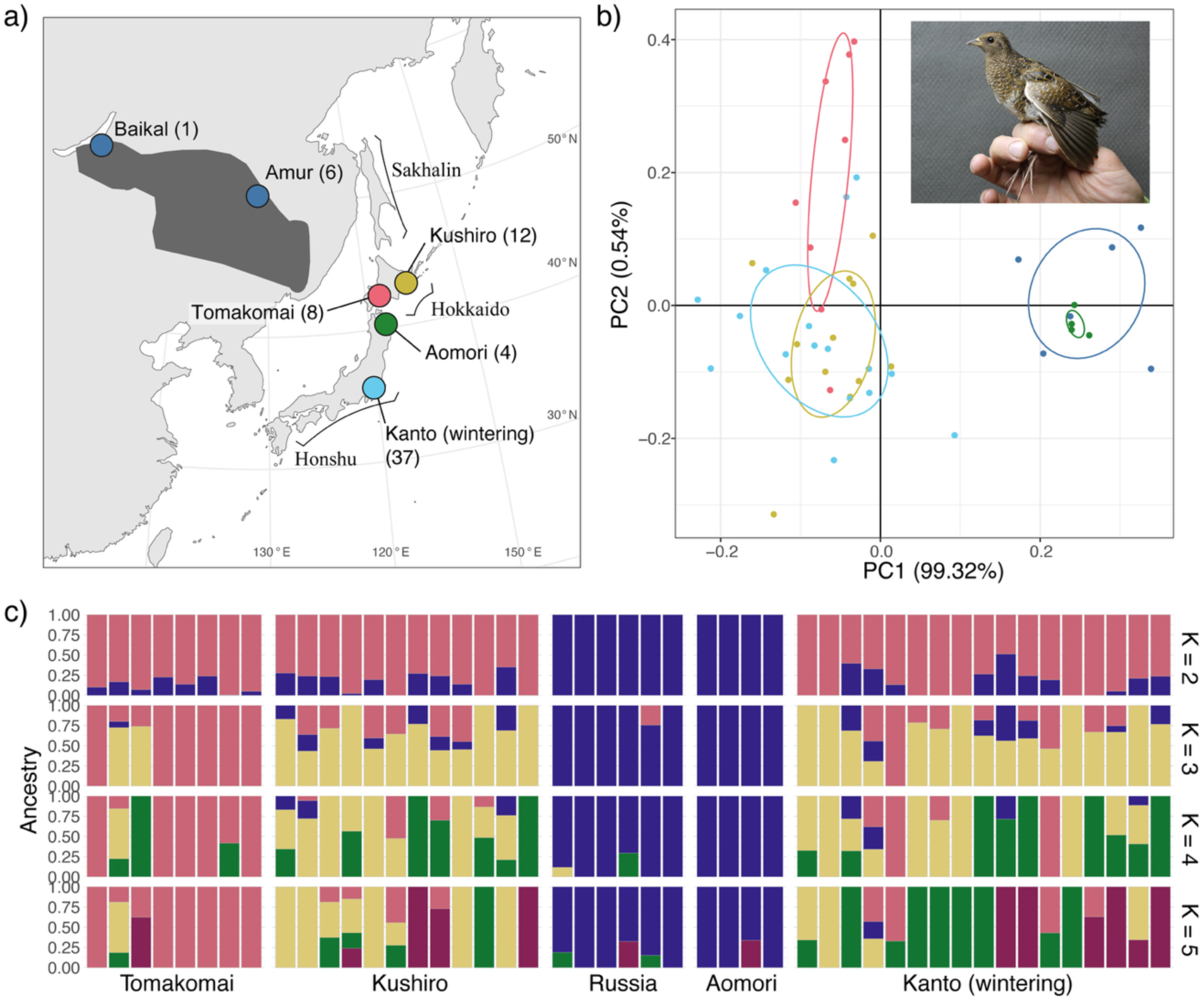
A map of sampling localities of Swinhoe’s rails (a), and their population genetic structure inferred by (b) a principal component analysis (PCA) and (c) an ADMIXTURE analysis conducted using genotype likelihoods of 1,147 single nucleotide polymorphisms (SNPs). (a, b) The colors of dots and inertia ellipses represent different regional populations. Dark-grayed regions are the expert range map of this species in BirdLife International and Handbook of the Birds of the World (2024), which also includes previously known ranges. (c) Genetic assignment of individuals to each number of genetic clusters (K = 2 to 5) is shown. Colors do not correspond to the sampling localities. Lambert Azimuthal Equal Area projection was used for the map. The inset photo by Marc Bastardot.

### Laboratory procedures & Bioinformatics

The whole genomic DNA was extracted from each tissue and used in two experiments: Sanger sequencing of the partial mitochondrial Cytochrome *b* (*Cytb*) region for 49 samples, and multiplexed inter-simple sequence repeats genotyping by sequencing (MIG-seq; Suyama and Matsuki 2015) for 52 samples (Table S1). MIG-seq, a polymerase chain reaction-based genotyping by sequencing approach, is effective for low-quality DNA at the cost of reduced genome coverage and is appropriate for swab and dried feather samples. It yielded 4,834,210 read pairs (9,668,420 reads), which were trimmed and aligned to a reference genome of a closely related species, *Laterallus jamaicensis coturniculus* (WGS Accession ID, JAKCOX01, Table S1). Three individuals with over 30% missing sites in ANGSD pre-runs (one from Amur and two from Kanto) were excluded (Fig. S1–S4), yielding a final dataset of 7,244,578 paired and 37,186 unpaired reads for 49 samples. See Supplementary Methods for parameter settings.

We accounted for genotype uncertainty by calculating genotype likelihoods (GLs) with ANGSD (Korneliussen et al. 2014) (see Supplementary Methods, Fig. S1–S2). GLs were first calculated separately for five regional populations, using half the sample size of each population as ‘-minInd’ and 2 as ‘-setMinDepthInd’, retaining sites with depth ≥2 in over half the samples. For downstream analyses, GLs were recalculated on intersecting sites among sample subsets using different filtering parameters, yielding varying site counts (Table 1, S2, Fig. S4). For analyses of variant sites, we set 1e-06 for ‘-SNP-pval’ and 0.05 for ‘-minMaf’. For SNAPP analyses, genotypes were called using ‘-doGeno’ with a posterior cutoff of 0.9. Populations were defined either by five sampling regions (Fig. 1a) or by two genetically supported breeding groups–“island” (Kushiro + Tomakomai) and “continental” (Russia + Aomori) (Table 1). This grouping was robustly supported by multiple analyses (Fig. 1b–c, 2b, S5–S10). The ‘SAMtools’ model was used in the entire GL calculations. Linkage disequilibrium (LD) was assessed and pruned using ngsLD (Fox et al. 2019) for some analyses (Table 1). Demographic inference used folded site frequency spectra (SFS). Full workflows and parameters are provided in Supplementary Methods and Fig. S4.

**Table 1.** Dataset and the number of sites used in each analysis.

| Analyses | Linkage | Site type | Outgroup | Populations | # Sites | # Samples | Notes |
| --- | --- | --- | --- | --- | --- | --- | --- |
| **PCA, ADMIXTURE** | unlinked | V | Non | 4 breeding<sup>1</sup>,<br>1 wintering<sup>2</sup> | 1,147 | 47 | - |
| **NeighborNet (ngsDist)** | linked | V | Laja/Atro | 4 breeding | 1,958 | 32 |  |
| **RAXML, D-statistic** | linked | V/I | Laja | 4 breeding | 108,095 | 31 | - |
| **TreeMix** | linked | V | Laja | 4 breeding | 1,669<br>(683) | 31 | Population-specific private alleles were removed by glactools. The number of sites inside the bracket is for the individual-level tree. |
| **SNAPP** | unlinked | V | Laja | 4 breeding | 206 | 16 | The top three/four samples with the lowest missing allele proportions in each population were selected. Excess heterozygous sites were removed by BCFtools |
| **Diversity indices, FSC2, SWP2** | linked | V/I | Non | 2 genetic<sup>3</sup> | 130,368 | 30 | - |

Abbreviation: PCA, principal component analysis; FSC2, fastsimcoal2; SWP2, Stairway Plot 2;
V, variants; V/I, both variants and invariants; Laja, *Laterallus jamaicensis*; Atro, *Atlantisia*
*rogersi*; <sup>1</sup> Tomakomai, Kushiro, Russia, and Aomori; <sup>2</sup> Kanto; <sup>3</sup> Island, and Continental.

### Genetic structure

We examined global genetic structure via principal component analysis (PCA) using PCAngsd (Meisner and Albrechtsen 2018), ADMIXTURE analysis with NGSadmix (Skotte et al. 2013) with K=1–6, NeighborNet (Bryant and Moulton 2004) based on pairwise genetic distance from ngsDist (Vieira et al. 2015), and the D-statistic using ‘doabbababa2’ in ANGSD (Soraggi et al. 2018) (see Supplementary Methods for details). Two Kanto samples, closely related to one Tomakomai and one Kushiro sample, showing kinship coefficients (KING, Waples et al. 2019) > 0.088 based on estimates from ngsRelate (Hanghøj et al. 2019), were excluded to meet the assumptions of these analyses. We included two American *Laterallus* group species (*Laterallus jamaicensis,* Accession SRX14572910; *Atlantisia rogersi,* Accession SRX6608087) as outgroups in ngsDist and supplementary PCA/ADMIXTURE analyses (Fig. S5–S7). After confirming their monophyly, only *L. jamaicensis* was used as an outgroup for phylogenetic reconstruction.

### Inference of phylogenetic relationship

Phylogenetic trees from different datasets and methods can reflect gene flow and incomplete lineage sorting (ILS) differently (Twyford and Ennos 2012). Thus, we inferred a phylogenetic relationship through four approaches: a mitochondrial phylogenetic tree (mtDNA tree) using *Cytb* data, a maximum likelihood tree (RAxML tree), a graph-based population tree based on allele frequency (TreeMix), and a coalescent-based population tree based on single nucleotide polymorphisms (SNPs) (SNAPP tree) using MIG-seq genome-wide nuclear data.

#### mtDNA tree

The time-calibrated Bayesian inference was performed on 772 bps of partial *Cytb,* a widely used marker suitable for avian phylogeography (e.g., Weir and Schluter 2004), using an uncorrelated lognormal relaxed clock model and coalescent exponential population prior in BEAST v.2.7.3 (Bouckaert et al. 2019). Samples included 12 haplotypes from 54 Swinhoe’s rails and three outgroups (two *C. noveboracensis*, one *L. jamaicensis*; Table S1). The standard mitochondrial molecular clock rate for birds, 1.05%/lineage/Myr (Weir and Schluter 2008), was applied to estimate divergence times. Markov chain Monte Carlo (MCMC) was run for 2 million generations, sampling every 1,000. Convergence was checked by following Aoki et al. (2018), and a maximum clade credibility tree with mean heights was drawn after discarding the first 10% as burn-in. We confirmed the inferred tree by a haplotype network. See Supplementary Methods for settings.

#### RAxML tree

A maximum likelihood tree was reconstructed from both variant and invariant sites using RAxML-ng (Kozlov et al. 2019) under the ‘GTGTR4+FO+G’ model. Genotype probabilities (GPs) calculated in a VCF file output of ANGSD were used to prepare a CATG input file. For missing sites, a low, uninformative probability of 0.01 was assigned. With ‘--prob-msa’ enabled, the tree was reconstructed considering genotype uncertainty. Standard non-parametric bootstrapping with 100 replicates was simultaneously conducted.

#### TreeMix

We used TreeMix (Pickrell and Pritchard 2012) to infer a population-level tree incorporating gene flow and drift. Allele frequencies were estimated based on GLs using glactools (Renaud 2017), and an ACF file was converted to TreeMix input by the ‘acf2treemix’ program with a ‘--noprivate’ option, to exclude population-specific alleles. Migration edges ranging from zero to five were tested across ten replicates with a block size of 20 (based on the LD decay; Supplementary Methods) for jackknife sampling. The best number of migration edges scoring the highest *Δm* score was determined by an R package ‘OptM’ v. 0.1.6 (Fitak 2021), selecting the run with the highest model likelihood as the best-fit model. We repeated the same analyses without grouping (i.e., individual-level tree) to confirm that inferred gene flow reflects actual migration rather than population grouping artifacts.

#### SNAPP tree

SNAPP infers a population tree based on a coalescent model using unlinked SNPs without missing data (Bryant et al. 2012). We selected three or four samples scoring the lowest missing proportion from each breeding population (three from Aomori due to the small sample size), whose missing proportions ranged between 2% and 12%. Unlinked hard-called SNPs without missing data were obtained from these samples by ANGSD. We estimated tree topology while simultaneously estimating divergence times (Stange et al. 2018). We calibrated the split between Swinhoe’s Rail and *L. jamaicensis* using two log-normal priors (lognormal(5.11, 0.09) and lognormal(8.95, 0.0225) in million years ago [Mya]), which corresponded to its divergence time estimates by our clock-calibrated mtDNA tree (Fig. 2a) and fossil-calibrated mtDNA tree by Garcia-R et al. (2020), respectively. We ran 500,000 MCMC iterations to get posterior trees and analyzed the posterior trees using ‘TreeSetAnalyser’ to derive a 95% credible set.

**Figure 2.**
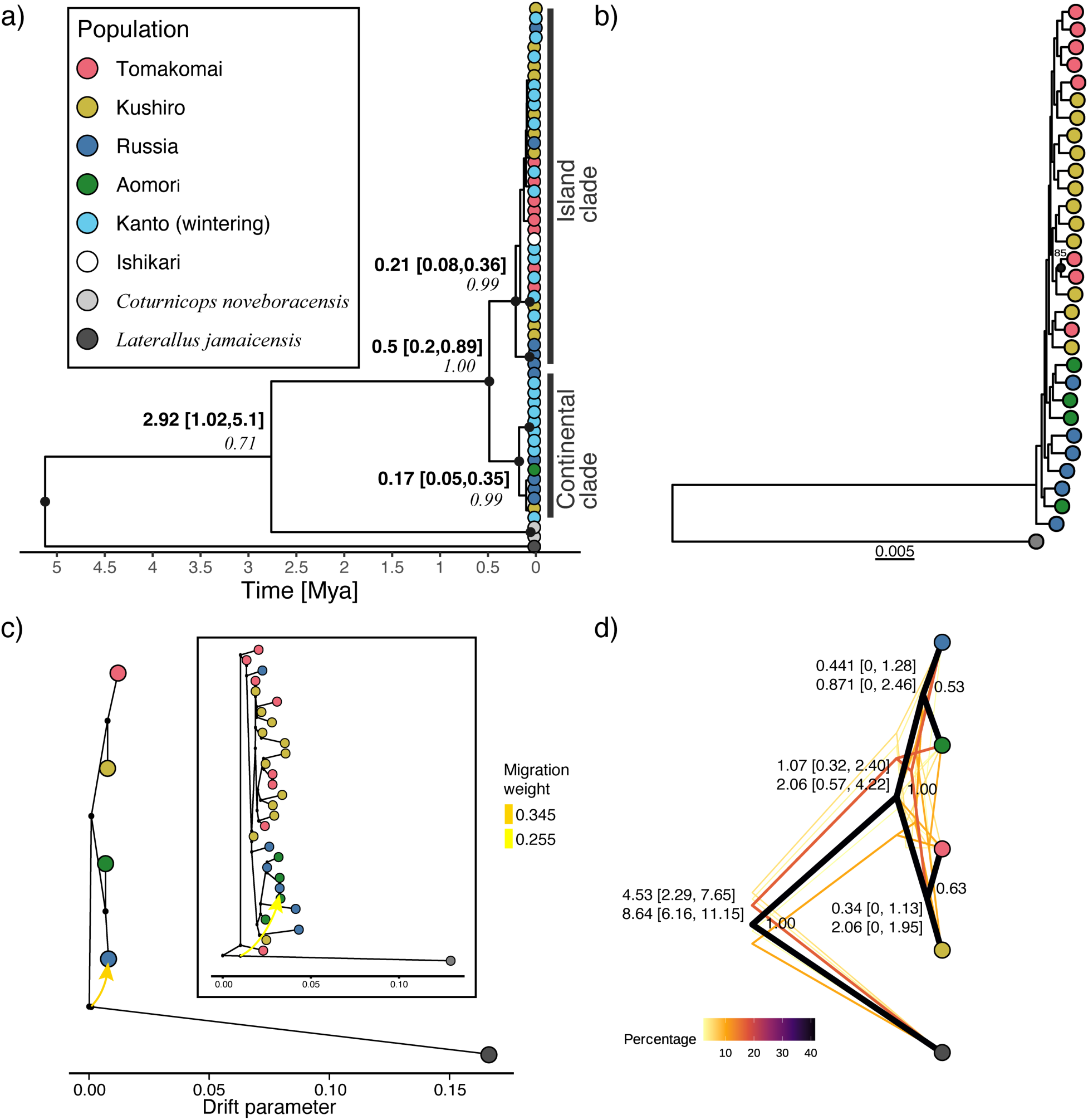
Phylogenetic trees of Swinhoe’s Rail reconstructed using different methods and genetic markers. (a) A mitochondrial phylogenetic tree (mtDNA tree) using 772 bp of the *Cytb* gene. Posterior probabilities >0.9 were indicated by black dots on nodes. Numbers at the nodes indicate median divergence time estimates with 95% highest posterior density within brackets (bold) and posterior probabilities (italic). (b) A maximum likelihood (ML) tree (RAxML tree) was reconstructed using 108,095 sites including both variant and invariant sites. Only nodes with bootstrap values >70 are annotated with black circles. (c) A population-level tree reconstructed by TreeMix using the allele frequency of 1,669 polymorphic loci, excluding population-specific private alleles. Branch lengths indicate drift parameters, the strength of drift specific to each branch. The inset indicates an individual-level tree reconstructed using non-private 683 polymorphic loci. The colored arrows indicate the estimated gene flow. (d) Posterior population-level trees reconstructed by SNAPP analyses using 206 single nucleotide polymorphisms (SNPs) with no missing data. Colors of trees indicate percentages of each posterior tree in the credible set obtained by the ‘TreeSetAnalyser’ program. Values at each node indicate their divergence time estimates with credible intervals within brackets. Two different divergence time estimates were obtained for different calibration constraints. Values beneath each node indicate posterior probabilities.

### Demographic inferences

We examined the historical demography of the continental and island populations via summary statistics, demographic trajectory inferences (Stairway Plot 2, SWP2), and parameter estimations in demographic models (fastsimcoal2, FSC2). These methods are complementary to each other because SWP2 is a nonparametric model, flexibly inferring complex effective population size (*N_e_*) trajectories yet without modeling gene flow, while FSC2 models both gene flow and *N_e_* changes but in user-specified simpler models (Nadachowska-Brzyska et al. 2022). We also estimated the contemporary *N_e_* of the island population by converting the estimated census population size (*N_c_*) in a plain of Hokkaido in the early nineteenth century (Kitazawa et al. 2022), using *N_c_*:*N_e_* = 1:0.44, a mean ratio for birds (Frankham 1995).

#### Summary statistics

We calculated Watterson’s theta θ_W_ (Watterson 1975) and Tajima’s *D* (Tajima 1983) per reference scaffold and population. Tajima’s *D* values around zero suggest demographic stability, and values below zero (especially <-1) indicate a strong ancient bottleneck (Gattepaille et al. 2013). Furthermore, the large and small variance of Tajima’s *D* among scaffolds reflects moderate bottleneck and population expansion, respectively (Gattepaille et al. 2013). Per-scaffold estimates of these indices were compared between populations using linear mixed models. We also estimated the fixation index (F_ST_) using two-dimensional SFS between four breeding populations. Details in Supplementary Methods.

#### SWP2

We inferred demographic trajectories of the continental and island populations using one-dimensional folded SFS in Stairway Plot 2 v. 2.1.1 (Liu and Fu 2020). We assumed a mutation rate of 1.91e-09 bp/year (Nam et al. 2010) and converted it to bp/generation by using a generation time of 1 year (a value is for its congener, *C. noveboracensis*; Alvo and Robert 1999). Default settings were used. The timing at which populations started to show diverging demographic trends was interpreted as a sign of population divergence.

#### FSC2

We tested five models differing in the presence and timing of migration matrices (Fig. 3c), using two-dimensional SFS. We estimated t_MIG_ at which both the continental and island populations experienced demographic changes and migration matrices may have switched, by fixing their divergence time at 0.5 Mya, which was roughly but collectively agreed by multiple approaches (Fig. 2a, 2d, 3f). Between the two populations, we modeled no migration throughout (“no migration”), migration only after (“recent migration”) or before (“ancient migration”) t_MIG_, and migration throughout at the same rates (“constant migration”) or different rates (“different migration”) across t_MIG_ (Fig. 3c). We conducted 100 replicates of 1 million coalescent simulations across 40–100 optimization ECM cycles for parameter estimates, in fastsimcoal2 v. 2.7.0.9 (Excoffier et al. 2021). The same mutation rate as the SWP2 analyses was used. A simulation with the smallest difference between ‘MaxEstLhood’ and ‘MaxObsLhood’ was selected from each model, and models with the ΔAIC (delta Akaike Information Criterion, the difference between the lowest AIC and an AIC in focus) <2 were considered best-fit. The ranges of parameter values for simulation are in Table S5.

**Figure 3.**
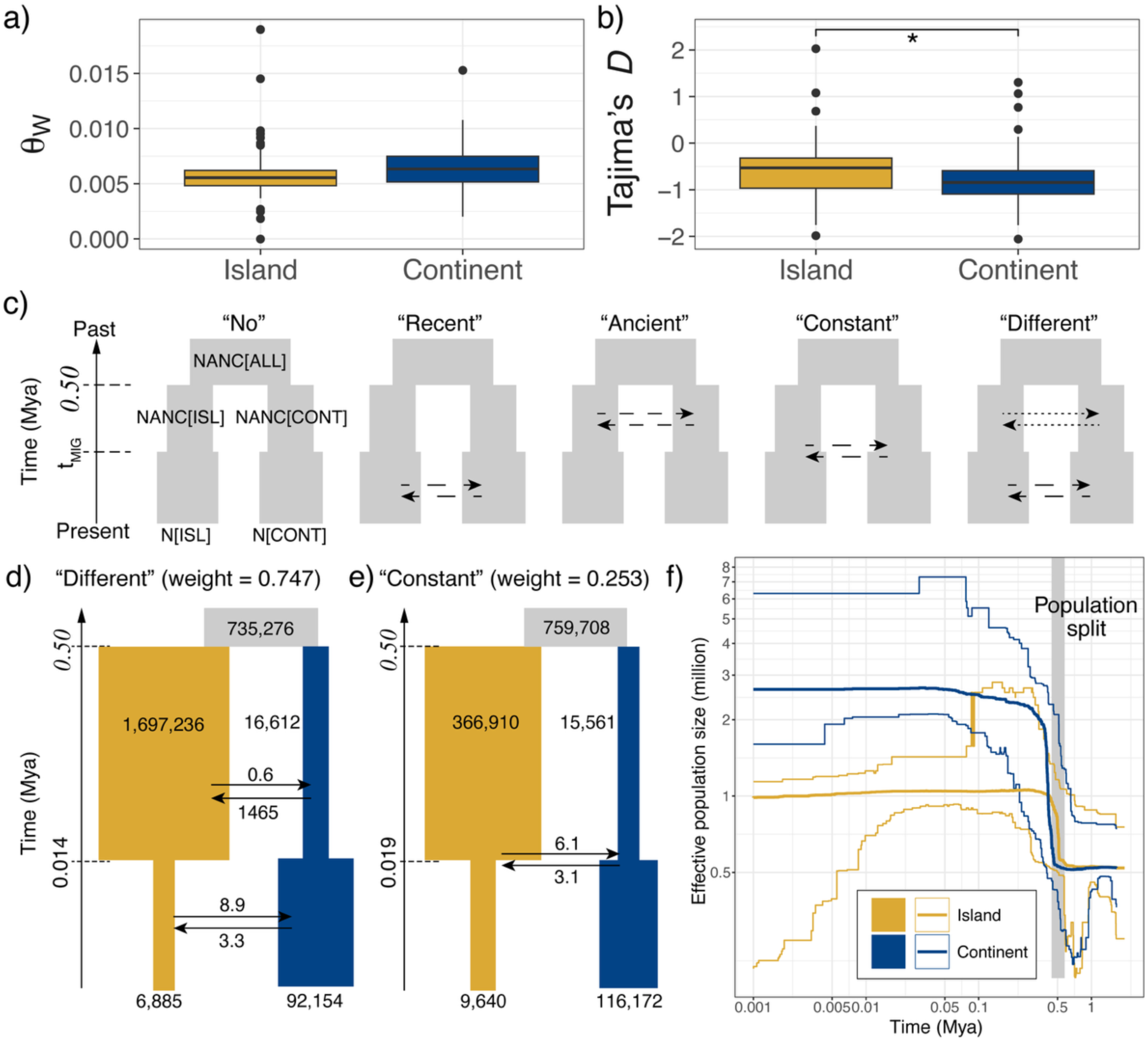
Demographic inference of the two genetic populations of Swinhoe’s Rail based on site frequency spectra, using (a, b) the summary statistics, (c–e) fastsimcoal2 (FSC2), and (f) Stairway Plot 2. Fixed values in each model are italicized. Effective population size and time are scaled by mutation rate (1.91e-09 bp/generation) and generation time (1 year). Effective population size shows the number of diploid individuals. (a, b) Comparison of scaffold-level statistics between the island and continental populations for (a) Watterson’s theta and (b) Tajima’s *D*. Statistically significant differences in the values of each index were annotated with asterisks (* *p* < 0.05). (c) Generalized schematic representation of the five models tested. Parameters with uppercase represent the effective population size we modeled. Hatched and dotted arrows represent different migration matrices within the model. (d, e) The two best-fit FSC2 models with estimated parameters. Numbers by or on branches and arrows represent effective population size and the number of migrants per year, respectively. The size of each box is not scaled to the corresponding population size. Divergence time is shown in million years. (f) Changes in effective population size (in millions) over time (in million years) are shown for the island and continental populations. Bold lines and thin ribbons around them show median and 95% confidence intervals, respectively. The gray range indicates the approximate period of population divergence.

### Species distribution modeling

SDMs were run using MaxEnt (Phillips et al. 2006) to model the species’ breeding extent. We compiled 48 breeding and 111 non-breeding records from previously published sources (Table S6), including both previously known and recently discovered sites (Fig. S20). After spatial thinning (one record per 0.5° grid cell), 33 breeding points were retained as an occurrence input (Fig. 4b). Background points were sampled to cover one-fourth of the total number of girds in a 1,000-km buffered convex hull around the full dataset (both breeding and non-breeding records), resulting in 2,104 points (Fig. S20). We extracted time slices of 22 environmental variables (Krapp et al. 2021) using an R package ‘pastclim’ v. 1.2.3 (Leonardi et al. 2023) and selected eight covariates with low collinearity, satisfying that any pair scores variance inflation factor <10. Twelve models with different parameter settings were tested via ‘ENMevaluate’ in an R package ENMeval v. 2.0.4 (Kass et al. 2021), and the best model was selected based on the lowest ΔAIC for a small sample size (ΔAICc). The best-fit model was projected across time slices from the present and 0.5 Mya at 1,000-year intervals (Fig. 4c). We compared the historical changes in the potential habitat of this species on the island and the continent (Fig. S23), by calculating three indices, habitat suitability (sum of values of the predicted presence grids), availability (the total number of predicted presence grids), and quality (suitability divided by availability). We also calculated centroid distances (weighted by suitability) between island and continental habitats to track historical changes in the level of geographic isolation. Effects of negative multivariate environmental similarity surface (MESS) were controlled for these calculations. See Supplementary Methods for details.

**Figure 4.**
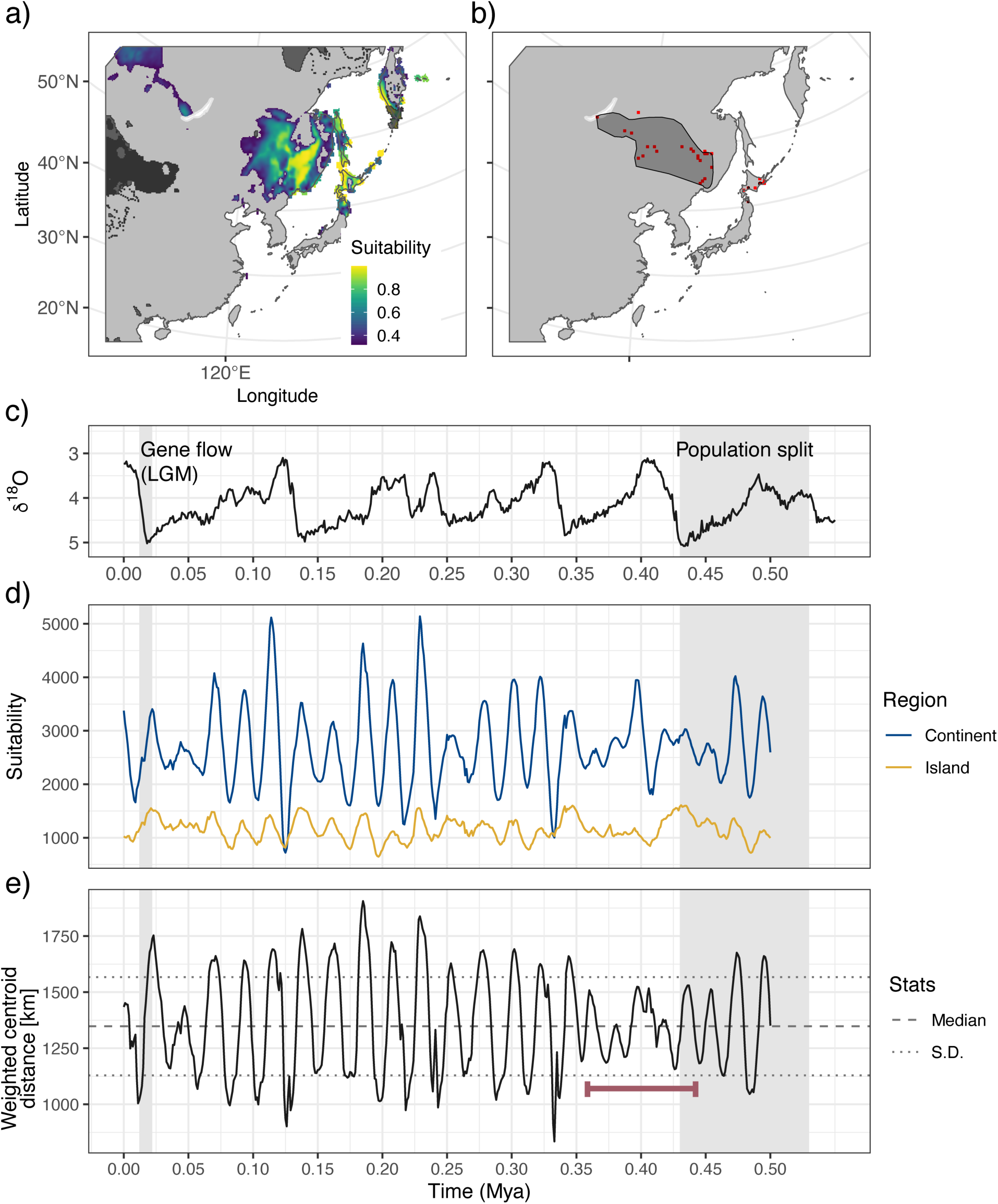
A predicted present distribution of Swinhoe’s Rail (a) by the species distribution model using its present occurrence records (b). (a) Color gradients from indigo to yellow represent environmental suitability for rails. Regions with dark gray shades are where model extrapolation was restricted due to the absence of the climate in the modeling domain (grids with negative MESS values). (b) The red grids indicate the used occurrence records at the resolution of the environmental layers, and gray areas indicate the expert range map of this species. (c) A temporal plot for a proxy for paleoclimatic temperature (*δ*^18^O; lower values indicate higher temperatures) (Lisiecki and Raymo 2005; Spratt and Lisiecki 2016). (d–e) Historical changes in the total suitability of the habitat in the continent (blue) and the island (yellow) (d) and the suitability-weighted centroid distance between the two (e). Black solid and dotted lines indicate the overall median and standard deviation of the distance, and a red bar indicates the period of the smallest amplitude in the temporal change. Grids covered by negative MESS values were also included in the calculation. Gray bars correspond to the estimated period of the population split (left) and gene flow (LGM, right). Lambert Azimuthal Equal Area projection was used for maps.

## Results

### Phylogenetic relationships and gene flow

Multiple phylogenetic methods collectively inferred population divergence. A bifurcation between an island clade (Tomakomai and Kushiro) and a continental clade (Russia and Aomori) was supported by TreeMix (Fig. 2c), SNAPP tree (Fig. 2d), and the D-statistic (Fig. S10). This structure was further supported by ADMIXTURE at K = 2 that scored the highest ΔK value (Fig. 1c, S6–7), the first axis of PCA explaining >99% of variation (Fig. 1b, S5), NeighborNet (Fig. S8), and F_ST_ (Fig. S9). On the other hand, some phylogenies indicated ILS and/or post-divergence gene flow. In the mtDNA tree, divergence between the two clades was supported, while Russian samples appeared in both clades, and most Tomakomai and Kushiro samples clustered in the island clade (except one from Kushiro) (Fig. 2a, S11). This pattern suggests either ILS due to the island population being derived from a large continental population or post-divergence gene flow from the island to the continental clade. Low bootstrap supports in the RAxML tree (Fig. 2b, S12–13) and low posterior probabilities for a bifurcation tree (41.5%) in the SNAPP tree (Fig. 2d) can also reflect this uncertainty. The divergence time of the two clades was estimated to be relatively old, around 0.5 Mya (ranging between 0.2–1.0 depending on analyses), corresponding to the Middle Pleistocene (the Chibanian epoch), collectively supported by different calibration methods, data, and algorithms, including the mtDNA tree (Fig. 2a), the SNAPP tree (Fig. 2d), and SWP2 (Fig. 3f).

Gene flow was also inferred by the best-fit population-level TreeMix tree with one migration edge scoring the highest *Δm* value (Fig. 2c, S14, S16), which was also supported by our individual-level tree (Fig. 2c, S15), indicating that gene flow inferred in our population-level analyses is not the artifact of population grouping (see Supplementary Discussion, Fig. S16– S19). Meanwhile, gene flow was directed from the common ancestor of this species to Russia. On our individual-level tree, the Tomakomai samples were located at the basal position. This suggests that ancient variation may have been introduced to the Russian population via gene flow from a population preserving ancient genetic variation, such as the Tomakomai population (Supplementary Discussion, Fig. S17–S19). This gene flow possibly occurred relatively recently (e.g., post-expansion of the continental population), as migration was not towards the ancestor of the continental cluster (Fig. 2c), but to Russia (at the population-level) or to an ancestor of one Aomori and one Russian sample (at the individual-level). One Russian individual clustered with island samples (Fig. 2c, inset), which was inferred to have the island cluster in our ADMIXTURE analysis at K = 3 (Fig. 1c), also supporting recent gene flow.

Aomori samples consistently clustered with and showed low genetic differentiation from Russian samples in all methods–phylogenetic trees (Fig. 2), PCA, ADMIXTURE (Fig. 1b–c), NeighborNet (Fig. S8), F_ST_ (Fig. S9), and the D-statistic (Fig. S10)–suggesting recolonization of Aomori by the continental population. This was unlikely to be a sample size artifact, as supported patterns held across genetic distance (NeighborNet) and F_ST_ that consider sample size and were robust to excluding Kanto wintering samples (i.e., smaller effect of sample size) (Fig. 1c, S5–S7). Additionally, the Baikal sample, collected ∼1,500 km from Amur, showed no major differentiation, despite the limited sample size. Together with the genetic position of Aomori, the present range of the continental population is a result of a recent expansion.

### Historical demography and contemporary genetic variation

The continental population had marginally higher genetic diversity than the island population (*p* = 0.089; Fig. 3a, Table S3). FSC2 results suggested that gene flow from the island contributed significantly to contemporary continental diversity. The best-fit FSC2 model (ΔAIC <2.0) was “different migration” model (ΔAIC = 0.0, Akaike model weight = 0.747), with “constant migration” as the second-best yet with a marginal ΔAIC value (ΔAIC = 2.17, Akaike model weight = 0.253). Both supported post-last-glacial-maximum (LGM, ∼0.02 Mya) gene flow (t_MIG_ = 0.014–0.019 Mya) from the island to the continent, at rates 1.8–2.7 times higher than in the opposite direction (Fig. 3c–e, Table S4). In contrast, gene flow before the LGM was inconclusive due to significant differences between the two models.

Our demographic analyses suggested that the island population remained relatively stable, or, if any, experienced a modest level of expansion since the population split until the LGM, supported by Tajima’s *D* values close to zero (Fig. 3b, Table S3), and FSC2 (0.5–2.3-fold change), and SWP2 (∼1.9-fold change) (Fig. 3d–e, Table S4). Meanwhile, different methods showed discordant results for the demographic history of the continental population. Expansion after a strong ancient population bottleneck was suggested by the significantly low value of Tajima’s *D* close to −1 (median value = −0.845) with a smaller inter-scaffold variance than the island population (Fig. 3b) and the best-fit FSC2 models (nearly 98% population decline, Fig. 3d–e), while expansion soon after the population split around 0.5 Mya without a population bottleneck was suggested by SWP2.

A sharp post-LGM bottleneck was suggested for the island population (97–99% reduction; Fig. 3d–e) by the best-fit FSC2 models. Our SWP2 results did not contradict it despite a large uncertainty towards the present (Fig. 3f). This was also supported by the small contemporary effective population size of Ishikari Plain in Hokkaido (including Tomakomai) (*N_e_* = 4,840) calculated from the census population size estimates around the early nineteenth century, which well corroborated the results of the FSC2.

### SDMs

The best-fit model (response curve ‘LQH’ + regularization multiplier 2.0, AICc = 515.86, Akaike model weight = 0.77, the Area Under the Curve (AUC) = 95.2%) accurately predicted the present breeding ranges of Swinhoe’s Rail (Fig. 4a–b, S21). During historical periods, some areas did not resemble any climate in the modeling domain of this species, indicated by the negative values in the MESS analyses (dark gray-shaded grids in Fig. 4a, S22, see Supplementary Methods). We restricted our interpretations to results robust to different treatments (inclusion or removal) of negative MESS grids.

The island served as a relatively stable glacial refugium, albeit smaller in extent than the continent. Continental habitats underwent drastic and repeated changes in habitat suitability and availability since 0.5 Mya (Fig. 4d). Habitat quality remained consistently higher on the island than on the continent throughout this period (Fig. S24). After the LGM, the island habitat contracted, corresponding to both SWP2 and FSC2, whereas the continental habitat expanded, only supporting the post-LGM demography of FSC2 (Fig. 4d, S22, S24). The temporal change in the level of habitat isolation between the island and the continent well supported the times of divergence and gene flow estimated by our genetic analyses. The 0.38–0.43 Mya interval showed stable isolation without periods of extreme proximity between the two regions, corresponding to the divergence period (Fig. 4e, S22, S25). The LGM was the most recent period of reduced isolation, consistent with post-LGM gene flow. Overall trends in these measures were similar between different treatments of negative MESS grids, although the amplitude of their changes differed between each other (Supplementary Results and Discussion, Fig. S24–S25).

## Discussion

Using an integrative phylogeography in Swinhoe’s Rail, we found that the contemporary continental population exhibited larger effective population size than the island population, consistent with general expectations (Leroy et al. 2021). However, multiple lines of evidence suggested that, historically, the island population was more stable and possibly larger than the continent, and that gene flow occurred from the island to the continent. These findings suggest that the island served as an important refugium for this species. In contrast, the direction of population establishment at the time of divergence remains unclear. Some phylogenetic trees supported bifurcating topologies at the population level, but individual-level analyses did not support reciprocal monophyly, leaving colonization direction unresolved.

### Gene flow and incomplete lineage sorting in the continent-island system

Incongruent phylogenetic patterns across analyses suggest a complex history of this species in the continent-island system, involving two non-mutually exclusive mechanisms, ILS and post-divergence gene flow. In recently diverged populations like Swinhoe’s Rail, gene genealogies may remain shared between two geographic populations (Twyford and Ennos 2012). For example, higher uncertainties were involved in the inference of the ancestral divergence and gene flow, including low posterior supports for bifurcation in the SNAPP tree and lack of reciprocal monophyly in the individual-level trees. These may reflect ILS, low marker resolution, or a combination of the two (Maddison and Knowles 2006), where ILS, derived from ancestral genetic variation, can obscure ancient evolutionary events (Twyford and Ennos 2012). Therefore, ancient gene flow inferred by FSC2 may be confounded with ILS, and thus its presence remains inconclusive. Nevertheless, the absence of monophyly due to ILS does not negate the existence of the two genetic clusters (i.e., continental and island populations), especially given multiple lines of support, and ILS is commonly retained to various degrees across populations and species (Maddison and Knowles 2006).

By contrast, recent gene flow is more readily distinguished from ILS (Twyford and Ennos 2012), and multiple results are consistent with this notion, supporting the presence of post-LGM gene flow from the island to the continent. First, the continental population showed signatures of strong historical bottlenecks–e.g., low Tajima’s *D*, FSC2 results, and SDM-inferred habitat instability–reducing the likelihood of retained ancestral variation. This is because a strong reduction in population size likely causes a fixation of genetic variants. Given the historically stable and relatively large island population, the island population may have preserved ancient genetic variations, causing ILS (see Supplementary Results & Discussion). Second, two-dimensional-SFS-based approaches, such as FSC2, and TreeMix– both of which explicitly account for ILS by estimating the level of genetic drift (Pickrell and Pritchard 2012; Murray et al. 2025)–consistently inferred gene flow to the continent. Therefore, while the relative contribution of gene flow versus ILS to the continental genetic variation remains unclear, our findings at least support the presence of recent gene flow from the island to the continent.

Discrepancies in population size histories between FSC2 and SWP2 may also reflect this gene flow. In demographic trajectory inference methods, such as SWP2, migration can inflate effective population size estimates, even without true expansion (Nadachowska-Brzyska et al. 2022). Therefore, the genetic bottleneck inferred in FSC2 might have been concealed in SWP2 by gene flow to the continental population. Alternatively, our simplified FSC2 models may have failed to capture complex fluctuating histories and rather showed averaged trends (long-term *N_e_*, Nadachowska-Brzyska et al. 2022), which may have exaggerated the impact of a bottleneck. However, because a long-term *N_e_* reflects a harmonic mean of a demographic trajectory (Nadachowska-Brzyska et al. 2022), the effect of past bottlenecks on the continental genetic variation should not be undervalued.

### Population persistence through a continent-island metapopulation

Historically large island populations and gene flow from islands have rarely been expected in continent-island systems. This is because small island size is typically assumed to limit population size (Leroy et al. 2021), leading to low dispersal potential and resultant gene flow (Poethke and Hovestadt 2001; Matthysen 2005). However, based on our FSC2 estimates, ancestral island population retained a large *N_e_* that was comparable to that of avian continental populations (mean *N_e_* = 362,456) (Leroy et al. 2021) and large tropical island populations, such as New Guinean birds (mean *N_e_* = 325,000) (Pujolar et al. 2022), whereas the *N_e_* of the ancestral continental population was as small as that of oceanic island populations (mean *N_e_* = 94,944) (Leroy et al. 2021). These differences may reflect contrasting habitat stability; our SDMs suggested more stable conditions on the island throughout glacial cycles than on the continent (Fig. 4d, S24), which could have allowed the persistence of a stable population and promoted dispersal (Ronce et al. 2000), leading to gene flow to the continent. In a large population, adaptive or ancient variation can be more likely to be preserved than in a small population where weakly deleterious mutations accumulate (Leroy et al. 2021; Lai et al. 2019). Asymmetric gene flow from such a large population may restore genetic variation and result in demographic and evolutionary consequences for a small population (Fitzpatrick et al. 2016). Therefore, the island population could have potentially contributed to the evolution and persistence of the continental population in this species. However, it remains unknown to what extent gene flow has contributed to the continental expansion, as the signature of gene flow was not found originating from the entire continental population (Fig. 2c).

Aomori of Honshu Island was inferred to have been recolonized recently in the wave of continental expansion, suggesting the possibility that the island population has also been sustained by migrants from the continent. Honshu might have been occupied previously by the present Hokkaido (“island”) population during a glacial period, supported by 1) the potential presence of suitable habitats on Honshu but not on Hokkaido during some glacial maxima inferred by the SDM (Fig. S22), and 2) a high affinity of Kanto samples towards Hokkaido breeding birds (Fig. 1b–c), given that wintering regions of short-distance seasonal migrants may reflect the past breeding range (Zink and Gardner 2017). Indeed, the population contraction of Swinhoe’s Rail in Japan reflects not only the post-LGM environmental changes but also anthropogenic factors, such as habitat loss and fragmentation due to land use changes (Kitazawa et al. 2022). This was also supported by our SDMs that predicted present suitable areas larger than currently recognized (Fig. 4a–b) and that the current Aomori habitat originated from an abandoned rice paddy (Ministry of the Environment, 2023). Hence, continental immigration resulting in re-establishment rather than gene flow may reflect the extinction of local island populations under anthropogenic impacts.

Altogether, our findings suggest that Swinhoe’s Rail has persisted as a dynamic continent-island metapopulation, where the Japanese Archipelago was not merely a one-way sink but played a role as pivotal as the continent, functioning as potentially a demographic and genetic source. Following divergence during a period of prolonged geographic isolation (∼0.5 Mya; Fig. 5a), spatiotemporal climatic heterogeneity likely led the island and the continent to shift roles within the metapopulation: the island supported relatively a large, stable population that retained ancient genetic variation, while the continent underwent drastic and repeated demographic changes that resulted in smaller long-term population size (Fig. 5b–c). Source-sink roles may have shifted over time via gene flow and population re-establishment, potentially promoting persistence of the metapopulation in the evolutionary timescale (Fig. 5b–c). Our findings were different from many previous studies that found islands to harbor small genetic variation and *N_e_* and act only as recipients of continental migrants. This is possibly because these patterns were often observed in systems with strong phenotypic or genomic divergence on islands (e.g., Wang et al. 2016; Delmore et al. 2020; Recuerda et al. 2021), where post-divergence gene flow from islands may be prevented or overlooked. In contrast, studies have reported that multiple island populations are maintained with gene flow on archipelagic or land-bridge islands (Seki et al. 2007; Mapel et al. 2021), and our results may be an expansion of such systems to the continent-island setting. Some Japanese endemic lineages have extended their ranges to the continental Russian Far East (e.g., *Gallinago hardwickii*, Nechaev and Fujimaki 1998) or Korean Peninsula (e.g., *Cettia diphone*, Clement and Kirwan 2020). Gene flow from Japanese lineages to their continental counterparts has also been suspected, yet without explicit inferences of the underlying historical processes (*Locustella amnicola* and *L. fasicolata*, Drovetski et al. 2015; *Lanius cristatus*, Aoki et al. 2021). Similar continent-island systems are recognized in other regions (Bellemain et al. 2008; Garcia-R et al. 2017), suggesting the universality of such a system across the globe.

**Figure 5.**
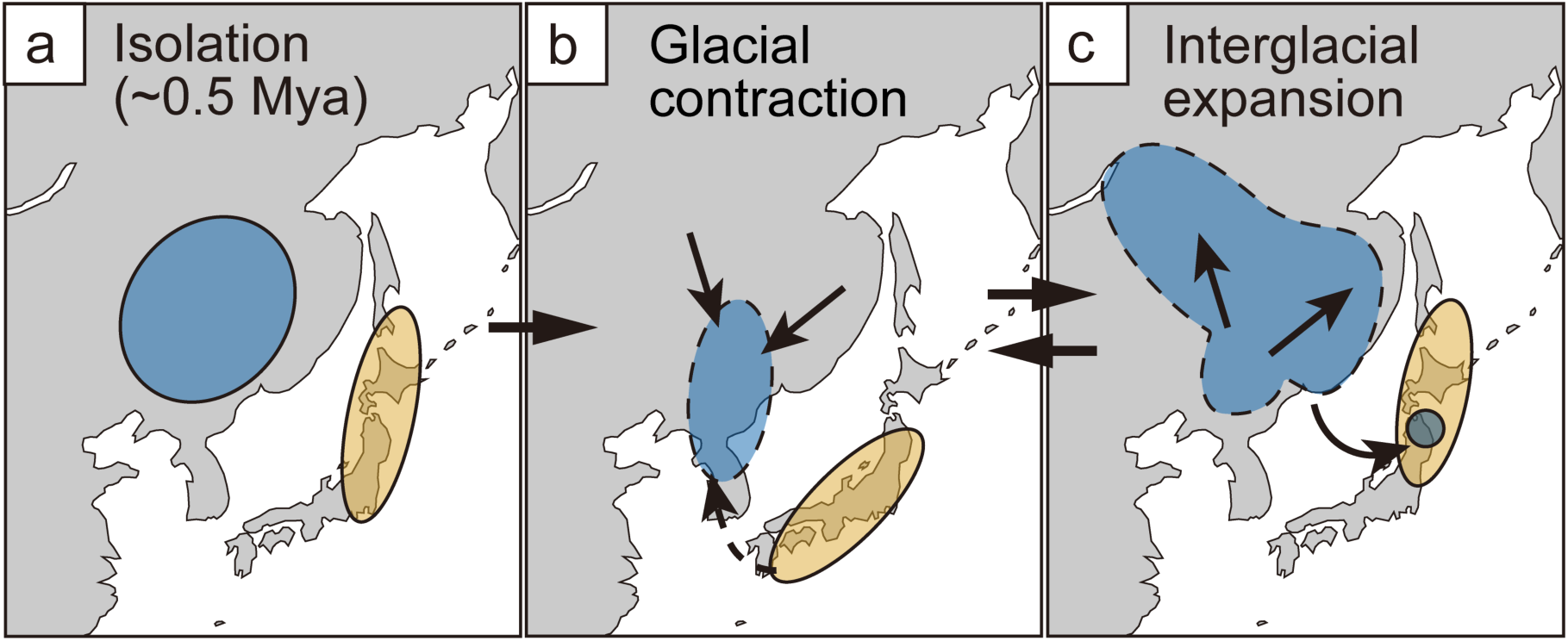
Schematic representation of our continent-island metapopulation scenario of Swinhoe’s Rail; geographic isolation occurred around 0.5Mya (a), and source-sink switching dynamics occurred in the continent-island system between the glacial (b) and interglacial period (c). Gene flow (a dotted arrow) and re-establishment or expansion of population (solid arrows) may have facilitated persistence and evolution of the metapopulation. Filled circles represent stable (solid) and unstable (hatched) populations.

A recent review emphasizes the importance of long-term population persistence in dynamic metapopulations for promoting speciation and adaptive radiation (Harvey et al. 2019). Our view also emphasizes the long-term consequences of metapopulation between continental and island populations, where source and sink relationships dynamically change, differently from the traditional perspective that focuses on the persistence of a metapopulation in the ecological timescale (Hanski 1998). A metapopulation structure across the continent-island system can be perceived as a transitional state for insular populations to undergo reverse colonization, thereby promoting long-term persistence and adaptation to the continental environment. Therefore, we highlight that our insights obtained from the population-level study may shed light on the origin of reverse colonization at the species level. Future research needs to incorporate more polymorphic sites to resolve ILS and gene flow and estimate the number, direction, and intensity of past migration, to understand the relative impact of these elements on the present genetic variation. This would disentangle the evolutionary importance of how a continent-island metapopulation may facilitate expansion, adaptation, and diversification on the continent, extending our microevolutionary idea to a macroevolutionary framework.

## Conclusion

Historical demography is fundamental to understanding a determinant of population genetic structure that is significantly linked with subsequent speciation and diversification (Harvey et al. 2019). In this study, we reconstructed the demographic history of a rail species in a continent-island system. We found strong evidence of asymmetrical gene flow from the Japanese Archipelago to the continental population, which may have influenced the continental genetic variation and demography. Furthermore, our results suggested that this species has persisted as a metapopulation spanning both the islands and the continent. These findings challenge conventional views of continent-island dynamics, particularly given that breeding populations of this species in Japan were only recently documented (Senzaki et al. 2020). To fully understand the processes that have given rise to the present biodiversity, evolutionary studies on the current and apparent species distribution are insufficient. Rather, reconstructing the long-term, spatially explicit population histories of organisms can shed light on the previously unfilled gap between micro- and macroevolutionary concepts and facilitate our understanding of global biodiversity patterns.

## Author Contributions

DA and MS conceived the ideas and designed the study with contributions by WH; MS, YO, WH, MK, and AF led the field work with substantial support by DA and MT; HA, NK, and DA led the laboratory procedures; DA conducted bioinformatics and analyzed the data with supports by HA and NK; DA coordinated the study, prepared the draft, led the writing and revised the manuscript with assistance from MS. All authors gave final approval for publication and agreed to be held accountable for the work performed therein.

## Acknowledgements

Some tissue and blood samples were kindly provided by the Yamashina Institute for Ornithology. We thank many members of our field teams; Toshio Sadakuni for sampling in Kushiro, Akio Miya and Jun-ichi Ebina for sampling at Hotokenuma, Aomori; Sergei M. Smirenski and Yury Anisimov for sampling in Russia; Shigeo Ozawa, Ryosuke Abe, Koichi Shirakawa, Daisuke Akoshima, Ayumu Hamachi, Hiroko Shirakawa, Mikiya Oikawa, and Shiomi Hakataya for sampling in the Kanto region. We also thank Nobuyoshi Nakajima for his support on sequencing runs on MiSeq. Our field surveys were conducted with permission from the landowners of each site, including Tomatoh. DNA samples were collected with permission from the Ministry of the Environment Government of Japan (permission # includes 21-31-0172-74, 19-129, 1810254, 2012217-16). We also acknowledge the provision of rings by the Bird Ringing Centre of Russia. Transportation of DNA samples was conducted under a Material Transfer Agreement between the National Institute for Environmental Studies and the Amur Bird Project based on the Access and Benefit-Sharing (ABS) agreement in the Nagoya Protocol. Bioinformatics was mostly performed on the supercomputer of AFFRIT, MAFF, Japan.

## Data Availability Statement

Scripts and data for analysis are available on Dryad: 10.5061/dryad.37pvmcvs1 (private link: https://datadryad.org/stash/share/2PRy64eM1UIjsjCJlNLx4B9ownPLtYVob4aFp_1MXys). Raw sequence reads are deposited to Sequence Read Archives (see Table S1). Settings and procedures for these processes are fully described in the main and supplementary texts and the original scripts.

