## supplementary_materials for "Phylogenomics supports island contribution to metapopulation dynamics in a predominantly continental bird species"

#### Title

#### Methods

##### *Materials*

Almost all of the samples of the Swinhoe's Rail were collected by us, except for the Ishikari sample (Accession number: LC781943). This sample was collected in a condition where it had become stranded along the coastline of Ishikari Bay. This sample is stored at the Yamashina Institute for Ornithology, and its mitochondrial DNA was once determined by the other authors (Ozaki et al. 2010). However, as has been pointed out by Heim et al. (2019), its phylogenetic position was presumed to be wrong, and therefore, we re-sequenced the *Cytb* gene of this sample together with the other samples.

##### *Laboratory procedures*

We conducted DNA extraction by using the DNeasy Blood & Tissue kit or QIAamp DNA Micro Kit (Qiagen) following the manufacturer's protocol. The same extracted DNA for each sample was used for both mitochondrial and MIG-seq experiments. Sequencing procedures for Sanger sequencing and MIG-seq are as follows.

**Sanger sequencing of a mitochondrial gene:** We used the standard PCR and BigDye terminator protocols to sequence the *Cytb* gene. We used the same primer pairs as Heim et al. (2019) did for Russian populations: forward primer (L14764: 5′-TGRTACAAAAAAATAGGMCCMGAAGG-3′ (Sorenson et al. 1999) or mt-C2: 5′-TGAGGACAAATATCATTCTGAGG-3 (Fritz et al. 2006)), reverse primer (mt-C4: 5′-AGTGTTGGGTTGTCTACTGA-3′ (Broders et al. 2003) or mt-FSH: 5′-TAGTTGGCCAATGATGATGAATGGGTGTTCTACTGGTT (van der Bank et al. 1998)) and followed PCR conditions from Ando et al. (2018).

**MIG-seq:** Using the originally described MIG-seq primer set (Suyama and Matsuki 2015), we first conducted the multiplexed PCR with tailed inter-simple sequence repeat (ISSR) primers, which consist of eight forward and eight reverse primers. This gave 64 SSR combinations to sequence ISSRs. We prepared a reaction mixture of total volume 7.0 µL, containing 1.0 µl template DNA, 3.5 µL 2X PCR buffer, 0.2 µM each of the 16 first PCR primers, 0.035 µL Multiplex PCR enzyme mix, and 0.225 µL of nuclease-free water, using Takara Multiplex PCR Assay Kit ver. 2 (Takara, Japan). The PCR condition was as follows: an initial denaturation for 1 minute at 94ºC, 25 cycles of PCR reactions consisting of 30 seconds of denaturation at 94ºC, 1 minute of annealing at 48ºC, and 1 minute of extension at 72ºC, and a final extension for 10 minutes at 72ºC. We repeated three independent runs of PCRs, and they were combined into one tube for each sample to increase the chance of PCR amplification. 10 µL of the combined mixture was used for purification with 18 µL of AMPure XP (Beckman Coulter, USA), which was rinsed twice with 70% ethanol. Finally, DNA captured by beads was eluted with up to 40 µL of TE, which was used in the second PCR. The reaction mixture of a total volume of 15 µL per sample consists of 3.0 µL 5X PrimeStar GXL buffer, 1.2 µL dNTP mixture, 0.3 µL second PCR forward primer, 0.3 µL PrimeSTAR GXL Polymerase, 6.7 µL of nuclease-free water, 1.5 µL of the uniquely indexed reverse primer, and 2 µL of the first PCR products after purification. The second PCR consists of 12 cycles of 10 seconds of denaturation at 98ºC, 15 seconds of annealing at 54ºC, and 60 seconds of extension at 68ºC. Amplification of PCRs was checked by agarose gel electrophoresis, and additional cycles of PCR were conducted for samples amplified less effectively. A volume of 5–8 µL of each second PCR product was combined into one 1.5 mL LoBind Tube, whose 100 µL was used in PCR purification using QIAGEN QIAquick PCR purification (QIAGEN, Germany), following the manufacturer's protocol. The purified second

PCR product was then subjected to size selection using SPRI Size Select (Beckman Coulter, USA) to obtain a fragment size of 300–800 bp, and DNA was eluted in 30 μL of TE buffer. To ensure that the fragment size is within the 300–800 bp range, size selection was also repeated using a 1.8% agarose gel, which was purified by QIAGEN Gel Extraction Kit (Qiagen, Germany) following the manufacturer's protocol. The purified solution was concentrated to 20 μL, which was applied to an Agilent BioAnalyzer (Agilent Technologies, USA) to measure its concentration. The solution was diluted to 4nM with deionized water. The final mixture was applied to MiSeq Reagent Kit v3 150 cycles (Illumina) to obtain 150 bp paired-end reads. All the nucleotide sequences and raw reads obtained were deposited in DDBJ (Table S1).

*Bioinformatics*

Raw reads from each indexed sample from MIG-seq were grouped, and adaptor sequences were removed simultaneously, using the index reads option of the sequencer. We quality-checked contamination of unremoved adapters and primers on the pre-trimmed raw reads by FastQC (http://www.bioinformatics.babraham.ac.uk/projects/fastqc) and MultiQC (Ewels et al. 2016). The raw FASTQ reads for each sample were trimmed and filtered using Trimmomatic v. 0.39 (Bolger et al. 2014). In the Trimmomatic quality filtering, options included the following; the paired-end option was applied with both 'TRAILING' and 'LEADING' set to 30, removing bases below 30 sequencing quality score at both trailing and leading edges, and 'MINLEN' set to 40 for sequences with length at least 40 bps to be retained.  Paired and unpaired reads detected through quality filtering were individually aligned with a paired-end and single-end option, using the BWA-MEM algorithm of Burrows-Wheeler Aligner (Li 2013), respectively, and they were then combined by SAMtools v.1.16.1 (Danecek et al. 2021). Reads aligned to the reference genome were directed to SAMtools (Danecek et al. 2021) to output them as BAM files. The bam files of both paired and unpaired reads were combined using the 'merge' function of SAMtools, which was sorted using a function 'sort' generating a bam file for each sample to be used in the population genomics analyses.

Individuals with a large proportion of missing data will decrease the number of potential polymorphic loci (Cerca et al. 2021) and cause wrong phylogenetic signals (Smith et al. 2020). Therefore, we calculated the levels of missingness at potentially variant sites for each sample by conducting ANGSD pre-analysis, with filtering options of '-SNP_pval' = 1E-06, '-Minmaf' = 0.01, and '-minind' = 26, and other settings same as the main ANGSD runs (see the Main Text). We determined samples that were with missing data proportions >30% but were not collected in the less intensively sampled population (Aomori) as those to be removed (i.e., two Kanto samples and one Russian sample, Fig. S4). We also determined relationships between the minimum depth thresholds to discard sites and the number of reads that can be retained by preliminary SNP calling using BCFtools v. 1.8 (Danecek et al. 2021). We set '-q', '-Q', and '-P' to be 30, 20, and 1e-6, respectively as filtering options, and called single nucleotide polymorphisms (SNPs) by using a range of minimum depth values (DP = 1, 2, 4, 7, 10, 15). We plotted changes in the per-sample mean proportion of missingness (Fig. S1a), per-sample mean depth (Fig. S1b), mean sample coverage (Fig. S2a), and the distribution of the number of sites for each size of sample coverage (Fig. S2b), against different minimum depth values. The number of sites retained decreased as DP was increased (Fig. S1–S2). In order to compensate for this trend, we rescued sites with low depths while with several filtering schemes to reduce biases from these sites, especially in the combinatory use of '-minInd' and '-setMinDepthInd' (Lou et al. 2021). By conducting population-specific ANGSD analyses and obtaining intersecting sites among them, we removed sites recovered specifically for some of the populations. We set the '-

minInd' option to be half of the maximum number of individuals of each population, to ensure that sites are well covered by samples of each population. We also set '-setMinDepthInd' to be 2 (see Main Text), determined by referring to the other study that used a similar analysis flow (Capblancq et al. 2020).

Raw reads downloaded from the sequence read archive (SRA) were trimmed, quality checked, and aligned against a reference genome by following a similar procedure described for the MIG-seq reads. Exceptions included the following: the 'MINLEN' option in Trimmomatic was adjusted by considering the average read lengths of each SRA FASTQ file; the aligner software bwa-mem2 v. 2.2.1 (Vasimuddin et al. 2019) was used with the AVX512 model to map short reads for fast alignment.

Additional parameters for genotype likelihood (GL) calculation by ANGSD were as follows. In all the genotype likelihood calculations, the following filtering options were included; the base with unique best hits ('-uniqueOnly' 1), alignment quality (BAQ) option 2 (Li 2011), minimum mapping quality ('-minMapQ') of 30, minimum base quality ('-minQ') of 20, skipping triallelic sites. For removing linkage disequilibrium by ngsLD (Fox et al. 2019), we set options max_kb_dist and min_weight to 50 and 0.2, respectively.

To include outgroups in these analyses, we also calculated GLs for the reads that were downloaded from the SRA and manipulated in the aforementioned procedure. Outgroup sequences downloaded through the Sequence Read Archive included *Laterallus jamaicensis* (reference sample) and *Atlantisia rogersi* (Table S1). To reduce the computation time, we generated a subset of the BAM files that include sequence reads mapped against the reference genome, only to include intersecting sites that were found in ANGSD analyses in each dataset.

*Global genetic structure*

**PCA and ADMIXTURE:** For the principal component analysis (PCA), 1,000 iterations with a minimum major allele frequency ('-minMaf') of 0.05 were used as settings, and we confirmed that any site used to conduct PCA was not under strong natural selection with the '--selection' option by checking whether there are any positions with the signature of selection below an alpha value 4.17e-05 after Bonferroni correction and conducting the chi-square test. For ADMIXTURE, 10 independent runs for each K, ranging from 1 to 6, were conducted with a maximum of 5,000 iterations and a minimum major allele frequency of 0.05. We calculated ΔK values, the rates of change in the log probability of data between successive K values, by following Evanno et al. (2005), and the best K value that scored the highest ΔK was determined. For plotting, we used the best-run scoring the maximum likelihoods for each K.

In addition to the PCA and ADMIXTURE analyses using both breeding and wintering samples (Fig. 1b–c), we also conducted these analyses on different datasets, including 1) only breeding samples, 2) both breeding and wintering samples with two outgroup samples (*Laterallus jamaicensis*, *Atlantisia rogersi*), and 3) breeding samples with outgroup samples. For each analysis, GLs were recomputed by ANGSD, and the same parameter settings were used for these analyses.

**NeighborNet**: We calculated pairwise genetic distance between samples by using ngsDist (Vieira et al. 2015). We used the beagle output of the ANGSD run conducted for the PCA and ADMIXTURE analyses using a dataset including both breeding samples and outgroup species. We calculated the average *p*-distance for each pair of samples while taking into account genotype uncertainties by using genotype likelihoods. A NeighborNet plot was constructed by a function of ‘neighborNet’ implemented in the R package ‘pahngorn’ v. 2.11.1 (Schliep 2011), which was visualized by using SplitsTree4 (v. 4.19.29) (Huson and Bryant 2006).

**D-statistic**: We calculated the D-statistic for each trio of breeding populations (Tomakomai, Kushiro, Russia, and Aomori) with one outgroup (*Laterallus jamaicensis*) to determine the correct tree topology or presence of gene flow. By setting two populations of the trio as ingroups (H1 and H2), the D-statistic determines whether one of the ingroups is unexpectedly closer to the non-ingroup population (H3). If a correct topology of H1, H2, and H3 is given, a statistically significant D-statistic value indicates gene flow between either H1 and H3 (negative D-statistic value) or H2 and H3 (positive D-statistic value). If an incorrect topology of the trio is given, the D-statistic also becomes statistically significant.

*Inference of phylogenetic relationship*

**mtDNA tree and haplotype network:** The HKY model (Hasegawa et al. 1985) with the gamma rate heterogeneity ($\alpha = 0.14$) was selected as the best substitution model scoring the lowest Bayesian information criterion value calculated by MEGA v. 11.0.13 (Tamura et al. 2021). A pre-burn-in included 10,000 chains to be discarded prior to the 2e+07 generations of the main chain. A maximum clade credibility tree was drawn using the R package ggtree v. 3.6.2 (Yu et al. 2017). We only annotated nodes with posterior probabilities >0.9 with black-filled circles.

A haplotype network was reconstructed using NETWORK v. 10.2.0.0 (Bandelt et al. 1999; Polzin and Daneshmand 2003), following the procedure in Aoki et al. (2021).

**RAxML tree**: We used the GTGTR4+FO+G model to reconstruct the maximum likelihood tree using RAxML-ng. GTGTR4 is suitable for our dataset that consists of genotype states without diploid unphased and has fewer parameters than GTGTR by not distinguishing genotype states (mutation rates from A/T, T/A, and T/T to another are all equal). The FO option was applied to use the stationary frequencies of genotypes estimated by maximum likelihood methods. The "+G" option was applied to consider the mutation rate varying among sites on the genome.

Similarly to the PCA and ADMIXTURE analyses, we reconstructed maximum likelihood trees using RAxML with different datasets, including 1) both variant and invariant sites from only breeding samples (Main Text, Fig. 2b), 2) both variant and invariant sites from both breeding and wintering samples, 3) only variant sites from breeding samples, and 4) only variant sites from both breeding and wintering samples. We used the same settings for the RAxML run described in the Main Text. To evaluate the effect of missing data on the inference of tree topology, we plotted the tree with the missing proportion of each sample by using ggtree v. 3.6.2. Then, we also re-ran the RAxML using datasets generated based on the ANGSD run with setting the minimum number of individuals for each site ('-minInd') to 80% of the number of entire samples for each dataset, and we compared the tree topologies with the proportion of missingness.

**TreeMix**: We determined the block size for the jackknife as the half-life of the linkage disequilibrium (LD) decay, which was estimated by ngsLD. The LD calculated for the PCA analysis was filtered to retain sites that the glactools selected as non-private variant sites in the 'acf2treemix' function. We fitted nonlinear models of LD decay for $r^2$ values against the physical distance between sites according to Remington et al. (2001) and estimated the recombination parameter ρ, which was used to draw the LD decay curve for our dataset. We found the half-life of the decay as the physical distance where the $r^2$ value becomes half of the initial value at physical distance = 1. We determined 22 and 19 as the block size for population-based and individual-based TreeMix models, respectively.

We conducted 10 replicates of runs with each number of migration edges ranging from 0 to 5. We evaluated the model performance by examining both standard metrics and conducting post-hoc analyses using the R package 'OptM' v. 0.1.6 (Fitak 2021). We calculated the total variance that each model explained in the data, using the R function 'calcVarExplain'. For each number of migration edges, means and standard errors were calculated for the total variance that each model explained. We also calculated means and standard errors for log-likelihoods over 10 replicates of each migration edge. We plotted the percentage variance explained and log-likelihoods against the number of migration edges. We calculated $\Delta m$ values that were similar to $\Delta$K values in the ADMIXTURE analysis to find the best number of migration edges. Higher $\Delta m$ values indicate the model likelihood increases significantly between successive numbers of migration edges. We determined the model performance by evaluating model likelihoods, total variance in population relatedness, and residual fits (Pickrell and Pritchard 2012; Fitak 2021), and model consistency was also checked among different replicates at the best number of migration edges.

**SNAPP tree**: From the unlinked genotypes called using ANGSD and ngsLD, we filtered out sites that showed excess heterozygous sites using BCFtools. This reduced the number of sites to 206 from the originally called unlinked SNPs (n = 246). We generated an input XML file by using a script called "snapp_prep.rb" provided by Stange et al. (2018) (available at https://github.com/mmatschiner/snapp_prep).

*Demographic inferences*

**Site frequency spectrum (SFS) and summary statistics:** We used the '-doSAF' option of ANGSD and the 'realSFS' function to estimate the folded SFS. We set the max iteration ('-

maxIter') to 50,000 and tolerance for optimization processes ('-tole') to 1e-06. The functions 'saf2theta' and 'do_stat' of the 'thetaStat' subprogram were used to compute a genetic diversity index (Watterson's theta) and a neutrality statistics index (Tajima's *D*), using the folded SFS as inputs.

**$F_{ST}$:** Weighted $F_{ST}$ (Reynolds et al. 1983) was calculated using the 'realSFS fst' function of ANGSD on two-dimensional SFS (2dSFS) among each pair of four breeding populations. The 2dSFS were calculated only using unlinked sites, determined for the ADMIXTURE and PCA analyses.

**Linear modeling:** To construct a linear model, we used the R package lmerTest v. 3.1.3 (Kuznetsova et al. 2017). We set population (the island and continental populations) and log-transformed scaffold size as covariates to explain the response variable of per-site diversity indices or neutrality test statistics that were calculated by dividing the output values calculated for each scaffold by the number of sites. The island population was set as the base of a dummy variable to be compared with the continental population. We removed scaffolds <2,000,000 bp from the models.

**Contemporary effective population size:** The census population size has been estimated as $N_c$=12,100 for Swinhoe's rails in Ishikari Plain (where the Tomakomai region is included) before human environmental modification (170 years before present) (Kitazawa et al. 2022). This value was used as the standard population size in the contemporary environment and used to calculate $N_e$ (see Main Text).

*SDMs*

We compiled an occurrence dataset mainly from previously published literature (Table S6). In addition to the literature mentioned in the Main Text reporting the breeding records of this species, Mlíkovský et al. (2024) reviewed both breeding and non-breeding records from the literature, museum specimens, and multiple online databases. Furthermore, we added newly discovered breeding and non-breeding records of this species from Japan (e.g., Sadakuni et al. 2018). For the background sampling extent, we generated a 1,000-km buffer around a convex-hull of both breeding and non-breeding records, because we intended to model historical range shifts of a potentially dispersive bird species. Given that natal dispersal distance in birds is large (median = 7.74 ± 10.49 km/individual/year; Fandos et al. 2021), we decided to extend the sampling extent to include most of East Asia by setting this buffer.

We used climatic covariates, including 17 of 19 annual bioclimatic variables, annual net primary productivity, altitude, rugosity, leaf area index, and biome, at a spatial resolution of 0.5° × 0.5°. We used the 'vifstep' function in an R package, usdm v. 2.1.7 (Naimi et al. 2014), to remove highly correlated variables. Eight variables were selected, including bio04 (temperature seasonality), bio07 (temperature annual range), bio10 (mean temperature of warmest quarter), bio13 (precipitation of wettest month), bio14 (precipitation of driest month), leaf area index (lai), altitude, and rugosity.

We used the 'ENMevaluate' function of an R package ENMeval v. 2.0.4 (Kass et al. 2021) with the 'maxnet' option, with 'block method' as a partition method for cross-validation, and each combination of the regularization multiplier ranging between 1 and 3 and four different feature classes determining the shape of response curves ('L', 'LQ', 'H', and 'LQH') was tested.

The best model was selected based on the lowest AICc, and presence and absence data were used to determine a presence/absence threshold of this species by using the 'eval.results' of ENMeval and 'threshold' functions of predicts v. 0.1.17 (https://github.com/rspatial/predicts), which was used to clip the model results projected to the present and past climates. Multivariate environmental similarity surface (MESS) (Elith et al. 2010) was calculated by using an R package 'predicts' v. 0.1.17, and cells with negative MESS values, indicating that pairs of the covariate values were not present in the modeling domain, were restricted for model projection and semi-transparent dark gray colors were overlaid on the prediction of Fig. 4 and Fig. S22. The model projection to the past climate layers was done using the 'predict' function of terra v. 1.8.21 (Hijmans et al. 2022), with the 'cloglog' transformation and clamping. MESS was recalculated for the projected layers to avoid overinterpretation of the model projection.

We quantified performance and significance of our SDM by using a null model approach (Bohl et al. 2019), using a function 'ENMnulls' of ENMeval. We provided our model object and the model settings as inputs, and 100 iterations were run to obtain null model performance metrics. We plotted the distribution of metrics of the null models with the real model output to compare the model performance, using the 'evalplot.nulls' function of ENMeval.

We calculated and compared the total suitability, availability, and quality of habitat on the continent and the island by using the SDM projection to the past climate layers. We defined the continent and the island by creating polygons (Fig. S23): The island polygon was defined to include Sakhalin, the Kuril Islands, and the Japanese Archipelago; the continental polygon was defined to cover the background extent, which was created to include the present breeding and non-breeding occurrence records on the continent (Fig. S20). Although the definitions of the "continent" and the "island" change in historical periods due to the formation of land bridges and the emergence of continental shelves, we did not include these areas in our calculations for the following two reasons. First, emerged sea floors (e.g., land bridges) were likely to be located in the middle of the present continent and island areas, and hence, these areas were likely to have been colonized by both the continental and island populations. Under this assumption, the same values will be added to the indices of continental and island habitats, which would not change the absolute differences in the calculated values between the continental and island habitats across time. Second, most of these areas were predicted to be unsuitable for this species (Fig. S22); thus, inclusion of the areas would not largely impact our inferences.

The centroid distance between the continental and island habitat was calculated as the measure of the level of isolation. We weight centroid distance by taking into account predicted values of each habitat grid in the following equation:

$$x_{centroid} = \frac{\sum(x_i \times w_i)}{\sum(w_i)}, y_{centroid} = \frac{\sum(y_i \times w_i)}{\sum(w_i)}$$

This equation considered the areas where suitable habitat was concentrated to calculate the distance between the two regions. The centroid distance was calculated in meter by using an equal-area projection map.

#### Results & Discussion

*The phylogenetic position of the stranded Ishikari sample*

The Ishikari sample was placed in one of the clades of Swinhoe's Rail ("island clade", Fig. 2a, S11) that were sister to the *Coturnicops noveboracensis* in our mitochondrial tree, which is contradicted by its position inferred based on the mitochondrial sequence previously determined (Ozaki et al. 2010; Heim et al. 2019). Therefore, we argue that the previous study wrongly reported its mitochondrial sequence either by a mix-up of a different sample or by any technical issues in the experiment. Nonetheless, we supported that the Swinhoe's Rail is a sister species of the *C. noveboracensis*, as has been previously suggested (Heim et al. 2019).

*The effect of missing data on tree topology in RAxML tree reconstruction*

The tree topologies were not affected strongly by the inclusion and exclusion of the Kanto samples (Fig. S12–S13). Furthermore, the proportion of missing data did not seem to affect the tree topologies either, given that mean missing proportion distributed randomly across the tree and that the values used to obtain '-minInd' (50% or 80%) did not seem to affect the tree topologies (Fig. S12–S13). However, these trees scored low bootstraps at their nodes (<70%), and therefore, the effect of missing data on the statistical confidence of trees remained unclear.

Nodal supports for the ML tree did not change the inferred tree topologies between a dataset consisting of both variant and invariant sites and that consisting of only variant sites (Fig. S12–S13). Branch lengths were relatively longer in variant-only trees, and this was possibly explained by trees reconstructed with ascertainment biases (Leaché et al. 2015).

*The breeding origin of Kanto samples*

While the results of PCA and ADMIXTURE suggested that Kanto birds were genetically related to Hokkaido populations, three of five mtDNA haplotypes from Kanto were found in continental clades or shared with Russian haplotypes (Fig. 2a, S11). Although it remained inconclusive on how these two mitochondrial clades diverged (see Main Text), if they originated from the glacial isolation of continental and insular populations, this may indicate a possibility that some Kanto birds migrated from an unsampled breeding population between Kushiro and continental Russia, such as Sakhalin Island. This is likely since Sakhalin Island was estimated to be suitable in our

SDM (Fig. 4a). In such a region, continental mitochondrial haplotypes may be introgressed to the island population while retaining insular genetic variation in nuclear DNA, given that there is a gap in suitable habitats of this around Khabarovsk region (coastal area of continental Russia) between Sakhalin Island and Amur or north China.

*Potential preservation of ancient genetic variation in the island population*

Although each support is relatively weak, multiple lines of evidence were collectively suggestive of the preservation of ancient genetic variation in the island population rather than the continental population. First, the second principal component explained variance within the Tomakomai population, despite the small percentage (0.54%) over all. Second, the ADMIXTURE analyses showed the presence of the continental cluster shared among the island samples ($K = 2$) despite a lack of consistent inferences of gene flow from the continent to the island in the TreeMix and FSC2 models. This may be explained by the recent bottleneck of the continental population and preservation of the ancient variation in the island population (Lawson et al. 2018). Third, the D-statistic inferred an asymmetrical relationship between the continent and island populations. When one of the continental populations (Aomori or Russia) was assigned as P3, and the other continental and one island population was assigned as P1 and P2, then the D-statistic was not significant (blue bars cross 0). On the other hand, when an island population (Tomakomai or Kushiro) was assigned as P3, then the D-statistic turned out to be significant. This may indicate that the shared genetic variants are similar between continent-island and continent-continent pairs, whereas this may not be true between island-continent and island-island pairs, suggesting that islands may have preserved ancient variation or the continental population has experienced severe genetic drift, losing ancient variation. Fourth, at the individual-based tree (TreeMix and RAxML), the position of the island and continental samples within trees switched depending on the consideration of gene flow, although the statistical support for these nodes may be weak.

*Model evaluation of TreeMix*

In the population-level tree, the total variance in the relatedness between populations explained by the model with migration edge = 0 was above 99.8%, and adding migration edges did not improve the variance explained by the TreeMix models any further (Fig. S14b–c). On the other hand, model likelihoods were improved drastically by adding one migration edge (Fig. S14a, d), and increasing the number of migration edges >1 did not further improve the model likelihoods (Fig. S14). According to Pickrell & Pritchard (2012), the 99.8% variation in the relatedness between populations is sufficient to infer the best number of migration edges. However, the number of migration edges could be underestimated by blindly relying on this threshold (Fitak 2021). Especially, our outgroup was relatively phylogenetically distant without the most closely related species, *C. noveboracensis*, leading to a larger proportion of genetic variance possibly being explained by interspecific genetic variation, resulting in higher variance explained with migration edge = 0. Adding one migration edge decreased unexplained genetic variance within the Swinhoe's Rail, while it increased unexplained variation in the outgroup (*Laterallus jamaicensis*) (Fig. S17), suggesting this possibility.

Although the source of gene flow to the Russian population remains inconclusive (see Main Text), the source is likely the Tomakomai population. Despite the low model likelihoods, only the replicates #1 and #2 showed as long external branches to the ancestor of Swinhoe's Rail as those to the outgroup (Fig. S19). Although the outgroup *L. jamaicensis* is a species isolated within a small island, and we expect high genetic drift, it is commonly expected that the level of overall drift may not strongly differ between the two species, as shown in most of the replicates of our TreeMix outputs (Fig. 2c, Fig. S19c–j). Given that the outgroup is phylogenetically distant from our target species and most of the genetic variance to be explained by the TreeMix is divergence between the two species, the long external branch to *L. jamaicensis* might be an artifact of the outgroup selection (Fig. S17). In this situation, the TreeMix model might have been fitted to the data to reduce the overall residuals of genetic variance, which explained the unshared variant between the species more likely by drift in the outgroup, scoring higher model likelihoods, than by drift in both external branches, scoring lower model likelihoods. Therefore, despite the low model likelihoods, the replicates #1 and #2 may reflect the actual phylogenetic relationships better than the other replicates, suggesting that the source of migration could be Tomakomai. This notion is also supported by the individual-level tree (see Main Text).

The addition of migration edge = 1 only improved inter-continent-island-sample genetic variance on the individual-level tree (Fig. S18). This suggests that the migration edge inferred in the individual-level tree with m = 1 basically inferred the same migration edge at the population-level tree with m = 1 (Fig. 2c). On the other hand, in the individual-level tree, unexplained variance was large overall (>0.6) (Figure S15). It is to be noted that the TreeMix is developed to model inter-population phylogenetic relationships by considering gene flow and drift. It does not intend to examine relationships of individuals within the same population that share many alleles among samples, but at different degrees among different pairs due to incomplete lineage sorting, and the total genetic variance cannot fully be explained by the best-fit tree.

*SDMs*

The extent of negative MESS grids, indicating that an environment was dissimilar to the present modeling domain, was pervasive in the historical periods (Fig. S22). Especially during the non-maxima of glacial periods (i.e., 0.05 Mya, Fig S22c; 0.461 Mya, Fig S22f), these grids greatly overlapped with the predicted presence grids, resulting in lower suitability and availability for this species both on the continent and island, when negative MESS grids were removed from the analyses (Fig. 4, Fig. S22, lower panels of Fig. S24). Because negative MESS values indicate that climates at the grids were not available in the present modeling ranges but do not always mean that they were not suitable for this species, how we consider these grids influence our inferences on the historical changes in the availability of the species' habitat and their differences between the island and continent. Therefore, the results of past climate projections need to be carefully examined. However, we can, at least, support our arguments 1) that the island was more stable than the continent, 2) that geographical isolation was prolonged around 0.38–0.43 Mya, and 3) that gene flow was likely to have occurred around the LGM, based on our SDM analyses, because these patterns were supported regardless of the treatment of negative MESS value grids.

### Supplementary Tables

**Table S1** Sample details. The Analysis column indicates the types of molecular techniques applied to each sample or used by other studies, including MIG-seq (MIG), mitochondrial sequencing (mt), and whole genome sequencing (WGS). The MIG-seq ID column indicates the positions of 96 wells in which libraries were prepared.

| Locality | Species | Analysis | Accession number (Cytb) | SRA Accession | Population | Date (YYYY/MM/DD) | Ring ID | Collector | Tissue | MIG-seq ID | Reference |
|---|---|---|---|---|---|---|---|---|---|---|---|
| Kamisu, Ibaraki, Japan | *Coturnicops exquisitus* | MIG | - | DRR505881 | Kanto (wintering) | 2018/01/14 | 3E42033 | OY | blood | S001 | This study |
| Kamisu, Ibaraki, Japan | *Coturnicops exquisitus* | mt, MIG | LC781682 | DRR505882 | Kanto (wintering) | 2018/02/03 | 3E42035 | OY | blood | S002 | This study |
| Kamisu, Ibaraki, Japan | *Coturnicops exquisitus* | MIG | - | DRR505883 | Kanto (wintering) | 2018/02/03 | 3E42036 | OY | blood | S003 | This study |
| Inashiki, Ibaraki, Japan | *Coturnicops exquisitus* | MIG | - | DRR505884 | Kanto (wintering) | 2018/02/03 | 3E42037 | OY | blood | S004 | This study |
| Inashiki, Ibaraki, Japan | *Coturnicops exquisitus* | mt, MIG | LC781696 | DRR505885 | Kanto (wintering) | 2018/02/03 | 3E42038 | OY | blood | S005 | This study |
| Kamisu, Ibaraki, Japan | *Coturnicops exquisitus* | mt, MIG | LC781697 | DRR505886 | Kanto (wintering) | 2018/02/12 | 3E42039 | OY | blood | S006 | This study |
| Kamisu, Ibaraki, Japan | *Coturnicops exquisitus* | MIG | - | DRR505887 | Kanto (wintering) | 2018/02/12 | 3E42040 | OY | blood | S007 | This study |
| Kamisu, Ibaraki, Japan | *Coturnicops exquisitus* | MIG | - | DRR505888 | Kanto (wintering) | 2018/02/12 | 3E42041 | OY | blood | S008 | This study |
| Kamisu, Ibaraki, Japan | *Coturnicops exquisitus* | mt, MIG | LC781698 | DRR505889 | Kanto (wintering) | 2018/02/12 | 3E42042 | OY | blood | S009 | This study |
| Kamisu, Ibaraki, Japan | *Coturnicops exquisitus* | mt, MIG | LC781699 | DRR505890 | Kanto (wintering) | 2018/02/12 | 3E42043 | OY | blood | S010 | This study |
| Inashiki, Ibaraki, Japan | *Coturnicops exquisitus* | mt, MIG | LC781683 | DRR505891 | Kanto (wintering) | 2018/02/12 | 3E42044 | OY | blood | S011 | This study |
| Kamisu, Ibaraki, Japan | *Coturnicops exquisitus* | mt, MIG | LC781700 | DRR505892 | Kanto (wintering) | 2018/02/19 | 3E42045 | OY | blood | S012 | This study |
| Kamisu, Ibaraki, Japan | *Coturnicops exquisitus* | MIG | - | DRR505893 | Kanto (wintering) | 2018/02/19 | 3E42046 | OY | blood | S013 | This study |
| Kamisu, Ibaraki, Japan | *Coturnicops exquisitus* | MIG | - | DRR505894 | Kanto (wintering) | 2018/03/12 | 3E42047 | OY | blood | S014 | This study |
| Inashiki, Ibaraki, Japan | *Coturnicops exquisitus* | MIG | - | DRR505895 | Kanto (wintering) | 2018/04/02 | 3E42048 | OY | blood | S015 | This study |
| Inashiki, Ibaraki, Japan | *Coturnicops exquisitus* | MIG | - | DRR505896 | Kanto (wintering) | 2018/04/10 | 3E42049 | OY | blood | S016 | This study |
| Kushiro, Hokkaido, Japan | *Coturnicops exquisitus* | mt, MIG | LC781705 | DRR505897 | Kushiro (breeding) | 2019/08/07 | 3E42068 | OY | blood | S017 | This study |
| Kushiro, Hokkaido, Japan | *Coturnicops exquisitus* | mt, MIG | LC781706 | DRR505898 | Kushiro (breeding) | 2019/08/07 | 3E42076 | OY | blood | S018 | This study |
| Kushiro, Hokkaido, Japan | *Coturnicops exquisitus* | mt, MIG | LC781707 | DRR505899 | Kushiro (breeding) | 2019/08/07 | 3E42077 | OY | blood | S019 | This study |
| Kushiro, Hokkaido, Japan | *Coturnicops exquisitus* | MIG | - | DRR505900 | Kushiro (breeding) | 2019/08/07 | 3E42078 | OY | blood | S020 | This study |

| | | | | | | | | | | | |
|---|---|---|---|---|---|---|---|---|---|---|---|
| Kushiro, Hokkaido, Japan | *Coturnicops exquisitus* | mt, MIG | LC781708 | DRR505901 | Kushiro (breeding) | 2019/08/08 | 3E42079 | OY | blood | S021 | This study |
| Kushiro, Hokkaido, Japan | *Coturnicops exquisitus* | mt, MIG | LC781709 | DRR505902 | Kushiro (breeding) | 2019/08/08 | 3E42080 | OY | blood | S022 | This study |
| Kushiro, Hokkaido, Japan | *Coturnicops exquisitus* | mt, MIG | LC781710 | DRR505903 | Kushiro (breeding) | 2019/08/08 | 3E42081 | OY | blood | S023 | This study |
| Kushiro, Hokkaido, Japan | *Coturnicops exquisitus* | mt, MIG | LC781711 | DRR505904 | Kushiro (breeding) | 2019/08/08 | 3E42083 | OY | blood | S024 | This study |
| Kushiro, Hokkaido, Japan | *Coturnicops exquisitus* | mt, MIG | LC781712 | DRR505905 | Kushiro (breeding) | 2019/08/09 | 3E42084 | OY | blood | S025 | This study |
| Kushiro, Hokkaido, Japan | *Coturnicops exquisitus* | mt, MIG | LC781713 | DRR505906 | Kushiro (breeding) | 2019/08/09 | 3E42085 | OY | blood | S026 | This study |
| Kushiro, Hokkaido, Japan | *Coturnicops exquisitus* | mt, MIG | LC781714 | DRR505907 | Kushiro (breeding) | 2019/08/09 | 3E42086 | OY | blood | S027 | This study |
| Kushiro, Hokkaido, Japan | *Coturnicops exquisitus* | mt, MIG | LC781715 | DRR505908 | Kushiro (breeding) | 2019/08/09 | 3E42087 | OY | blood | S028 | This study |
| Kamisu, Ibaraki, Japan | *Coturnicops exquisitus* | MIG | - | DRR505909 | Kanto (wintering) | 2019/01/26 | 3E42063 | OY | blood | S029 | This study |
| Kamisu, Ibaraki, Japan | *Coturnicops exquisitus* | MIG | - | DRR505910 | Kanto (wintering) | 2019/01/26 | 3E42064 | OY | blood | S030 | This study |
| Kamisu, Ibaraki, Japan | *Coturnicops exquisitus* | MIG | - | DRR505911 | Kanto (wintering) | 2019/01/26 | 3E42065 | OY | blood | S031 | This study |
| Kamisu, Ibaraki, Japan | *Coturnicops exquisitus* | MIG | - | DRR505912 | Kanto (wintering) | 2019/01/26 | 3E42066 | OY | blood | S032 | This study |
| Kamisu, Ibaraki, Japan | *Coturnicops exquisitus* | MIG | - | DRR505913 | Kanto (wintering) | 2019/03/23 | 3E42067 | OY | blood | S033 | This study |
| Hotokenu ma, Aomori, Japan | *Coturnicops exquisitus* | mt, MIG | LC781724 | DRR505914 | Aomori (breeding) | 2017/6/23 | 4C91124 | TM | blood | S034 | This study |
| Tomakom ai, Hokkaido, Japan | *Coturnicops exquisitus* | mt, MIG | LC781723 | DRR505915 | Tomakomai (breeding) | 2019/07/29 | 3H53596 | KM | blood | S035 | This study |
| Tomakom ai, Hokkaido, Japan | *Coturnicops exquisitus* | mt, MIG | LC781716 | DRR505916 | Tomakomai (breeding) | 2019/08/04 | 3H23455 | KM | blood | S036 | This study |
| Tomakom ai, Hokkaido, Japan | *Coturnicops exquisitus* | mt, MIG | LC781717 | DRR505917 | Tomakomai (breeding) | 2019/08/04 | 3H23456 | KM | blood | S037 | This study |
| Tomakom ai, Hokkaido, Japan | *Coturnicops exquisitus* | mt, MIG | LC781718 | DRR505918 | Tomakomai (breeding) | 2019/08/04 | 3H23457 | KM | blood | S038 | This study |
| Tomakom ai, Hokkaido, Japan | *Coturnicops exquisitus* | mt, MIG | LC781719 | DRR505919 | Tomakomai (breeding) | 2019/08/06 | 3H23458 | KM | blood | S039 | This study |
| Tomakom ai, Hokkaido, Japan | *Coturnicops exquisitus* | mt, MIG | LC781720 | DRR505920 | Tomakomai (breeding) | 2019/08/06 | 3H23459 | KM | blood | S040 | This study |
| Tomakom ai, Hokkaido, Japan | *Coturnicops exquisitus* | mt, MIG | LC781721 | DRR505921 | Tomakomai (breeding) | 2019/08/06 | 3H23460 | KM | blood | S041 | This study |
| Tomakom ai, Hokkaido, Japan | *Coturnicops exquisitus* | mt, MIG | LC781722 | DRR505922 | Tomakomai (breeding) | 2019/08/06 | 3H23461 | KM | blood | S042 | This study |
| Muraviovk a, Amur, Russia | *Coturnicops exquisitus* | mt, MIG | LC781725 | DRR505923 | Russia (breeding) | 2017/05/22 | N06529 | HW, WT | swab | S043 | This study |
| Muraviovk a, Amur, Russia | *Coturnicops exquisitus* | mt, MIG | LC781726 | DRR505924 | Russia (breeding) | 2017/05/24 | N06530 | HW, WT | swab | S044 | This study |
| Muraviovk a, Amur, Russia | *Coturnicops exquisitus* | mt, MIG | LC781727 | DRR505925 | Russia (breeding) | 2017/05/24 | N06531 | HW, WT | swab | S045 | This study |

| | | | | | | | | | | | |
|---|---|---|---|---|---|---|---|---|---|---|---|
| Muraviovka, Amur, Russia | *Coturnicops exquisitus* | mt, MIG | LC781728 | DRR505926 | Russia (breeding) | 2017/05/25 | N06532 | HW, WT | swab | S046 | This study |
| Muraviovka, Amur, Russia | *Coturnicops exquisitus* | mt, MIG | LC781729 | DRR505927 | Russia (breeding) | 2017/06/05 | N06533 | HW, WT | swab | S047 | This study |
| Muraviovka, Amur, Russia | *Coturnicops exquisitus* | mt, MIG | LC781730 | DRR505928 | Russia (breeding) | 2017/06/11 | N06534 | HW, WT | swab | S048 | This study |
| Kabansky Zakaznik, Baikal, Russia | *Coturnicops exquisitus* | MIG | - | DRR505929 | Russia (breeding) | 2019/05/31 | KA09927 | BM, HW | feather | S049 | This study |
| Hotokenuma, Aomori, Japan | *Coturnicops exquisitus* | MIG | - | DRR505930 | Aomori (breeding) | 2015/7/15 | 4C91106 | TM | blood | S050 | This study |
| Hotokenuma, Aomori, Japan | *Coturnicops exquisitus* | MIG | - | DRR505931 | Aomori (breeding) | 2015/7/24 | 4C91107 | TM | blood | S051 | This study |
| Hotokenuma, Aomori, Japan | *Coturnicops exquisitus* | MIG | - | DRR505932 | Aomori (breeding) | 2015/7/24 | 4C91108 | TM | blood | S052 | This study |
| Inashiki, Ibaraki, Japan | *Coturnicops exquisitus* | mt | LC781684 | - | Kanto (wintering) | 2018/10/13 | 3E42051 | OY | blood | - | This study |
| Inashiki, Ibaraki, Japan | *Coturnicops exquisitus* | mt | LC781685 | - | Kanto (wintering) | 2018/10/21 | 3E42052 | OY | blood | - | This study |
| Inashiki, Ibaraki, Japan | *Coturnicops exquisitus* | mt | LC781686 | - | Kanto (wintering) | 2018/10/21 | 3E42053 | OY | blood | - | This study |
| Inashiki, Ibaraki, Japan | *Coturnicops exquisitus* | mt | LC781687 | - | Kanto (wintering) | 2018/10/27 | 3E42054 | OY | blood | - | This study |
| Kamisu, Ibaraki, Japan | *Coturnicops exquisitus* | mt | LC781688 | - | Kanto (wintering) | 2018/11/25 | 3E42055 | OY | blood | - | This study |
| Kamisu, Ibaraki, Japan | *Coturnicops exquisitus* | mt | LC781689 | - | Kanto (wintering) | 2018/11/25 | 3E42056 | OY | blood | - | This study |
| Kamisu, Ibaraki, Japan | *Coturnicops exquisitus* | mt | LC781690 | - | Kanto (wintering) | 2018/11/25 | 3E42057 | OY | blood | - | This study |
| Kamisu, Ibaraki, Japan | *Coturnicops exquisitus* | mt | LC781691 | - | Kanto (wintering) | 2018/11/25 | 3E42058 | OY | blood | - | This study |
| Kamisu, Ibaraki, Japan | *Coturnicops exquisitus* | mt | LC781692 | - | Kanto (wintering) | 2018/11/25 | 3E42059 | OY | blood | - | This study |
| Kamisu, Ibaraki, Japan | *Coturnicops exquisitus* | mt | LC781693 | - | Kanto (wintering) | 2018/11/25 | 3E42060 | OY | blood | - | This study |
| Inashiki, Ibaraki, Japan | *Coturnicops exquisitus* | mt | LC781694 | - | Kanto (wintering) | 2018/11/27 | 3E42061 | OY | blood | - | This study |
| Inashiki, Ibaraki, Japan | *Coturnicops exquisitus* | mt | LC781695 | - | Kanto (wintering) | 2018/11/27 | 3E42062 | OY | blood | - | This study |
| Kamisu, Ibaraki, Japan | *Coturnicops exquisitus* | mt | LC781701 | - | Kanto (wintering) | 2019/01/26 | 3E42063 | OY | blood | - | This study |
| Kamisu, Ibaraki, Japan | *Coturnicops exquisitus* | mt | LC781702 | - | Kanto (wintering) | 2019/01/26 | 3E42064 | OY | blood | - | This study |
| Kamisu, Ibaraki, Japan | *Coturnicops exquisitus* | mt | LC781703 | - | Kanto (wintering) | 2019/01/26 | 3E42065 | OY | blood | - | This study |
| Kamisu, Ibaraki, Japan | *Coturnicops exquisitus* | mt | LC781704 | - | Kanto (wintering) | 2019/03/23 | 3E42067 | OY | blood | - | This study |
| Ishikari, Hokkaido, Japan | *Coturnicops exquisitus* | mt | LC781943 | - | Ishikari (migrating) | 2002/04/22 | YI-2002-0813 | YIO | muscle | - | This study |
| Muraviovka, Amur, Russia | *Coturnicops exquisitus* | mt | MG708233 | - | Russia (breeding) | 2016/06/11 | | WT, HW | swab | - | Heim et al. 2019; 10.1017/S0959270918000138 |
| Muraviovka, Amur, Russia | *Coturnicops exquisitus* | mt | MG708234 | - | Russia (breeding) | 2016/06/22 | | WT, HW | swab | - | Heim et al. 2019; 10.1017/S0959270918000138 |
| Muraviovka, Amur, Russia | *Coturnicops exquisitus* | mt | MG708235 | - | Russia (breeding) | 2016/06/26 | | WT, HW | swab | - | Heim et al. 2019; 10.1017/S0 |

| | | | | | | | | | | |
|---|---|---|---|---|---|---|---|---|---|---|
| | | | | | | | | | | 9592709180 00138 |
| Muraviovka, Amur, Russia | *Coturnicops exquisitus* | mt | MG708236 | - | Russia (breeding) | 2016/06/26 | WT, HW | swab | - | Heim et al. 2019; 10.1017/S0959270918000138 |
| | *Coturnicops noveboracensis* | mt | KC614068 | | | | | | | Garcia-R et al. 2014; 10.1016/j.ympev.2014.09.008 |
| | *Coturnicops noveboracensis* | mt | OM992117 | | | | | | | - |
| | *Laterallus jamaicensis* | WGS | CM040151 | SRX14572910 | | 2020/08/03 | | | | PRJNA794174 |
| | *Atlantisia rogersi* | WGS | - | SRX6608087 | | | | | | PRJNA545868 |

* Abbreviations for collectors; ODAYA Yoshiya (OY), TAKAHASHI Masao (TM), HEIM Wieland (HW), KITAZAWA Munehiro (KM), WULF Tom (WT), BASTARDOT Marc (BM), the Yamashina Institute for Ornithology (YIO)

**Table S2** Datasets and the number of sites used for each supplementary analysis

| Supplementary analyses | Linkage | Site type | Out-group | Populations | # Sites | # Samples | Notes |
|---|---|---|---|---|---|---|---|
| **PCAngsd, ngsAdmixture** | unlinked | V | Non | 4 breeding[1] | 1,198 | 30 | - |
| **PCAngsd, ngsAdmixture** | unlinked | V | Laja/ Atro | 4 breeding, 1 wintering[2] | 1,585 | 51 | - |
| **PCAngsd, ngsAdmixture** | unlinked | V | Laja/ Atro | 4 breeding | 1,958 | 32 | - |
| **RAxML** | linked | V/I | Laja | 4 breeding, 1 wintering | 103,130 | 48 | minind = sample size × 0.5 |
| **RAxML** | linked | V | Laja | 4 breeding, 1 wintering | 1,579 | 48 | minind = sample size × 0.5 |
| **RAxML** | linked | V | Laja | 4 breeding | 1,850 | 31 | minind = sample size × 0.5 |
| **RAxML** | linked | V/I | Laja | 4 breeding, 1 wintering | 97,563 | 48 | minind = sample size × 0.8 |
| **RAxML** | linked | V/I | Laja | 4 breeding | 101,345 | 31 | minind = sample size × 0.8 |
| **RAxML** | linked | V | Laja | 4 breeding, 1 wintering | 1,503 | 48 | minind = sample size × 0.8 |
| **RAxML** | linked | V | Laja | 4 breeding | 1,642 | 31 | minind = sample size × 0.8 |
| **$F_{ST}$** | Unlinked | V | Non | 4 breeding | 1,198 | 30 | - |

Abbreviation: V, variants; V/I, both variants and invariants; Laja, *Laterallus jamaicensis*; Atro, *Atlantisia rogersi*; 1, Tomakomai, Kushiro, Russia, Aomori; 2, Kanto; 3, Tomakomai, Kushiro, Continental.

**Table S3** Coefficients estimated for each comparison of the summary indices between the island and continental populations

| Response variable | Explanatory variable | Base population | Compared population | Estimate | Std. Error | t value | Pr(>\|t\|) |
|---|---|---|---|---|---|---|---|
| $\theta_w$ | Intercept | - | - | 0.010 | 0.003 | 3.666 | <0.001 |
| | Population | Island | Continent | 5.75E-04 | 3.36E-04 | 1.709 | 0.089 |
| | log(contig length) | - | - | -2.69E-04 | 1.73E-04 | -1.553 | 0.122 |
| Tajima's *D* | Intercept | - | - | 0.817 | 0.721 | 1.132 | 0.259 |
| | Population | Island | Continent | -0.239 | 0.088 | -2.719 | 0.007 |
| | log(contig length) | - | - | -0.087 | 0.045 | -1.924 | 0.056 |

**Table S4** Model parameters and details about model evaluation on the best runs of each model. See Fig. 3 and Table S5 for parameter names

| Model | Best run | NPOP0 | NPOP1 | NANC0 | NANC1 | NANCALL | N0M01 | N1M10 | NANC0M01 | NANC1M10 | $t_{MIG}$ | k | MaxObsLhood-MaxEstLhood | AIC | dAIC | Akaike model weight |
|---|---|---|---|---|---|---|---|---|---|---|---|---|---|---|---|---|
| **Different migration** | run97 | 92,154 | 6,885 | 16612 | 1697236 | 735276 | 3.3 | 8.9 | 1465.3 | 0.6 | 14205 | 10 | 326.39 | 32379.80 | 0 | 0.747 |
| **Constant migration** | run2 | 116,172 | 16,172 | 15561 | 366910 | 759708 | 3.1 | 6.1 | - | - | 18831 | 8 | 327.73 | 32381.97 | 2.17 | 0.253 |
| **Migration** | run81 | 4,246,942 | 313,641 | 2266898 | 376643 | 557872 | 3.5 | 12.1 | - | - | 123563 | 8 | 330.62 | 32395.26 | 15.46 | 3.29E-04 |
| **Past migration** | run12 | 209,538 | 236,942 | 3551530 | 289272 | 517234 | - | - | 2.8 | 0.8 | 1337 | 8 | 332.60 | 32404.40 | 24.59 | 3.41E-06 |
| **No migration** | run34 | 4,966,816 | 4,979,830 | 1682 | 2422774 | 4761341 | - | - | - | - | 2613337 | 6 | 429.07 | 32844.68 | 464.87 | 8.46E-102 |

**Table S5** Parameter settings for the fastsimcoal2 analyses

| Integer or decimal | Parameter name | Distribution | Minimum | Maximum | Models that included the parameter*** |
|---|---|---|---|---|---|
| Integer | N[CONT] | logunif | 1000 | 10000000 | All |
| Integer | N[ISL] | logunif | 1000 | 10000000 | All |
| Integer | NANC[ALL] | logunif | 1000 | 10000000 | All |
| Integer | NANC[CONT] | logunif | 1000 | 10000000 | All |
| Integer | NANC[ISL] | logunif | 1000 | 10000000 | All |
| Decimal | N0M01* | logunif | 1.00E-02 | 100000 | Recent, Constant, Different |
| Decimal | N1M10* | logunif | 1.00E-02 | 100000 | Recent, Constant, Different |
| Decimal | NANC0M01* | logunif | 1.00E-02 | 100000 | Ancient, Different |
| Decimal | NANC1M10* | logunif | 1.00E-02 | 100000 | Ancient, Different |
| Decimal | TPROP** | logunif | 0.002 | 1 | All |

*0 and 1 indicate the continent and island, respectively, and N*x*M*xy* or NANC*x*M*xy* (where 0 or 1 replaces italicized letters) indicates a migration matrix where the number of migrants that *x* receives from *y* population. This notation with "ANC" indicates the migration matrix before $t_{MIG}$.

**TPROP is a value that is translated to the time of demographic changes for both the island and continental populations ($t_{MIG}$) by multiplying it by 50,000. ***If "All" is indicated, the parameter was included in all five models tested; otherwise, a specific model name was provided. Refer to the Main Text and Fig. 3 for other parameters.

**Table S6** Sources of occurrence records for the species distribution model

| Sources | Seasons | Number of records | Country | Note |
|---|---|---|---|---|
| Anisimova et al. (2019) Baikal Zoological Journal | breeding | 1 | Russia | |
| Fukuda et al. (2019) | wintering | 9 | Japan | |
| Heim et al. (2019) | breeding | 9 | Russia | |
| Matsumiya & Numano (2022) | wintering | 12 | Japan | |
| Mlíkovský et al. (2024) | breeding, autumn migration, spring migration, wintering, unknown | 88 | Russia, Mongolia, China, Japan, North Korea, South Korea | This is a review paper that summarises records collected from multiple sources. Refer to this publication for their original sources. |
| Sadakuni et al. (2018) | breeding | 20 | Japan | |
| Takahashi et al. (2018) | wintering | 17 | Japan | |
| Sampling site | breeding | 6 | Japan, Russia | The sampling sites used in our analyses were added to this dataset to ensure that they are included in the analysis |
| Senzaki (personal communication) | autumn migration | 1 | Japan | |

### Supplementary Figures

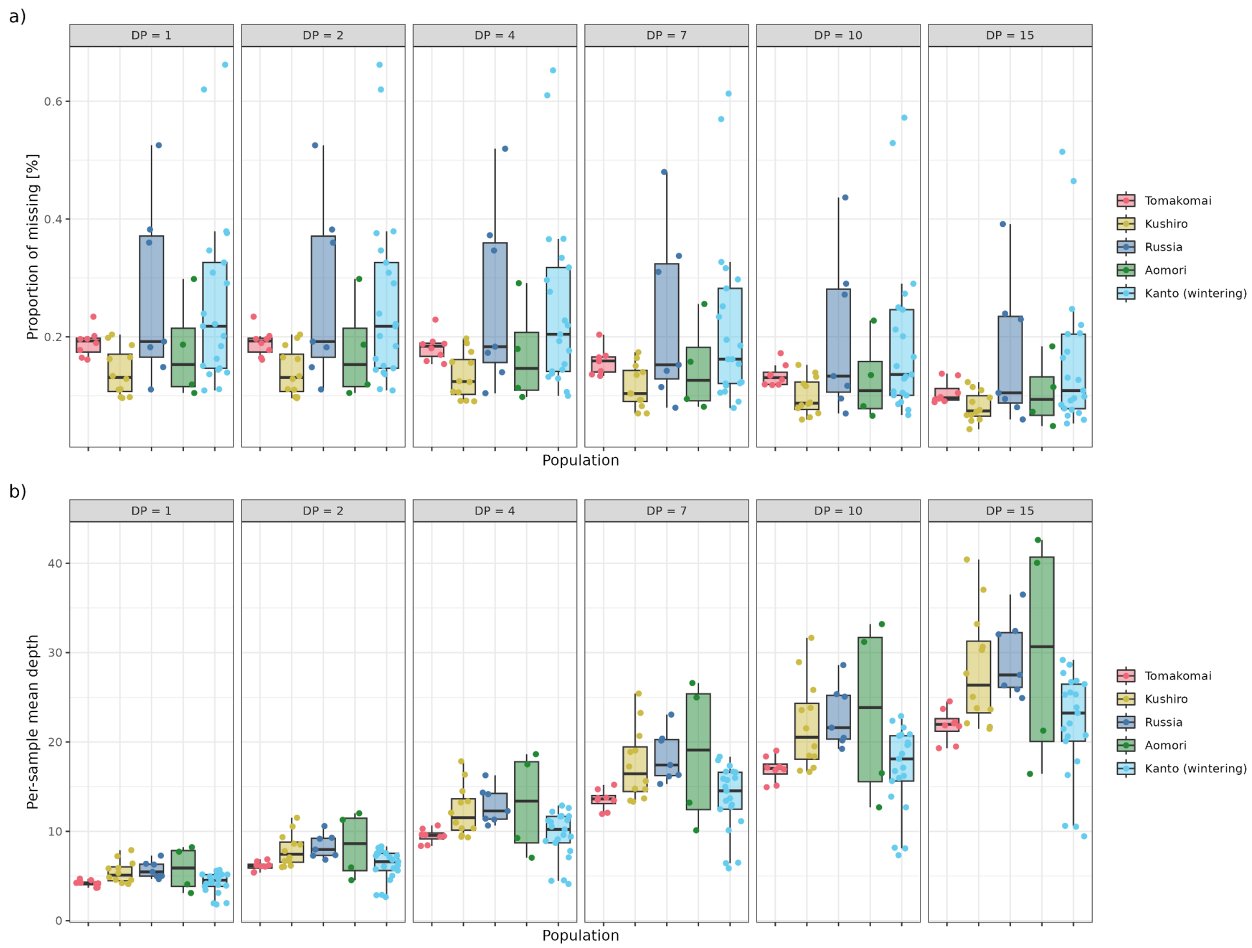


**Figure S1** Inter-population comparisons of (a) per-sample proportion missingness and (b) per-sample mean depth under different minimum depth conditions (DP = 1 to 15) for single nucleotide polymorphisms (SNPs) called in the pre-analyses. SNPs were called by BCFtools with a SNP *p*-value of 1e-06 ('-p' flag). No filter for minor allele frequency (MAF) was used for these analyses.

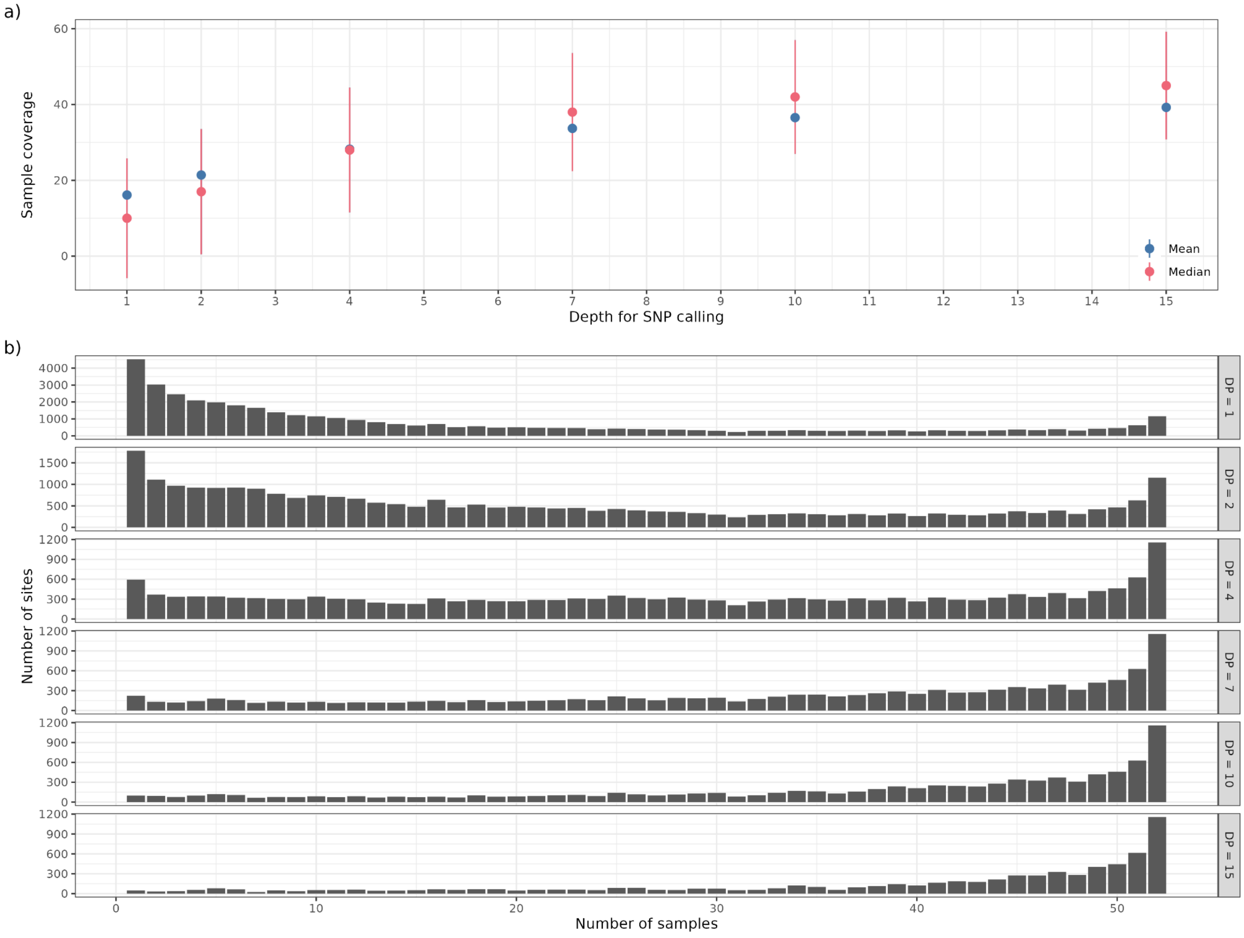


**Figure S2** The distribution of sample coverage by called SNPs using different minimum depth filters in the analyses. a) A plot shows mean and median numbers of sample coverage with error bars computed by ggplot2 v. 3.5.0. b) The number of called SNPs is plotted against the number of samples covered by them. The same SNPs as Figure S1 were used.

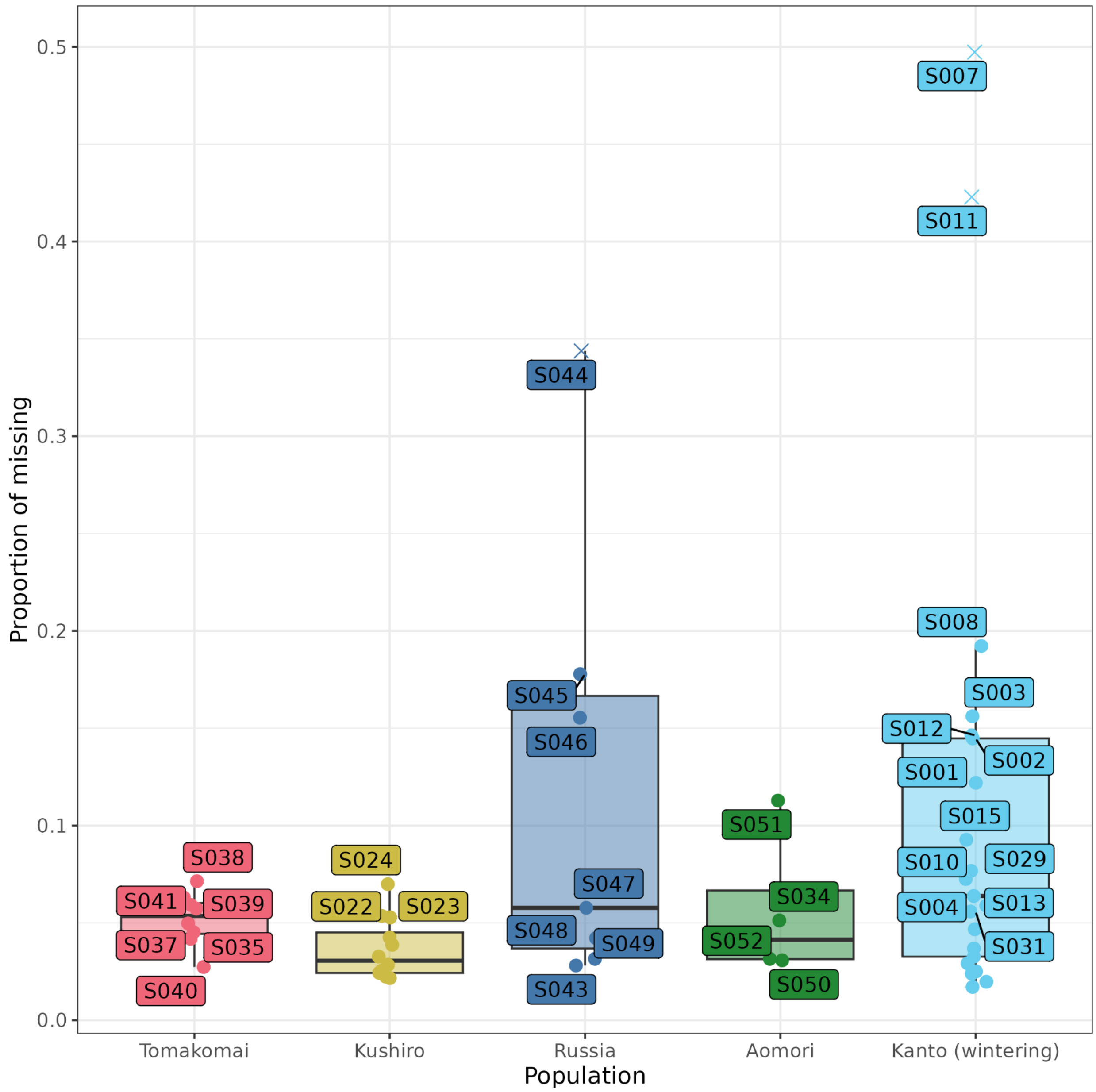


**Figure S3** Per-sample proportions of Ns (missing values) in the called SNPs obtained from the ANGSD pre-analysis were compared among different populations. Crosses indicate samples exceeding missing proportion >0.3 (i.e., 30%) in the pre-analysis that were removed from the downstream analyses.

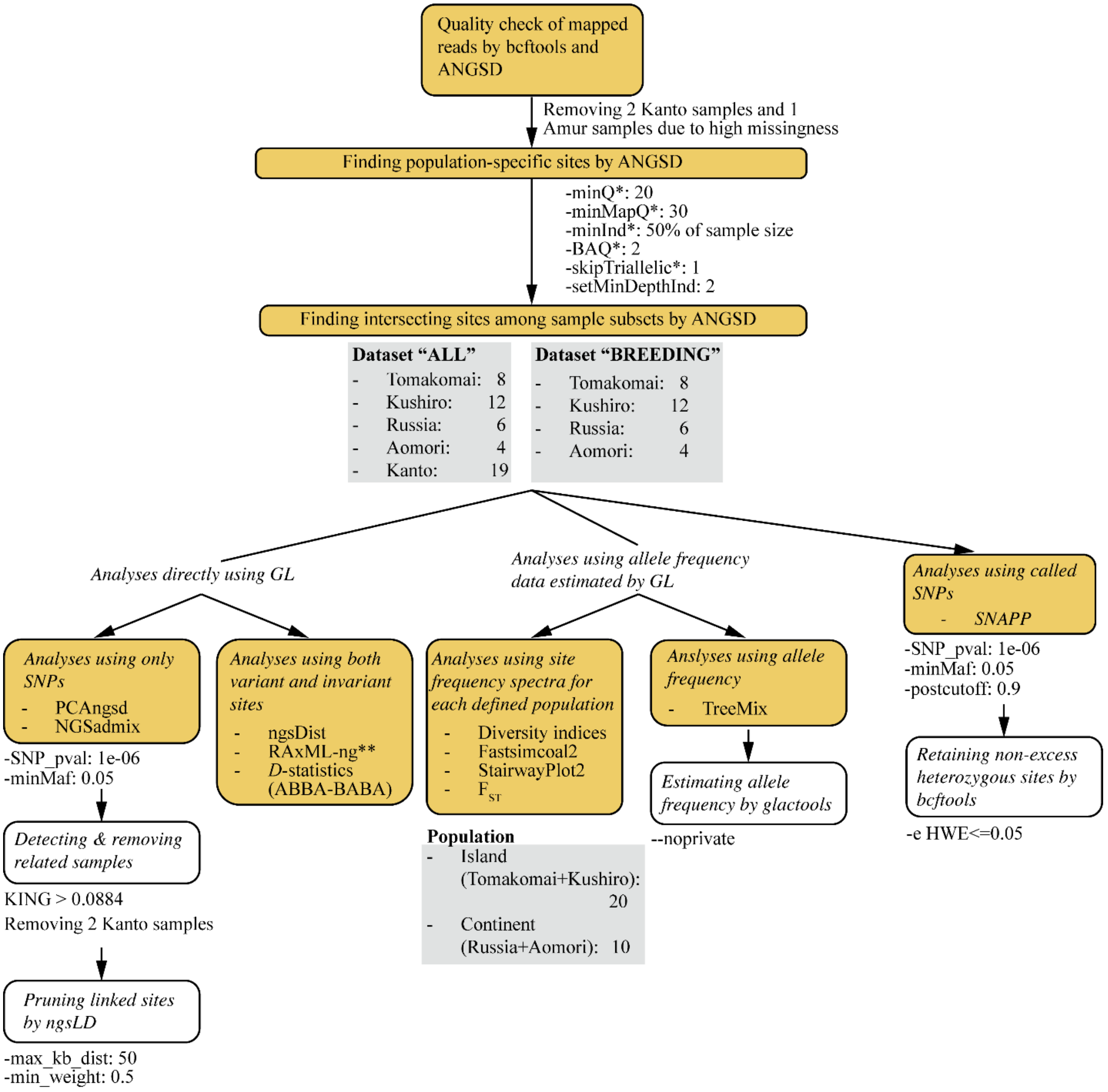


**Figure S4** A summary of the analysis flow using genotype likelihoods (GLs). Filtering parameters for each analysis are listed beneath the boxes. Yellow boxes indicate ANGSD analyses, white boxes indicate supplementary analyses for filtering, and grey boxes indicate datasets used for different analyses. Symbols: *Parameters that were used in all the ANGSD analyses; **Supplementary RAxML analyses were done using different datasets with different filtering parameters (see Supplementary Methods).

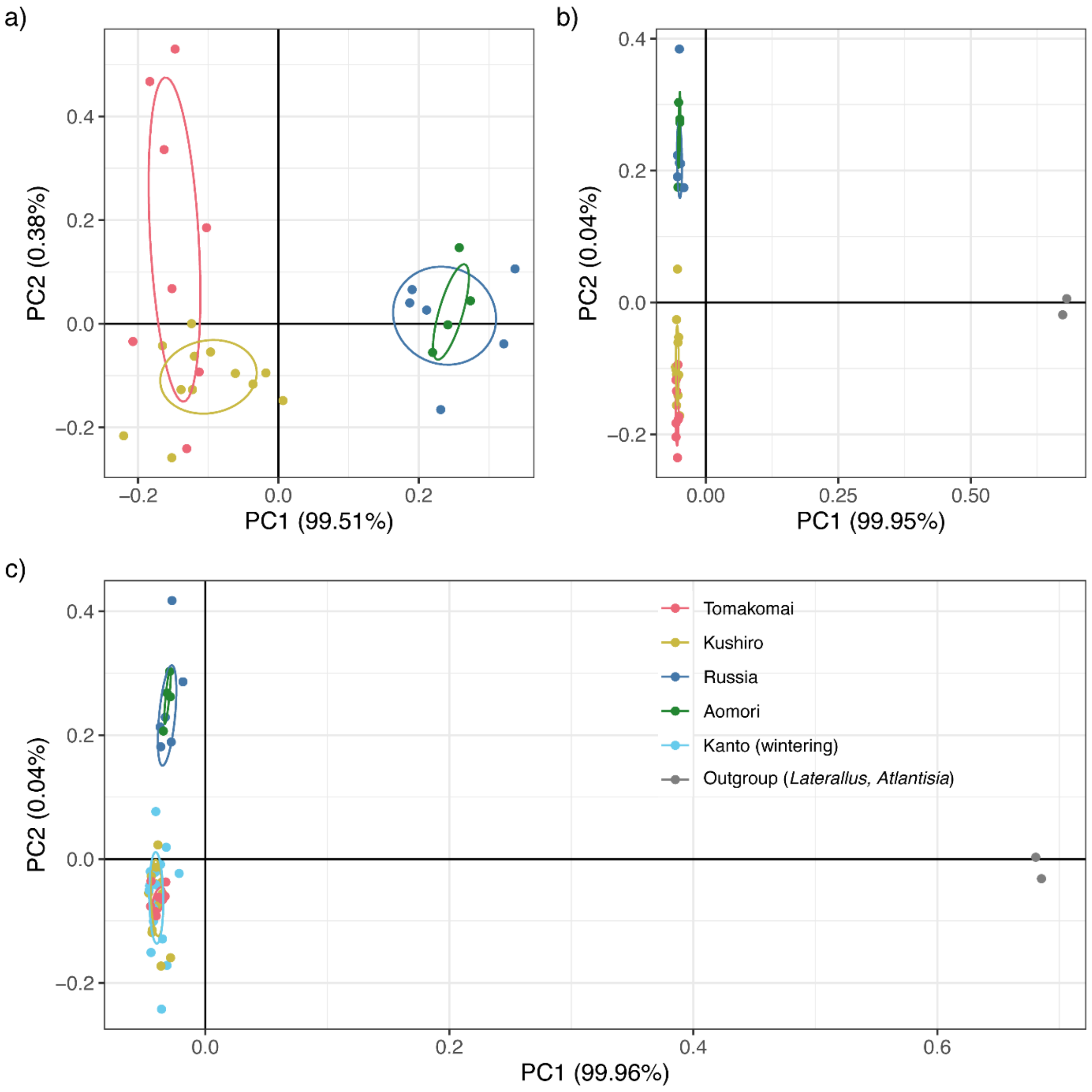


**Figure S5** Results of principal component analysis (PCA) computed by PCAngsd for different datasets, consisting of samples a) from breeding populations only, b) from breeding populations with two outgroup species and c) breeding and wintering populations as well as two outgroup species

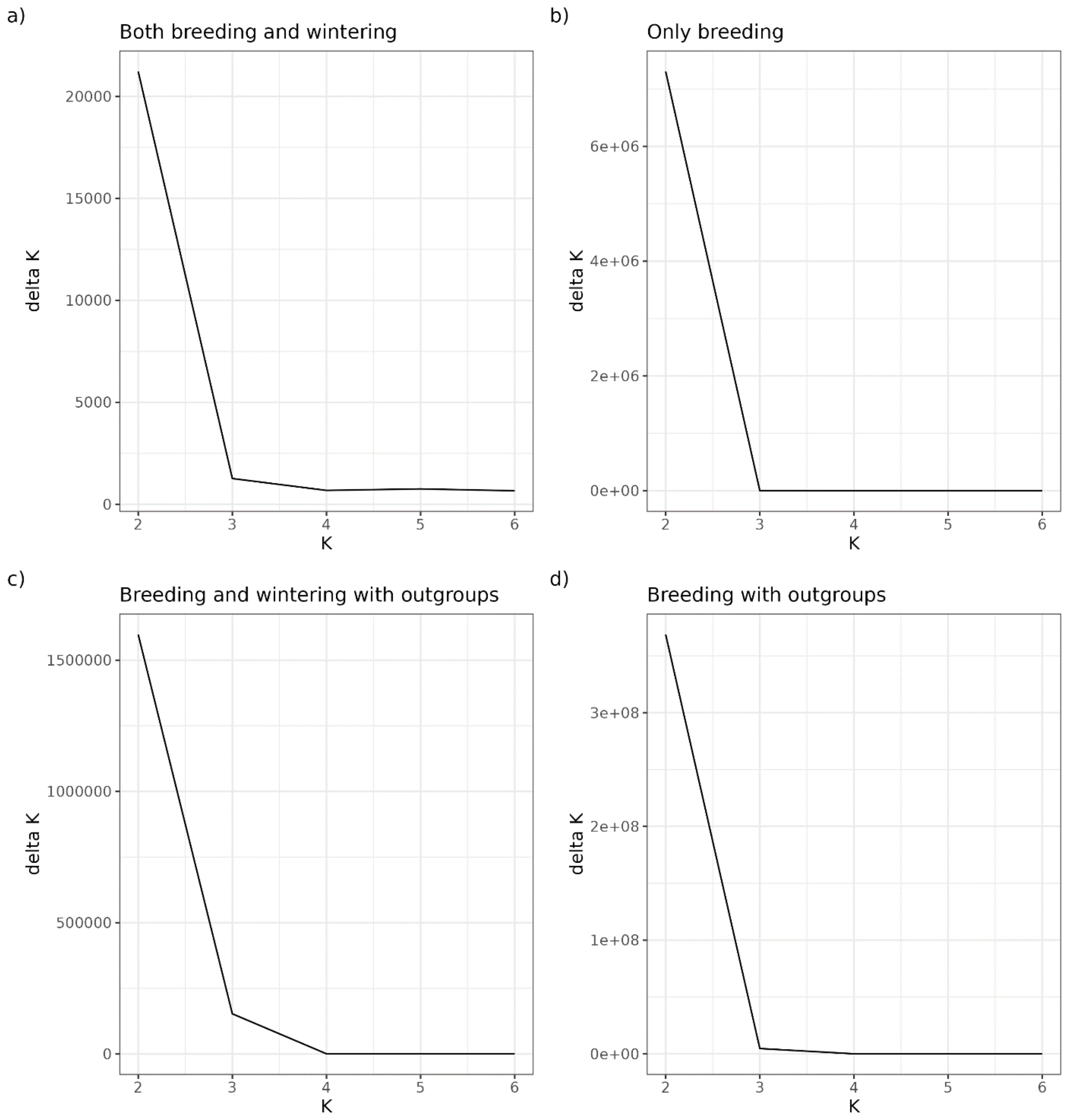


**Figure S6** The ΔK values calculated for each K in different ADMIXTURE analyses using different datasets.

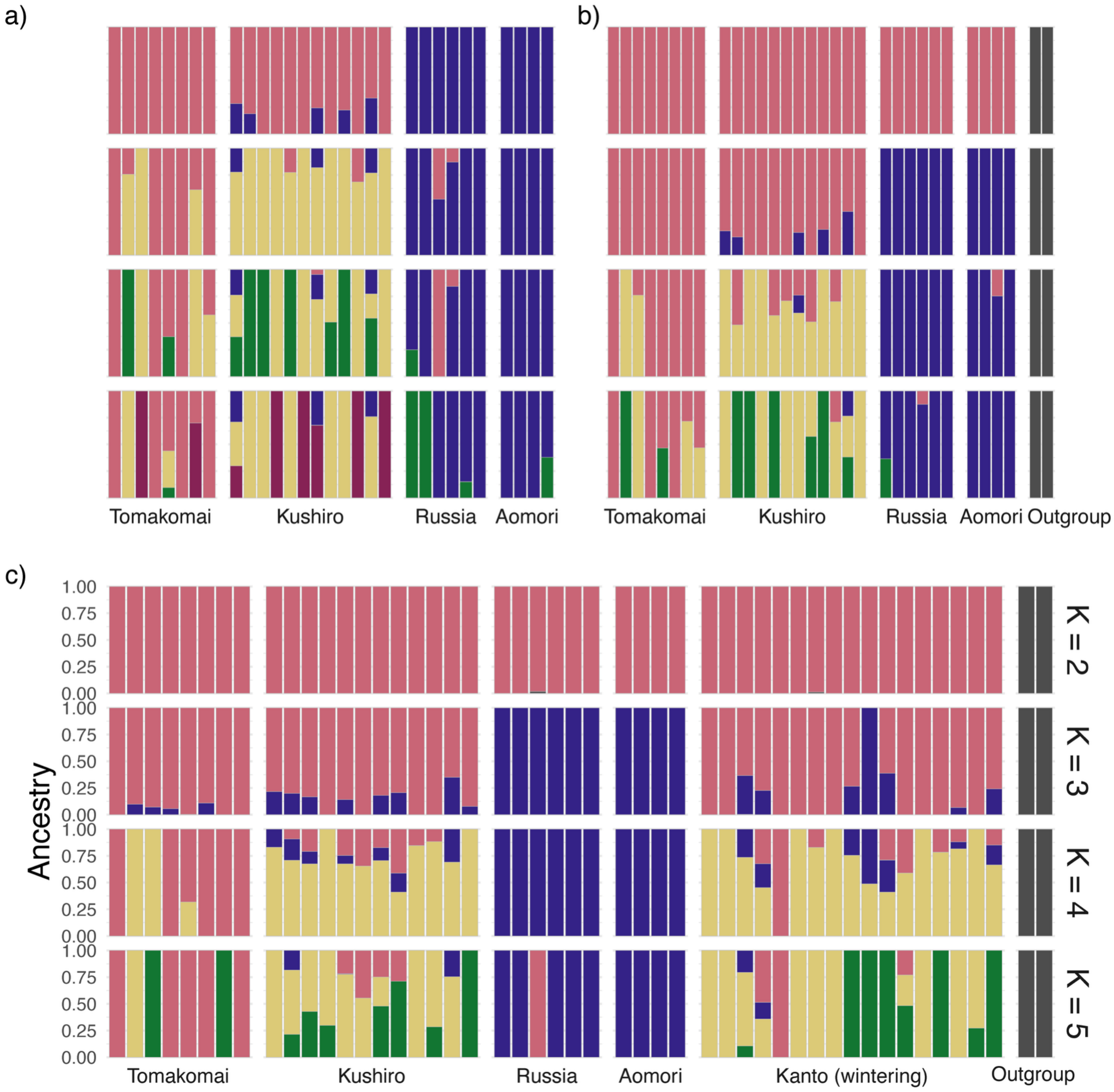


**Figure S7** Results of ADMIXTURE analyses using datasets consisting of samples from a) only breeding populations, b) breeding populations with two outgroup species, and c) breeding and wintering populations with two outgroup species

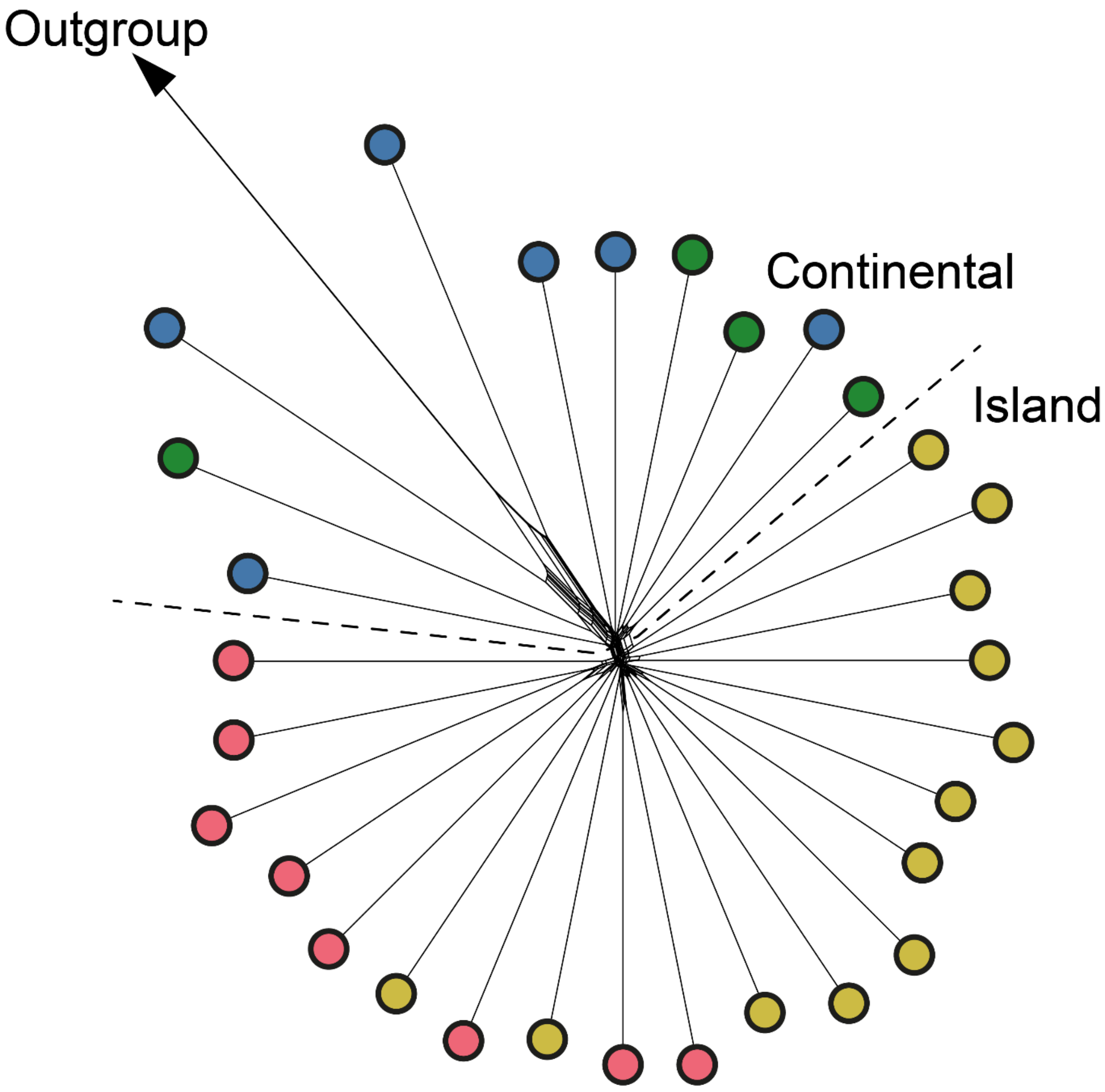


**Figure S8** A NeighborNet plot constructed based on the pairwise genetic distance among samples from breeding populations calculated by ngsDist. The dotted line indicates where the continental and island clades are separated. The outgroup samples (*Laterallus jamaicensis* and *Atlantisia rogersi*) were genetically distant from the ingroup and not shown in this figure.

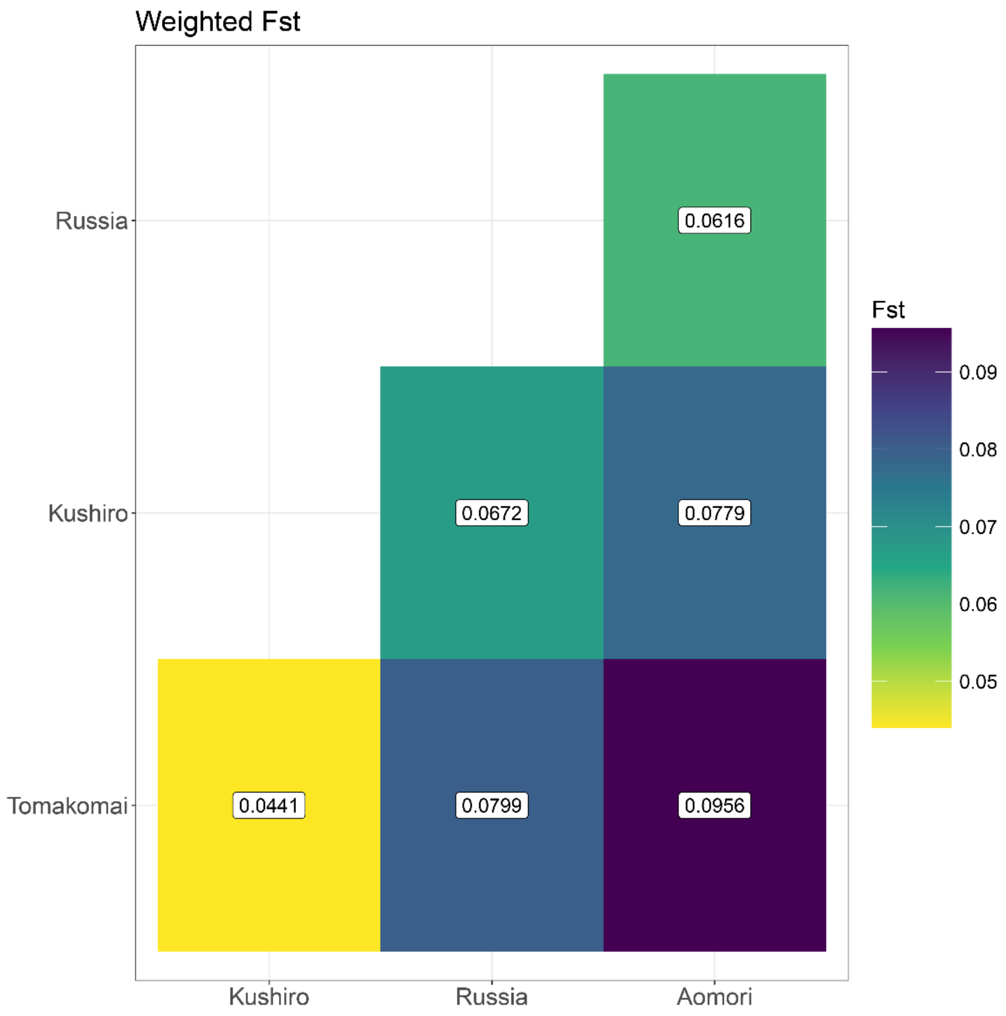


**Figures S9** Weighted $F_{ST}$ calculated among the four breeding populations using unlinked two-dimensional site frequency spectra. $F_{ST}$ increases from yellow, green, to indigo.

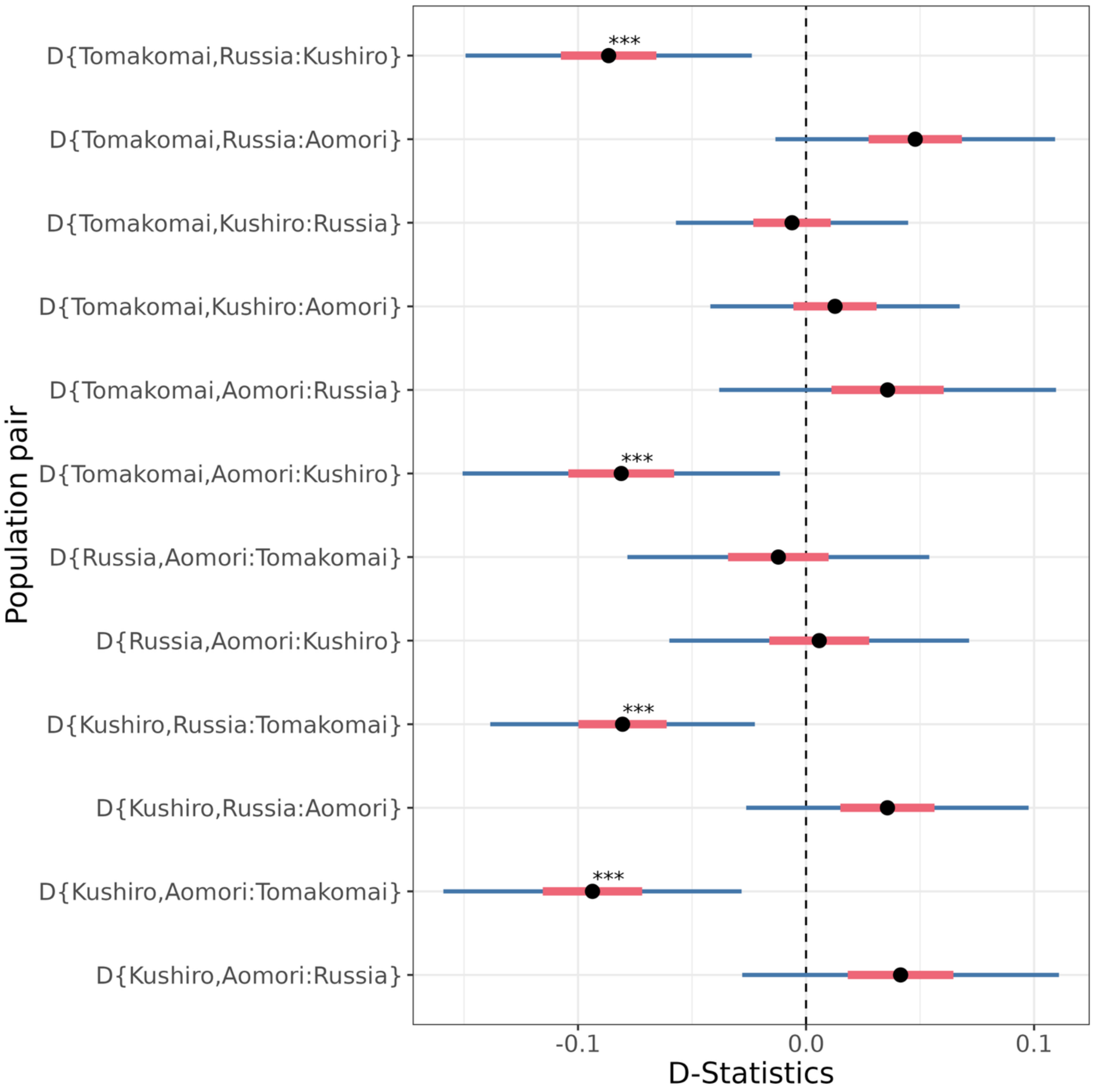


**Figure S10** D-statistic calculated for each trio of four breeding populations with *Laterallus jamaicensis* as an outgroup. The tree topology tested (((H1, H2)H3)Outgroup) is presented in a format D{H1,H2:H3} on the y-axis. For each D-statistic, the standard deviation (red) and the threshold of $|Z| = 3$ (blue) are shown. Those that do not cross 0 indicate that values are statistically significant (denoted by $^{***}$).

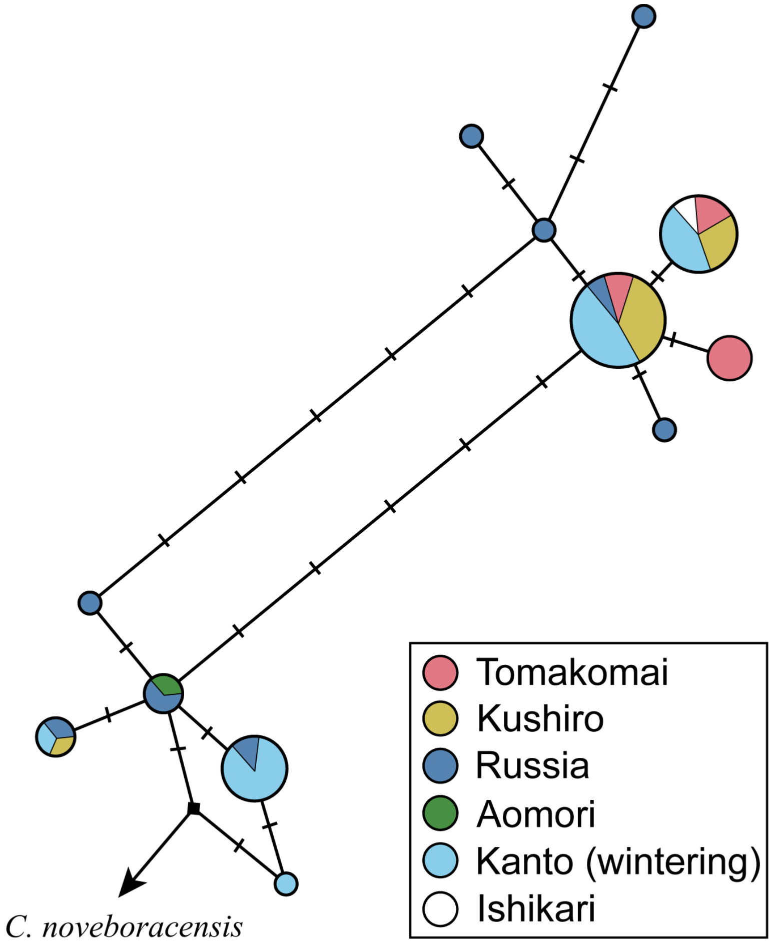


**Figure S11** A haplotype network reconstructed for the partial mitochondrial *Cytb* region (772 bp). The color of the pies corresponds to the sampling locality. The size of the circles reflects the number of samples representing the haplotype.

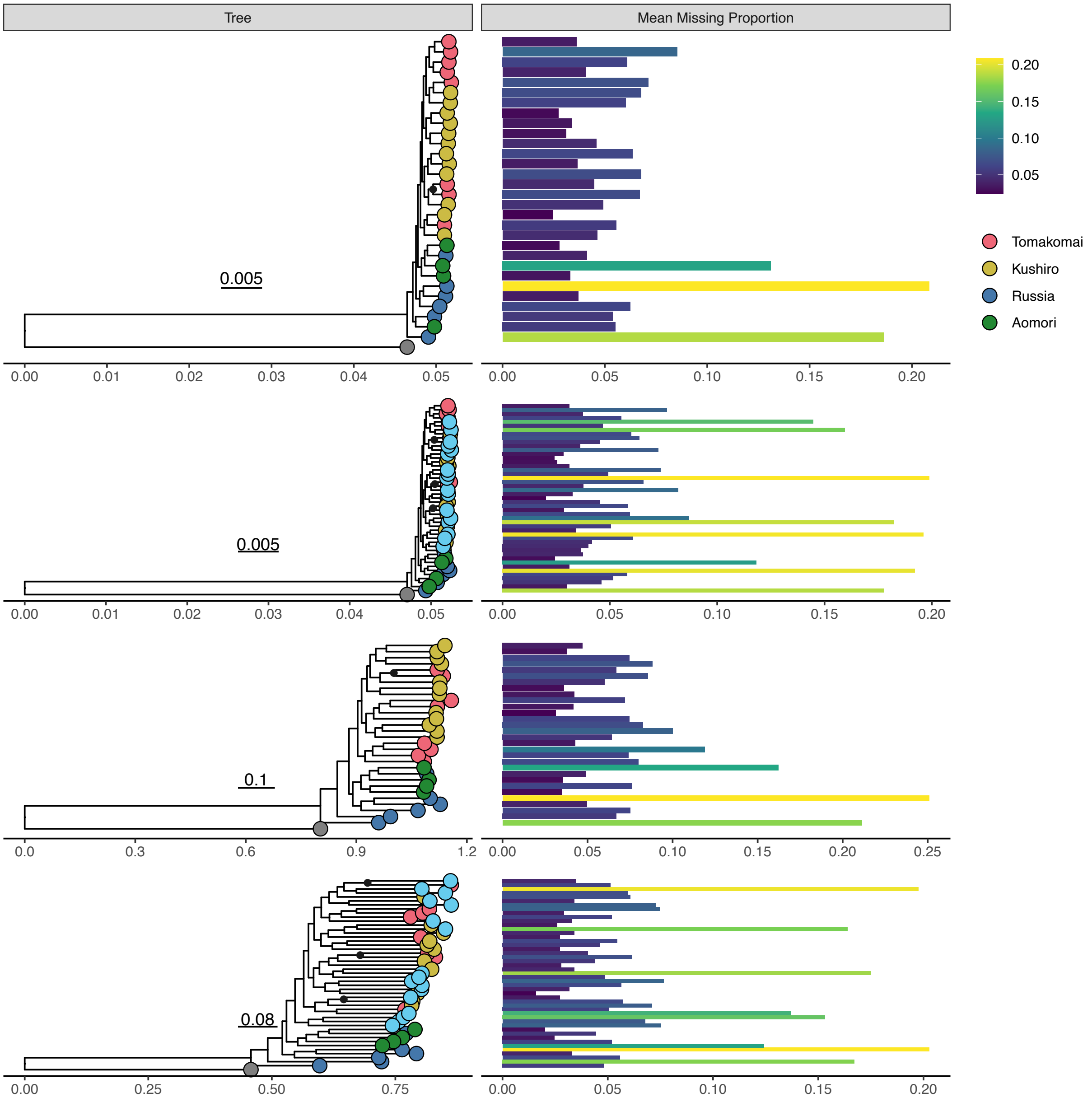


**Figure S12** Maximum likelihood phylogenetic trees reconstructed by RAxML using sites that covered at least 50% of the total sample size, based on GLs calculated for different datasets, a) both variant and invariant sites from only breeding populations, b) both variant and invariant sites from both breeding and wintering populations, c) only variant sites from both breeding and wintering populations, and d) only variant sites from breeding populations. Black circles indicate nodes with bootstrap values >70. The right panels indicate the proportion of missing values (Ns) for each sample.

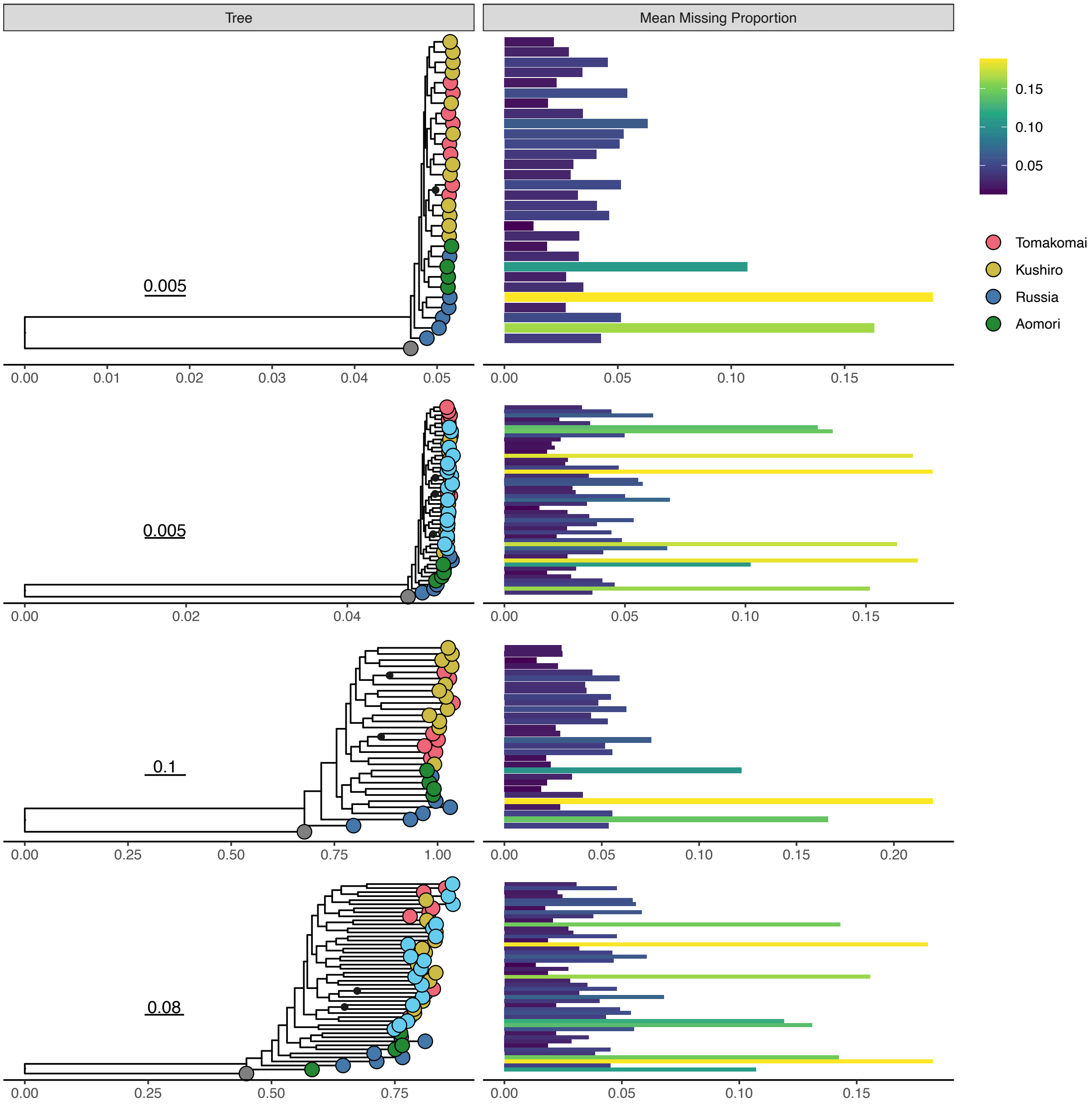


**Figure S13** Maximum likelihood phylogenetic trees reconstructed by RAxML using sites that covered at least 80% of the total sample size, based on GLs calculated for different datasets. Labels and plot annotations are the same as in Figure S9.

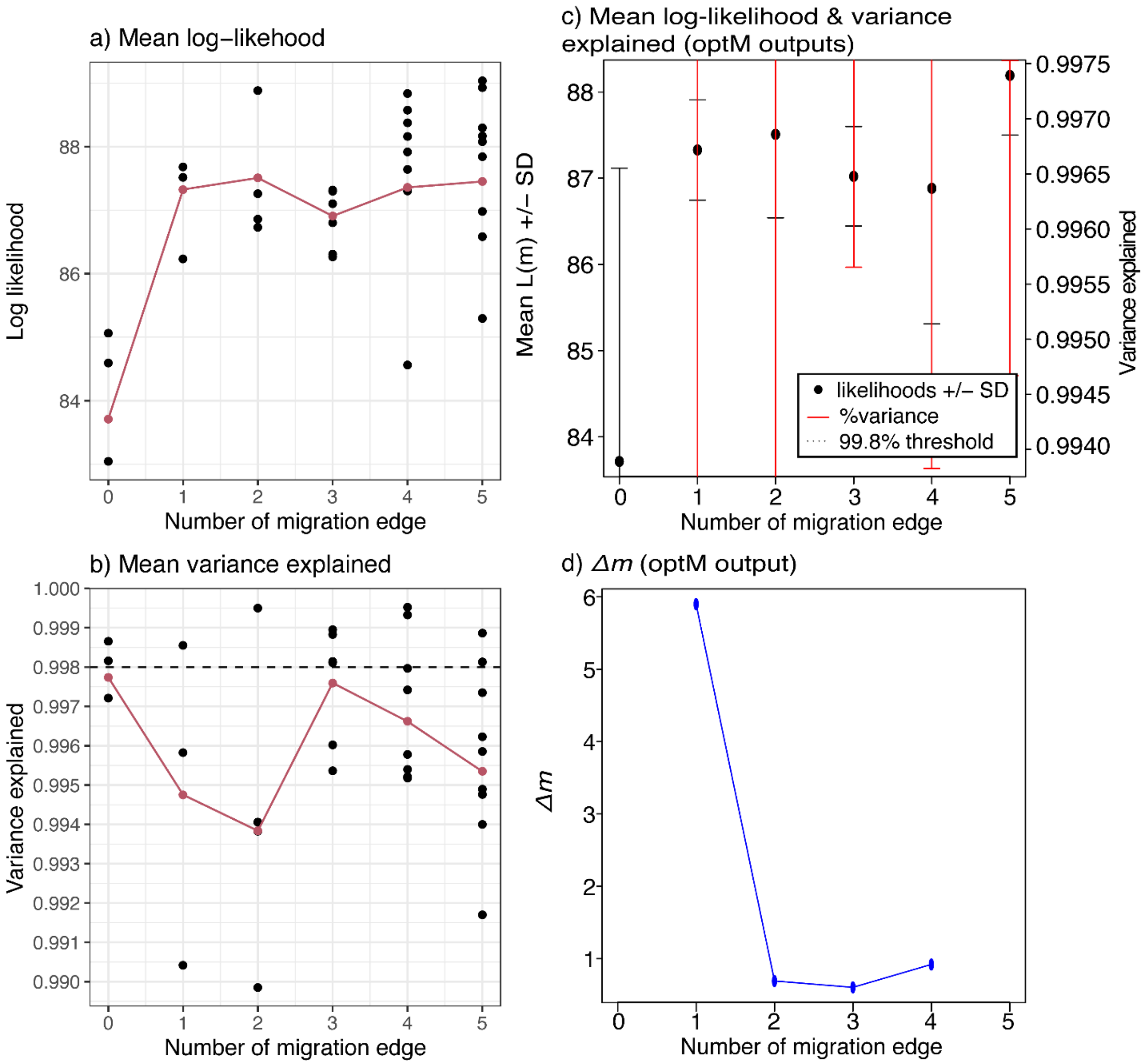


**Figure S14** Metrics for the evaluation of model performance of population-level TreeMix runs with different numbers of migration edges (m = 0 to 5). a) Model likelihoods and b) the mean variance explained for the population relatedness by the models were plotted against different numbers of migration edges. Each black dot shows the values of each run, and the red lines summarize their mean values. (c) A similar plot was generated by the R package 'OptM', which shows almost the same data except for the clipping of large values. (d) $\Delta m$ values for each migration edge calculated by OptM. Note that $\Delta m$ cannot be computed for runs with the lowest (m = 0) and highest (m = 5) migration edges.

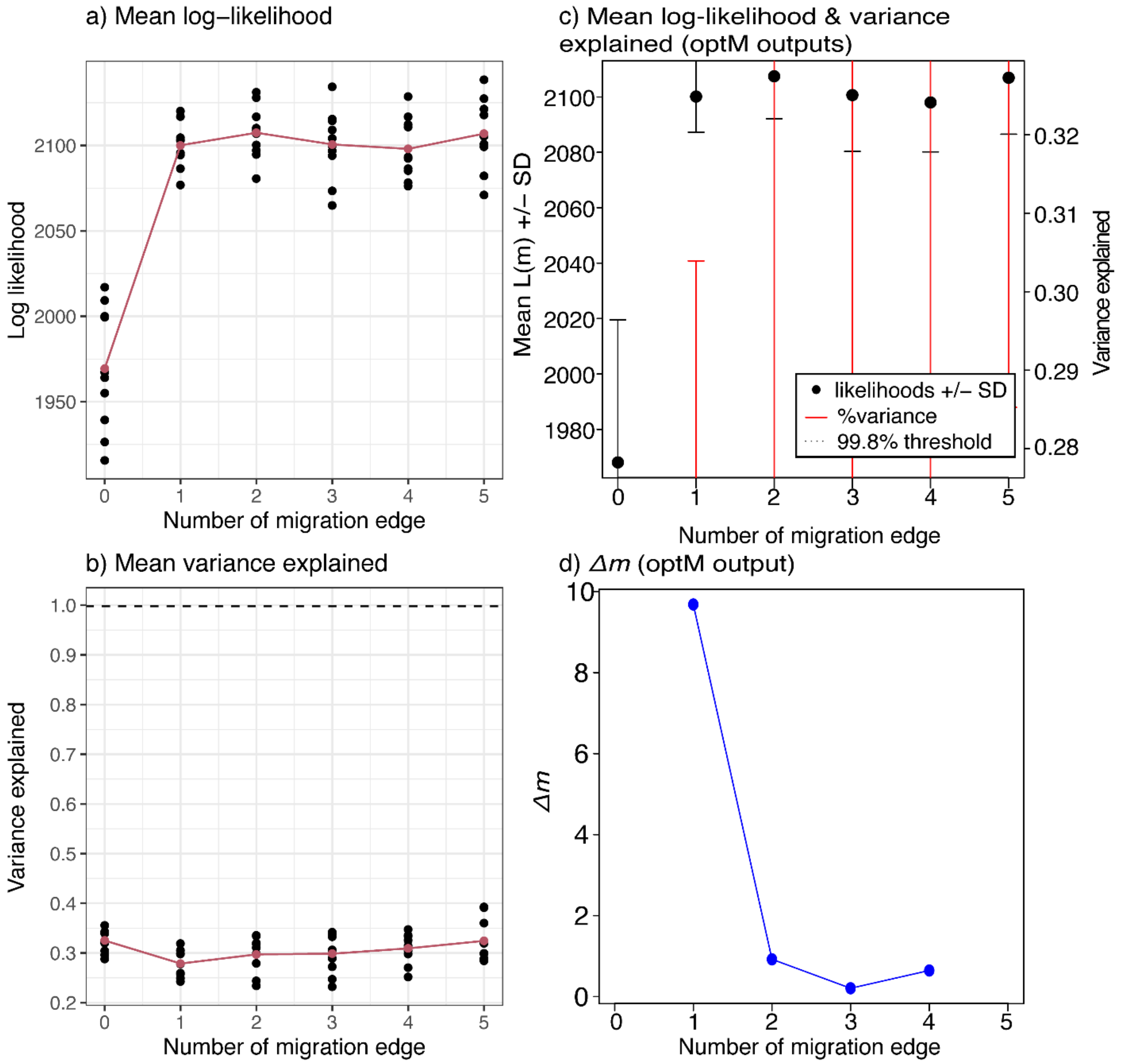


**Figure S15** Metrics for the evaluation of model performance of individual-level TreeMix runs with different numbers of migration edges (m = 0 to 5). See Figure S14 for details

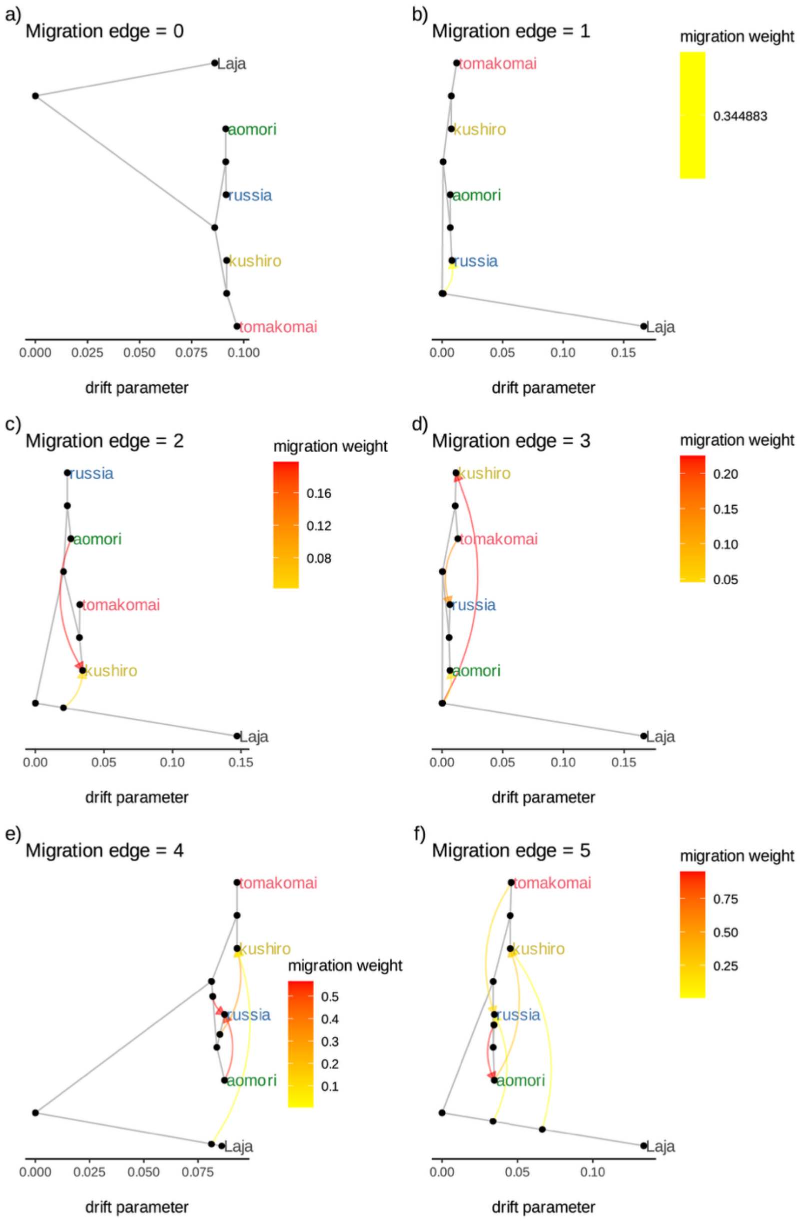


**Figure S16** The best-fit model of TreeMix runs for each migration edge from 0 to 5, which scored the highest log-likelihood values among models with the same migration edge (see Fig. S17 for their log-likelihood values). Colored arrows show possible migration between populations, and different colors indicate the intensity of such events. Laja is the abbreviation for *Laterallus jamaicensis* (outgroup).

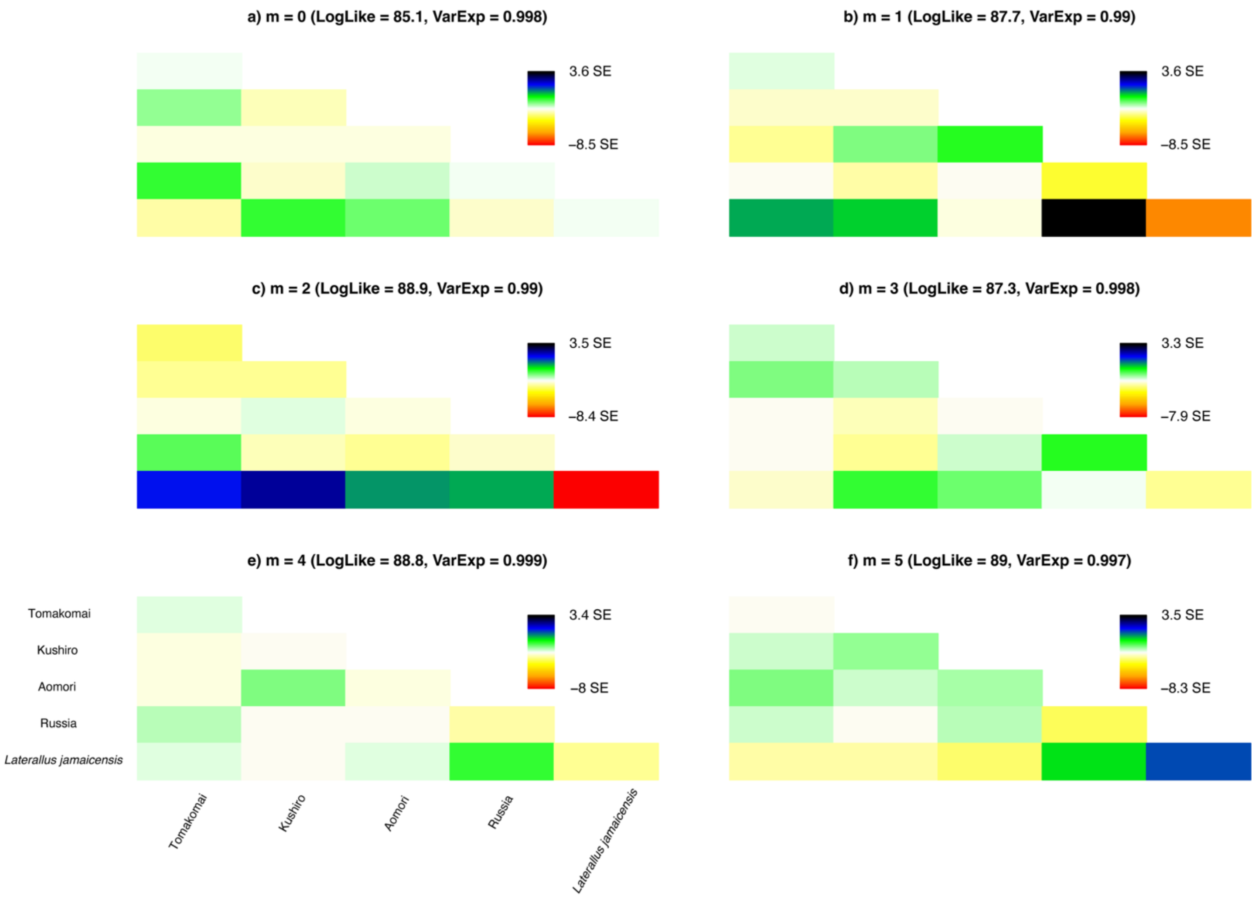


**Figure S17** The residual fit of variance explained for each pair of populations in the best-fit models for each migration edge. Paler colors indicate lower residuals. Positive values represent populations that are more closely related to each other in the data than in the best-fit tree.

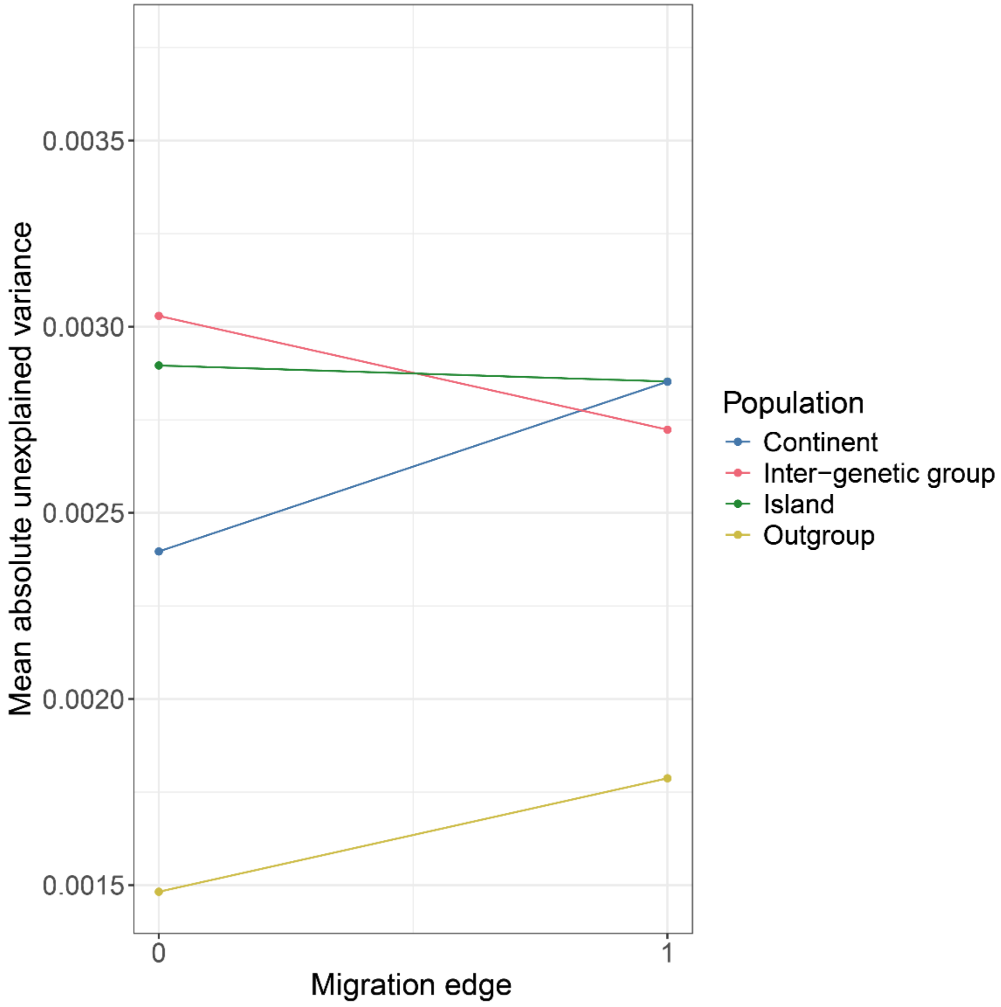


**Figure S18** The change in mean absolute unexplained variance of the best-performed individual-based TreeMix outputs between migration edge = 0 and 1. Higher values indicate higher unexplained variance. Colors indicate groups of samples to calculate the mean values (Continent, within-continent variance [blue]; Island, within-island variance [green]; Inter-genetic group, between-continent-island variance [red]; Outgroup, between-outgroup-ingroup variance [yellow]).

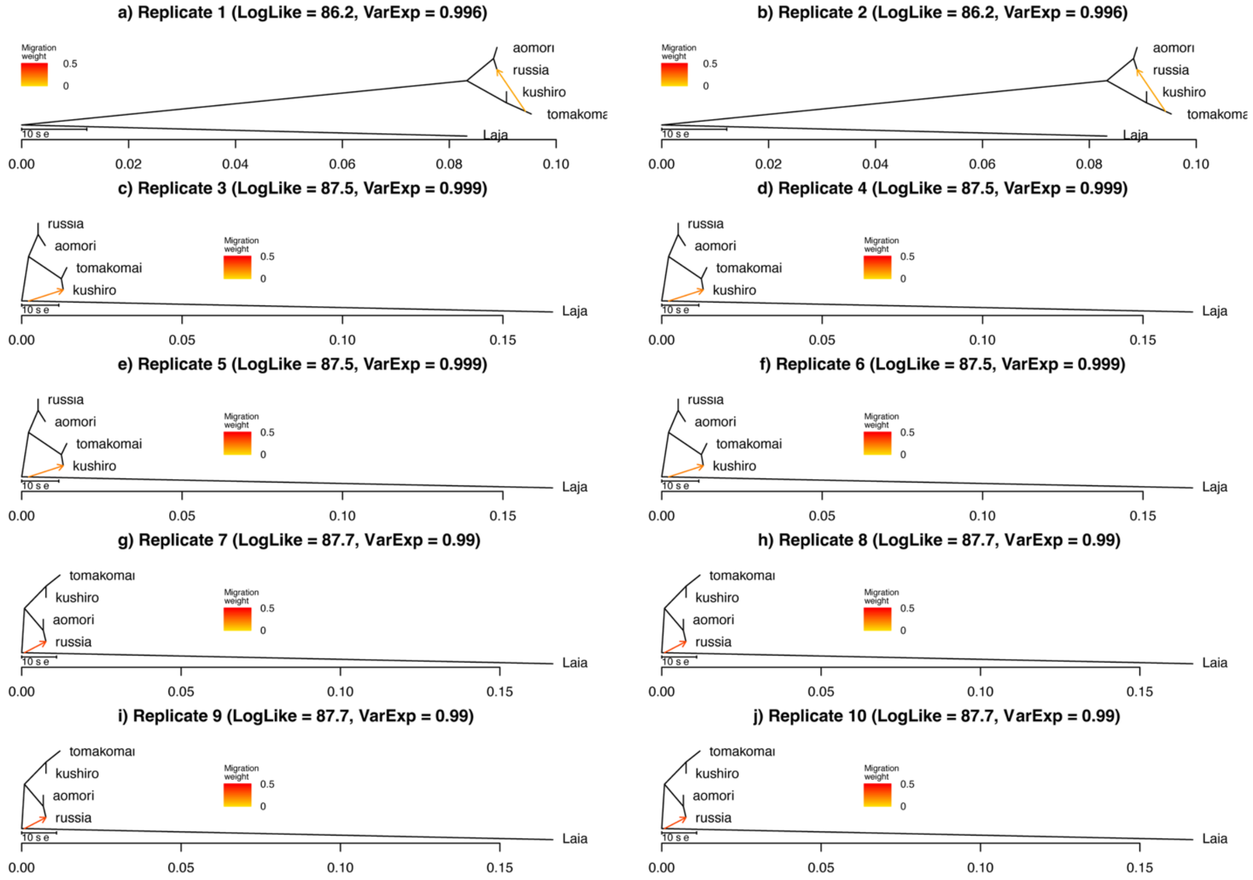


**Figure S19** Results of 10 replicates of the TreeMix model with the migration edge = 1.

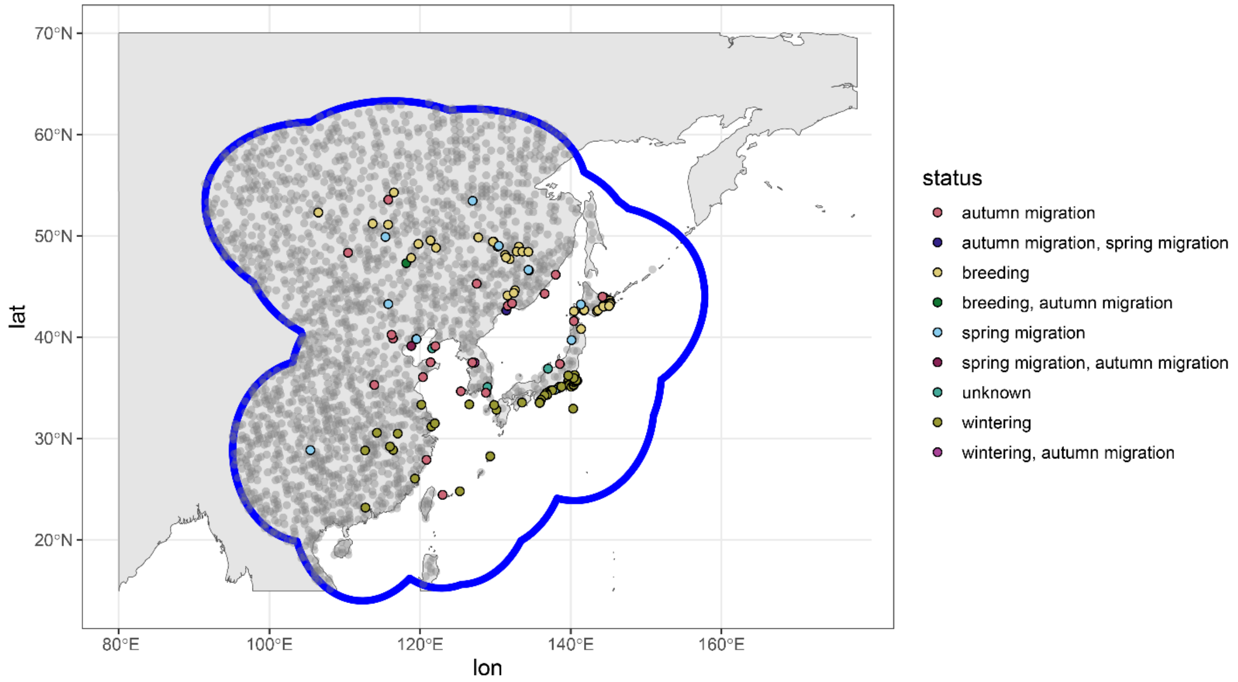


**Figure S20** Background points (transparent gray) and their sampling extent (blue). The extent for the background point collection was determined by the 1,000-km buffered polygon of the convex hull of all the occurrence records, including both breeding and non-breeding.

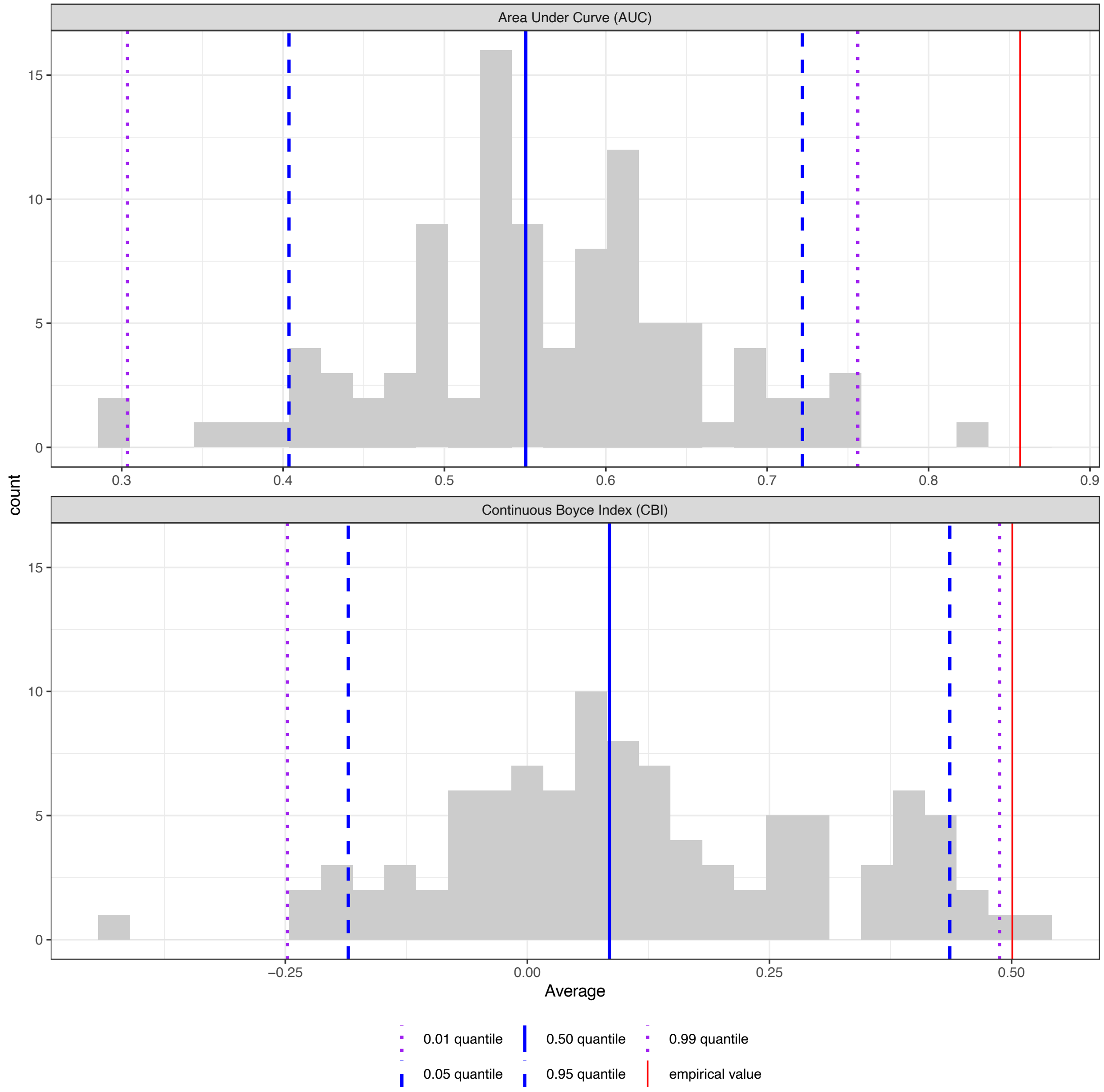


**Figure S21** Evaluation of the constructed distribution model by comparisons with null models based on the Area Under the Curve (AUC) and Continuous Boyce Index (CBI). The gray bars indicate the calculated indices for null models using random occurrence records with the same model settings as the real model. The red lines indicate the values for the indices of the real model, and blue and purple lines indicate quantiles of the indices for the null models.

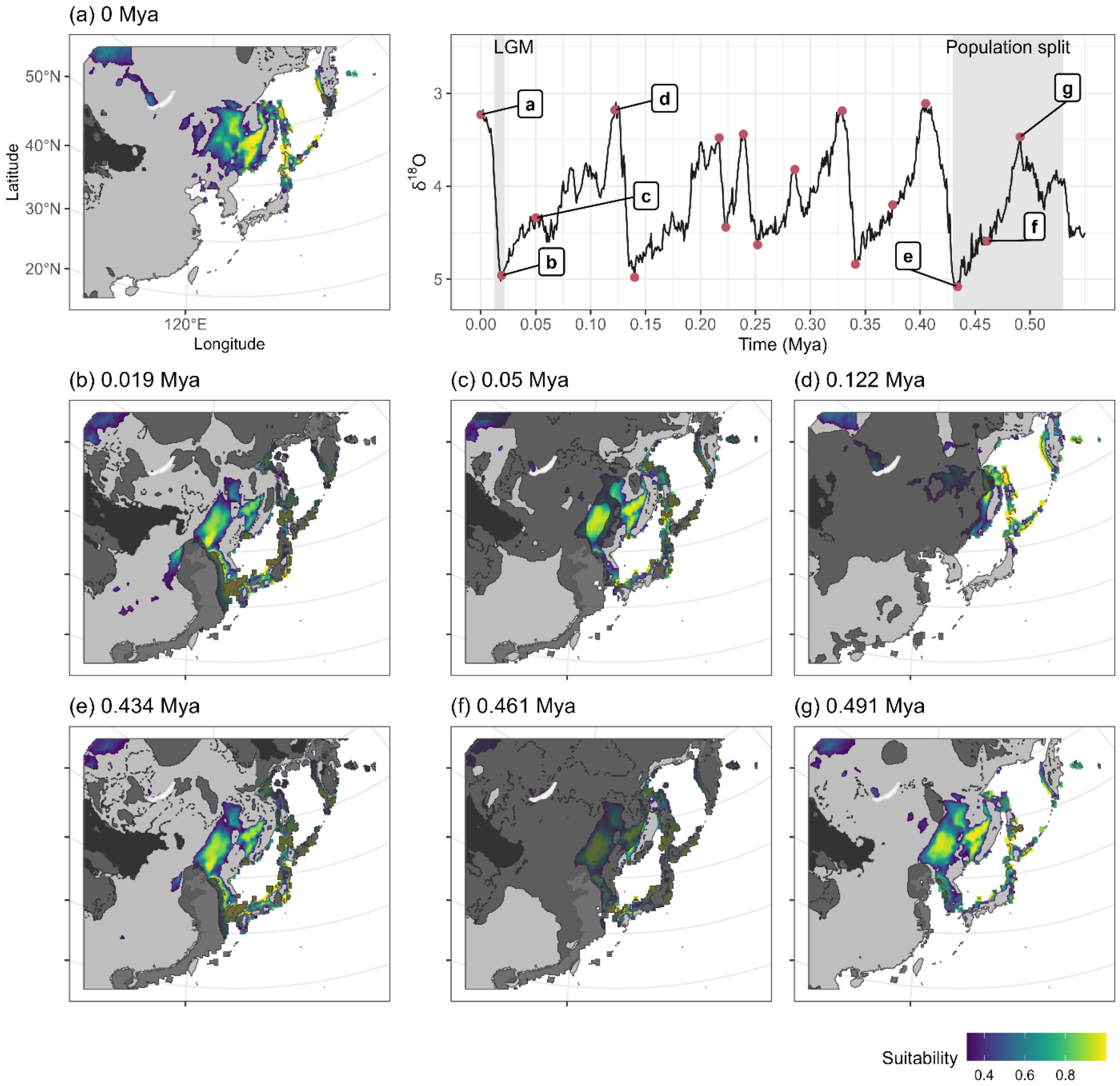


**Figure S22** Past climate projection of the species distribution model for the breeding range of the Swinhoe's Rail (a-g). On the top right panel, a proxy for paleoclimatic temperature ($\delta^{18}$O) (Lisiecki and Raymo 2005; Spratt and Lisiecki 2016) was plotted, and periods that corresponded to each time slice (a-g) were indicated on it. Higher $\delta^{18}$O values indicate higher temperatures. The time period (a), (b)-(d), and (e)-(g) correspond to the present, the last glacial period when asymmetric gene flow was estimated in the FSC2, and the time of population split inferred by multiple results. Color gradients from indigo to yellow represent environmental suitability for rails. Black arrows in the projection to the present climate layer (a) indicate the presently or previously known distribution areas of this species. Regions with dark gray shades and colors overlaid are where model extrapolation was restricted due to the absence of the climate in the modeling domain (i.e., grids with negative MESS values, see Main Text) and regions with ice sheets or deserts, respectively. Lambert Azimuthal Equal Area projection was used for plotting.

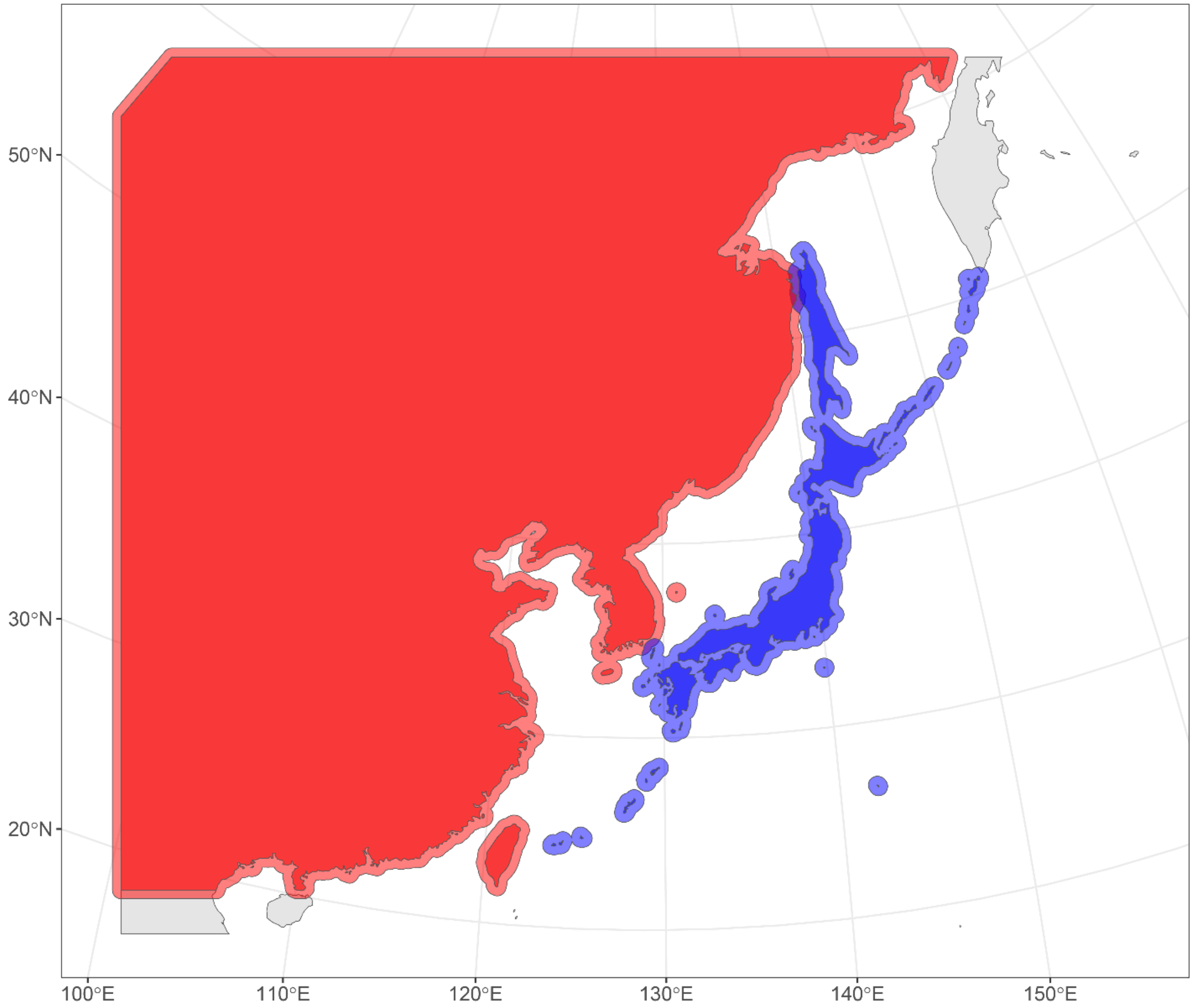


**Figure S23** The definitions of the continent and island to compare historical changes in the habitat between the two regions. Blue and red indicate the island and continent, respectively. Overlapping regions are where values were included in both regions.

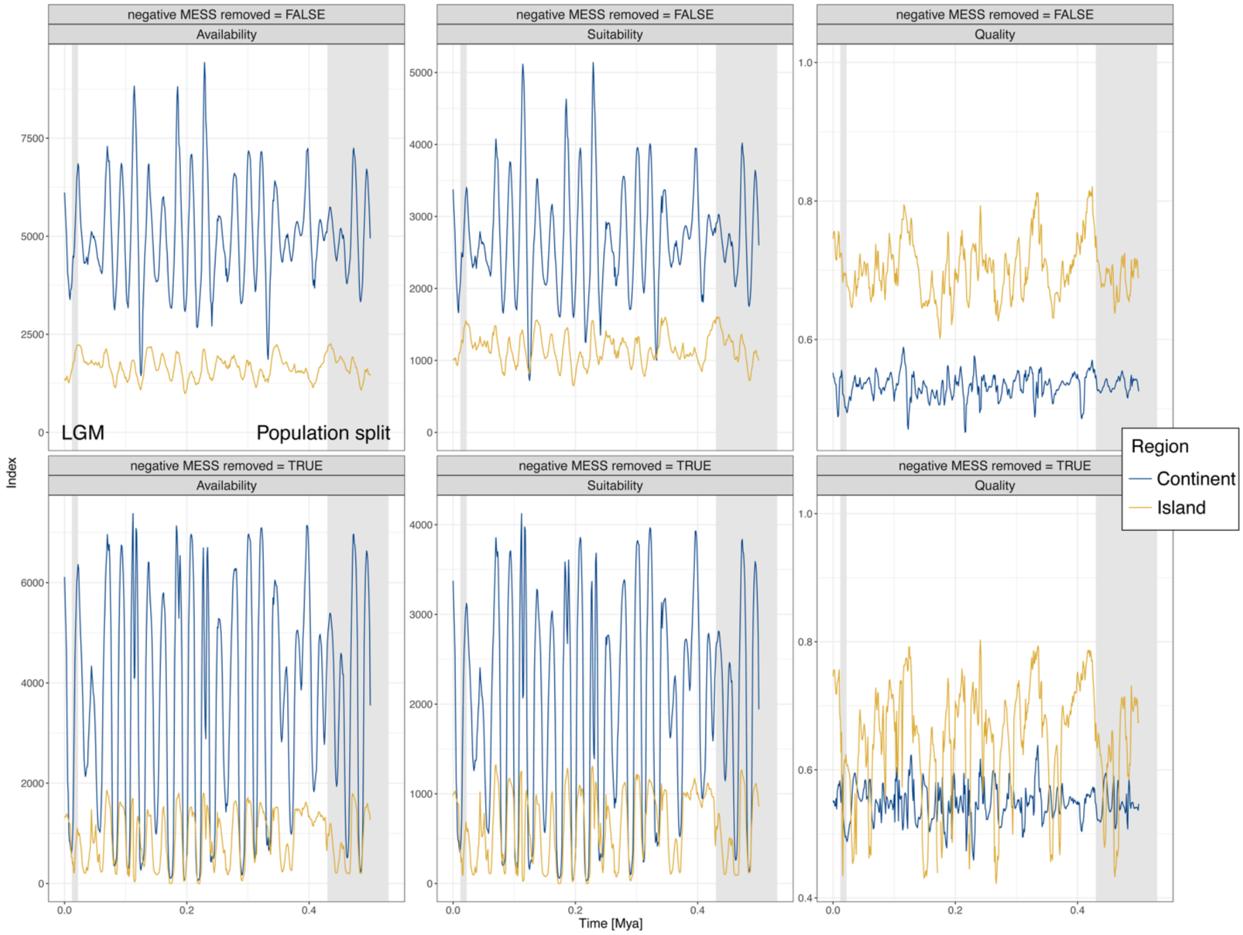


**Figure S24** Historical changes of habitat availability, suitability, and quality for the predicted presence grids on the continent (blue) and island (yellow). These values were calculated using predicted grids regardless of their values of MESS (top three panels) and using grids that were not covered by negative MESS values (bottom three panels). Availability is the total number of presence grids, suitability is the sum of the values of presence grids, and quality is suitability/availability. The two gray bars indicate the estimated times of population split (‘Population split’ around 0.5 Mya) and gene flow (‘LGM’) in our demographic modeling.

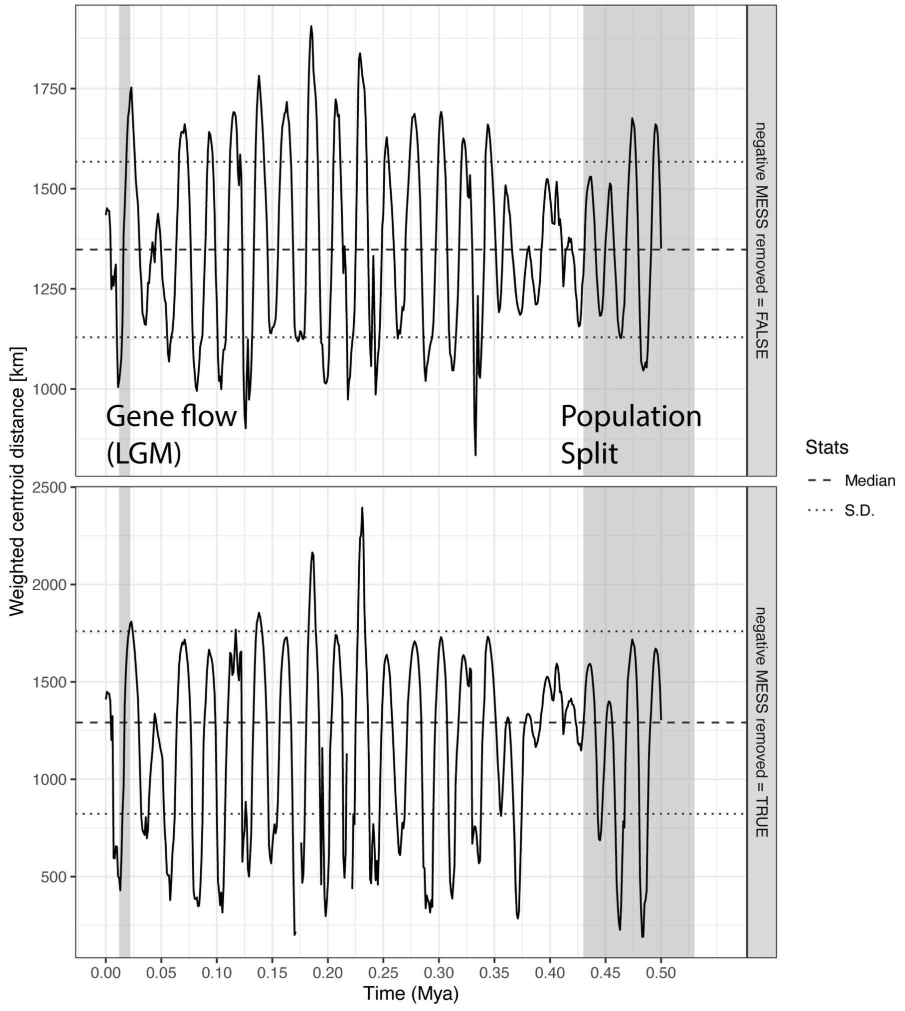


**Figure S25** Historical changes of centroid distance between island and continental habitat, calculated using predicted grids regardless of their values of MESS (top three panels) and using grids that were not covered by negative MESS values (bottom three panels). Refer to Figure 4 for other annotations.

Fukuda, A., Y. Odaya, K. Shirakawa, H. Shirakawa, S. Ozawa, M. Imai, M. Fukagawa, Y.

Umegaki, and Y. Yamaguchi. 2019. Swinhoe's Rail regularly overwinters in Ibaraki Prefecture, Japan. Bird Res. 15:A1–A14.

Hasegawa, M., H. Kishino, and T. Yano. 1985. Dating of the human-ape splitting by a molecular clock of mitochondrial DNA. J. Mol. Evol. 22:160–174.

Heim, W., D. Trense, A. Heim, and J. Kamp. 2019. Discovery of a new breeding population of the vulnerable Swinhoe's Rail *Coturnicops exquisitus* confirmed by genetic analysis. Bird Conservation International 29:454–462.

Hijmans, R. J., R. Bivand, K. Forner, J, Ooms, E. Pebesma, and M. D. Sumner. 2022. Package 'terra'. Maintainer: Vienna, Austria, 384.

Huson, D. H., and D. Bryant. 2006. Application of phylogenetic networks in evolutionary studies. Molecular Biology and Evolution, 23:254–267.

Kass, J. M., R. Muscarella, P. J. Galante, C. L. Bohl, G. E. Pinilla-Buitrago, R. A. Boria, M. Soley-Guardia, and R. P. Anderson. 2021. ENMeval 2.0: Redesigned for customizable and reproducible modeling of species' niches and distributions. Methods Ecol. Evol. 12:1602–1608.

Kitazawa, M., Y. Yamaura, M. Senzaki, M. Hanioka, H. Ohashi, M. Oguro, T. Matsui, and F. Nakamura. 2022. Quantifying the impacts of 166 years of land cover change on lowland bird communities. Proc. Biol. Sci. 289:20220338.

Kuznetsova, A., P. B. Brockhoff, and R. H. B. Christensen. 2017. lmerTest Package: tests in linear mixed effects models. J. Stat. Softw. 82:1–26.

Lawson, D. J., L. van Dorp, and D. Falush. 2018. A tutorial on how not to over-interpret STRUCTURE and ADMIXTURE bar plots. Nat. Commun. 9:3258.

Leaché, A. D., B. L. Banbury, J. Felsenstein, A. N.-M. de Oca, and A. Stamatakis. 2015. Short tree, long tree, right tree, wrong tree: new acquisition bias corrections for inferring SNP phylogenies. Syst. Biol. 64:1032–1047.

Li, H. 2013. Aligning sequence reads, clone sequences and assembly contigs with BWA-MEM.

Li, H. 2011. Improving SNP discovery by base alignment quality. Bioinformatics 27:1157–1158.

Lou, R. N., A. Jacobs, A. P. Wilder, and N. O. Therkildsen. 2021. A beginner's guide to low-coverage whole genome sequencing for population genomics. Mol. Ecol. 30:5966–5933.

Naimi, B., N. A. S. Hamm, T. A. Groen, A. K. Skidmore, and A. G. Toxopeus. 2014. Where is potential uncertainty a problem for species distribution modelling? Ecography 37:191–203.

Matsumiya, H., and M. Numano. 2022. Swinhoe's Rail *Coturnicops exquisitus* wintering on the Pacific side of Chubu and Kinki region, Japan. Bird Res. 18:A87–A97.

Mlíkovský, J., D. G. D. Russell, and P. Stýblo. 2024. Breeding distribution and seasonal movements of Swinhoe's Rail *Coturnicops exquisitus* (Aves: Rallidae). J. Natl. Mus., Nat. Hist. Ser. 193:103–120.

Ozaki, K., Y. Yamamoto, and S. Yamagishi. 2010. Genetic diversity and phylogeny of the endangered Okinawa Rail, Gallirallus okinawae. Genes Genet. Syst. 85:55–63.

Pickrell, J. K., and J. K. Pritchard. 2012. Inference of population splits and mixtures from genome-wide allele frequency data. PLoS Genet. 8:e1002967.

Polzin, T., and S. V. Daneshmand. 2003. On Steiner trees and minimum spanning trees in hypergraphs. Oper. Res. Lett. 31:12–20.

Remington, D. L., J. M. Thornsberry, Y. Matsuoka, L. M. Wilson, S. R. Whitt, J. Doebley, S. Kresovich, M. M. Goodman, and E. S. Buckler 4th. 2001. Structure of linkage disequilibrium and phenotypic associations in the maize genome. Proc. Natl. Acad. Sci. 98:11479–11484.

Reynolds, J., B. S. Weir, and C. C. Cockerham. 1983. Estimation of the coancestry coefficient: basis for a short-term genetic distance. Genetics 105:767–779.

Sadakuni, T., M. Takahashi, A. Miya, K. Isogai, K. Matsuoka, M. Senzaki, M. Kitazawa, and O. Harada. 2018. Status of the summer occurrence of Swinhoe's Rail in Hokkaido. Proceedings of The Ornithological Society of Japan Annual Meeting 2018, P037. [in Japanese]

Schliep, K. P. 2011. phangorn: phylogenetic analysis in R. Bioinformatics 27:592–593.

Sorenson, M. D., J. C. Ast, D. E. Dimcheff, T. Yuri, and D. P. Mindell. 1999. Primers for a PCR-based approach to mitochondrial genome sequencing in birds and other vertebrates. Mol. Phylogenet. Evol. 12:105–114.

Stange, M., M. R. Sánchez-Villagra, W. Salzburger, and M. Matschiner. 2018. Bayesian divergence-time estimation with genome-wide single-nucleotide polymorphism data of sea catfishes (Ariidae) supports Miocene closure of the Panamanian Isthmus. Syst. Biol. 67:681–699.

Suyama, Y., and Y. Matsuki. 2015. MIG-seq: an effective PCR-based method for genome-wide single-nucleotide polymorphism genotyping using the next-generation sequencing platform. Sci. Rep. 5:16963.

Takahashi, M., J. Ebina, A. Miya, K. Isogai, T. Furuyama, T. Takeda, M. Horikoshi, C. Ohe, and T. Kanouchi. 2018. Swinhoe's Rail *Coturnicops exquisitus* wintering in the Kanto region of Japan. Jpn. J. Ornithol. 67:109–116.

Tamura, K., G. Stecher, and S. Kumar. 2021. MEGA11: Molecular Evolutionary Genetics Analysis Version 11. Mol. Biol. Evol. 38:3022–3027.

van der Bank, F. H., G. D. Engelbrecht, H. Sauer-Gürth, M. Wink, and P. F. Mulder. 1998.

Allozyme and DNA sequence data support speciation of northern and southern populations of silver catfish, *Schilbe intermedius* (Rüppel, 1832). Comp. Biochem. Physiol. A Mol. Integr. Physiol. 120:531–543.

Vasimuddin, M., S. Misra, H. Li, and S. Aluru. 2019. Efficient architecture-aware acceleration of BWA-MEM for multicore systems. Pp. 314–324 *in* 2019 IEEE international parallel and distributed processing symposium (IPDPS). IEEE.

Vieira, F. G., F. Lassalle, T. S. Korneliussen, and M. Fumagalli. 2015. Improving the estimation of genetic distances from Next-Generation Sequencing data. Biol. J. Linnean Soc. 117:139–149.

Yu, G., D. K. Smith, H. Zhu, Y. Guan, and T. T.-Y. Lam. 2017. Ggtree: An r package for visualization and annotation of phylogenetic trees with their covariates and other associated data. Methods Ecol. Evol. 8:28–36.
